# Can we assume the gene expression profile as a proxy for signaling network activity?

**DOI:** 10.1101/643866

**Authors:** Mehran Piran, Reza Karbalaei, Mehrdad Piran, Jehad Aldahdooh, Mehdi Mirzaie, Naser Ansari-Pour, Jing Tang, Mohieddin Jafari

## Abstract

Studying relationships among gene-products by gene expression profile analysis is a common approach in systems biology. Many studies have generalized the outcomes to the different levels of central dogma information flow and assumed correlation of transcript and protein expression levels. All these efforts partook in the current understanding of signaling network models and expanded the signaling databases. In fact, due to the unavailability or high-cost of the experiments, most of the studies do not usually look for direct interactions, and some parts of these networks are contradictory. Besides, it is now a standard step to accomplish enrichment analysis on biological annotations, to make claims about the potentially implicated biological pathways in any perturbation. Explicitly, upon identifying differentially expressed genes, they are spontaneously presumed the corresponding dysregulated pathways. Then, molecular mechanistic insights are proposed for disease etiology and drug discovery based on statistically enriched biological processes. In this study, using four common and comprehensive databases, we extracted all relevant gene expression data and all relationships among directly linked gene pairs. We aimed to evaluate the ratio of coherency or sign consistency between the expression level and the causal relationships among the gene pairs. We illustrated that the signaling network was not more consistent or coherent with the recorded expression profile compared to the random relationships. Finally, we provided the pieces of evidence and concluded that gene-product expression data, especially at the transcript level, are not reliable or at least insufficient to infer causal biological relationships among genes and in turn, describe cellular behavior.

## 1. Introduction

In network biology, defining causal relationships among nodes is crucial for the static and dynamic analysis (1, 2). The most available high-throughput data to infer molecular relationships are arguably whole-transcriptome expression profiles analyzed with statistical models (3). The challenge is extrapolating causality in signaling and regulatory mechanisms from a significant correlation between any given gene pair. Lots of spurious correlations among gene pairs may occur without any causal relationship that could happen indirectly or stochastically (4). So far, reverse engineering algorithms are developed to tackle this challenge and to infer gene networks and regulatory interactions from expression profiles (5–7).

When considering signaling networks, as a portrait of causal relationships among the molecular entities in biology, their leading players are proteins whose activity is often regulated by post-translational modifications such as phosphorylation. Hence, inference of signaling networks can be directly inferred from (Phospho) proteomic and protein-protein interaction data (8). However, these kinds of data are cost-consuming and tough processing to acquire. Given the correlation between protein and gene expression, a common alternative approach is to use gene expression to estimate interactions between proteins. We know that the gene expression or transcriptome talk about *“what appears to happen in a biological system”*, while the signaling network exhaust to *“what makes it happens and has happened in a complex view of the system”* (9). This, therefore, begs the question of whether gene expression profiles strengthen the logic mechanism of signaling circuits, i.e. activatory/inhibitory relationships.

In this study, we aimed to examine the coherency between expression profiles and the signs of relationship, in signaling networks, for all possible gene pairs. Imagine in a gene pair (I, II) where gene I activates gene II. If the expression profiles of both were correlated positively; we infer that expression data strengthen the logic of this signaling relationship and are thus coherent In contrast, let gene I inhibits gene II. In this case, the coherent gene pairs are negatively correlated. If gene I activates gene II and there is a negative correlation between them or if gene I inhibits gene II and there is a positive correlation between them, this implies the incoherency between the gene pairs relationship. In addition to these simple scenarios, we have also considered more complicated subgraphs in a signaling network to answer the question raised above (See Table 1).

**Table 1:**
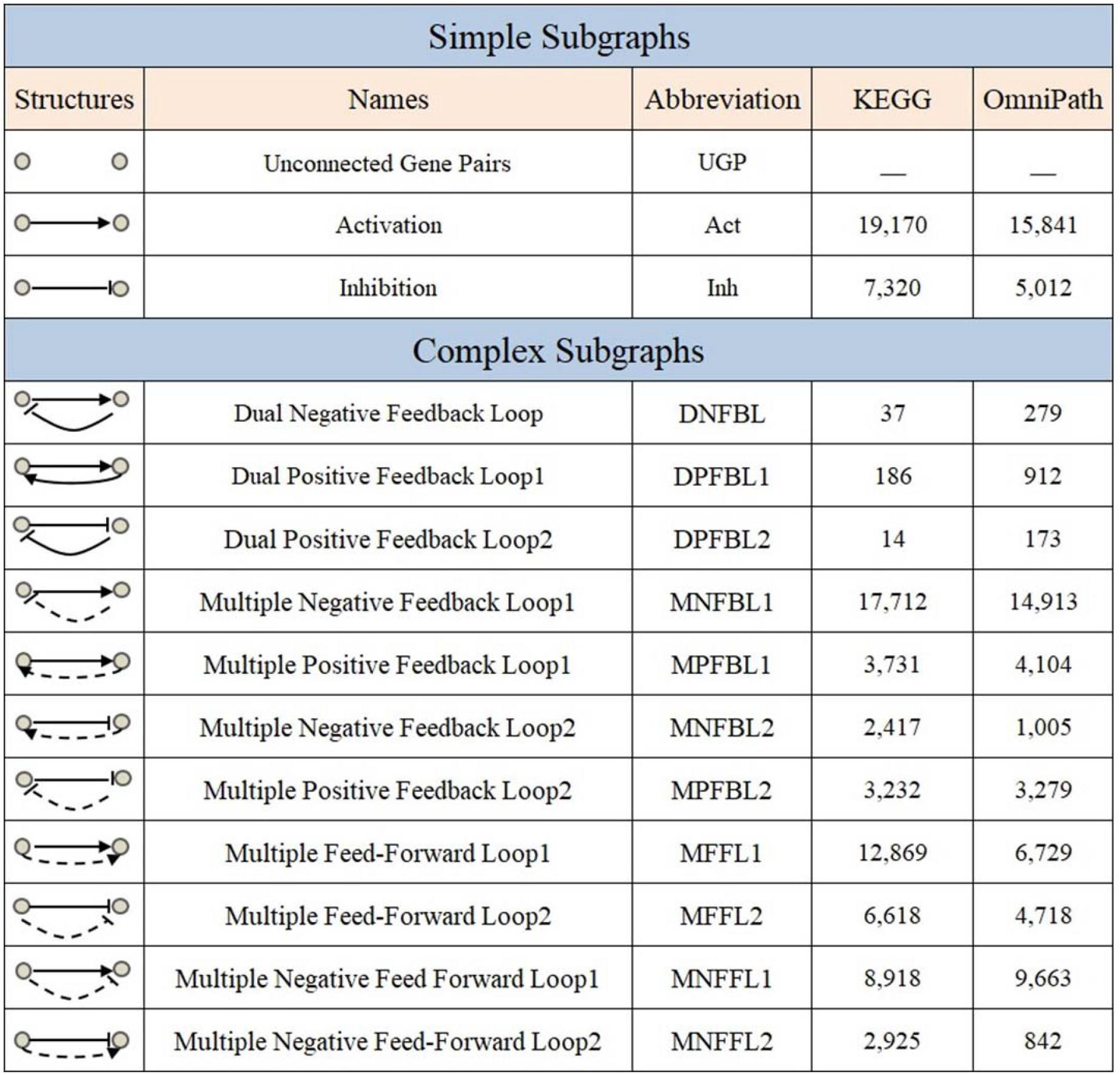
Details of different subgraphs present in ail biological signaling networks. The dashed lines indicate multiple edges between nodes. The last two columns provide the number of each subgraph in the two signaling databases.

| Simple Subgraphs |  |  |  |  |
| --- | --- | --- | --- | --- |
| Structures | Names | Abbreviation | KEGG | OmniPath |
|  | Unconnected Gene Pairs | UGP | — | — |
|  | Activation | Act | 19,170 | 15,841 |
|  | Inhibition | Inh | 7,320 | 5,012 |
| Complex Subgraphs |  |  |  |  |
|  | Dual Negative Feedback Loop | DNFBL | 37 | 279 |
|  | Dual Positive Feedback Loop1 | DPFBL1 | 186 | 912 |
|  | Dual Positive Feedback Loop2 | DPFBL2 | 14 | 173 |
|  | Multiple Negative Feedback Loop1 | MNFBL1 | 17,712 | 14,913 |
|  | Multiple Positive Feedback Loop1 | MPFBL1 | 3,731 | 4,104 |
|  | Multiple Negative Feedback Loop2 | MNFBL2 | 2,417 | 1,005 |
|  | Multiple Positive Feedback Loop2 | MPFBL2 | 3,232 | 3,279 |
|  | Multiple Feed-Forward Loop1 | MFFL1 | 12,869 | 6,729 |
|  | Multiple Feed-Forward Loop2 | MFFL2 | 6,618 | 4,718 |
|  | Multiple Negative Feed Forward Loop1 | MNFFL1 | 8,918 | 9,663 |
|  | Multiple Negative Feed-Forward Loop2 | MNFFL2 | 2,925 | 842 |

To reach this end, we used expression datasets in the Gene Expression Omnibus (GEO) (10) and Genomics of Drug Sensitivity in Cancer (GDSC) (11) to extract the relevant gene expression profiles. Two literature-curated databases for signaling pathways, namely the Kyoto Encyclopedia of Genes and Genomes (KEGG) (12) and OmniPath (13) (which integrates literature-curated human signaling pathway from 34 resources) were used to extract the sign of relationships among directly linked gene pairs. Therefore, coherency analysis was undertaken independently for all four combinations of databases in parallel (see Figure 1).

**Figure 1:**
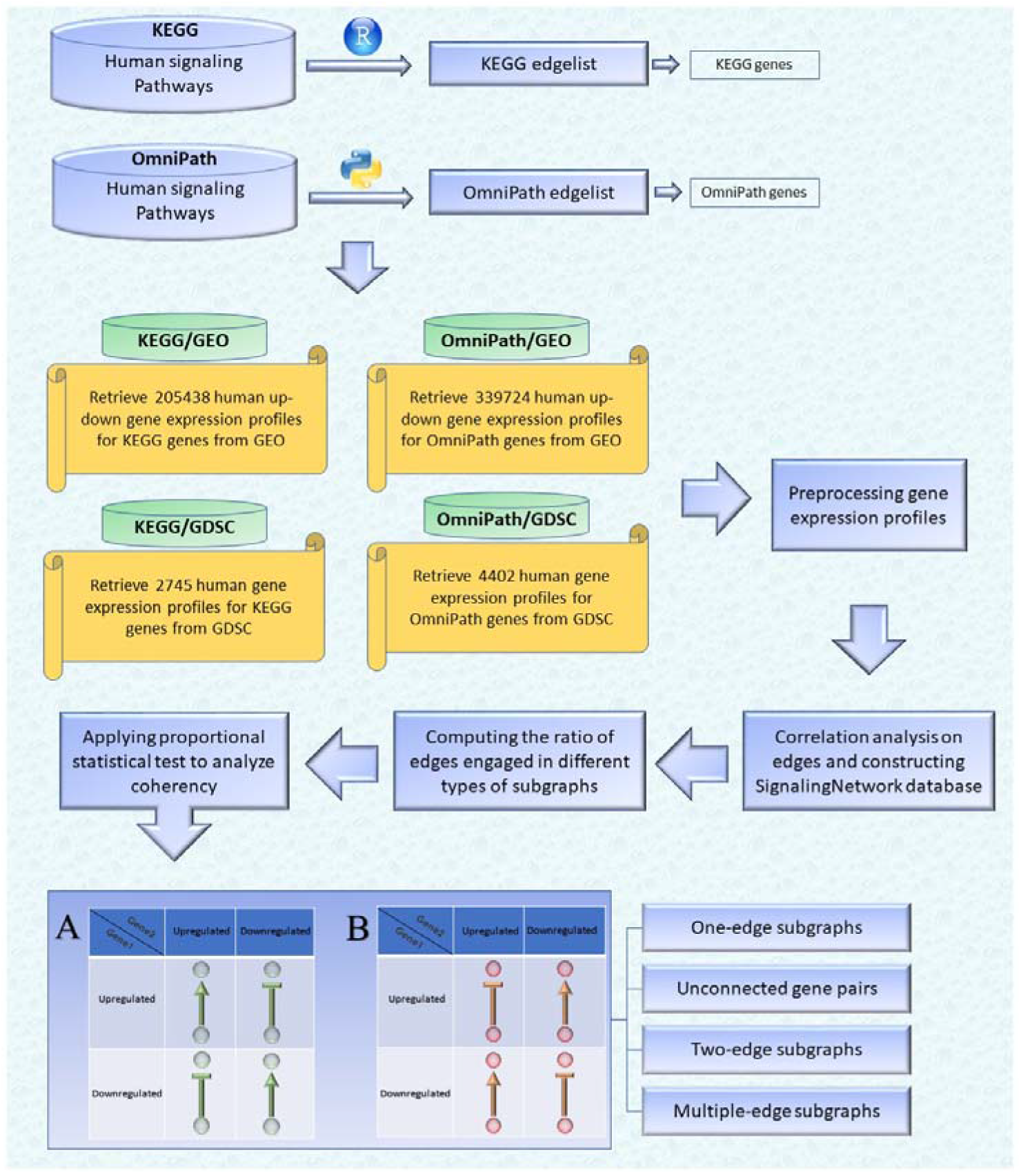
Visual overview of how information from different databases was integrated to analyze the coherency. An edge list was constructed from KEGG and OmniPath databases. Ail the gene expression profiles for the edge list genes were then downloaded from GEO and GDSC databases. Next, data were preprocessed and a suitable structure was created for correlation analysis among the gene pairs. By interpreting the information from correlation tests and the proportional tests, coherency analysis was implemented on different forms of subgraphs. There is a total of four coherent conditions in panel A and four incoherent conditions in panel B. For instance, in panel A, ifgenel is up-regulated and there is an activation between the gene pair, gene2 must be upregulated. In panel B, ifgenel is up-regulated and there is an inhibitory relationship between the gene pair, gene2 is expected to be up-regulated.

## 2. Materials and Methods

In this study, four independent analyses were performed based on two gene expression databases i.e., GEO and GDSC and two signaling pathway databases i.e., KEGG and OmniPath in parallel (Figure 1) (10–13). Thus, the signaling pathway databases were independently used to reconstruct a whole signaling network, and the gene expression databases were separately used to apply correlation analysis on each gene pair in the pathways to do and compare KEGG/GEO, KEGG/GDSC, OmniPath/GEO, and OmniPath/ GDSC distinct analyses and findings. To briefly introduce the used gene expression databases, GEO is an NCBI international public repository that archives microarray and next-generation sequencing expression data. The GDSC database is the largest public repository that archives information about drug sensitivity in cancer cells and biomarkers of drug response in these cells. In this work, gene expression profiles from GDSC cell lines and GEO studies were used to extract pairwise association between genes.

### 2.1 Signaling network reconstruction

Here, we focused on human signaling pathways based on available datasets. All human-related signaling pathways were downloaded from the KEGG database. Using the *KEGGgraph* package (14), these pathways were imported into the R environment (15). Edge information was extracted, and each graph was converted to an edge list Next, all edges (n=26490) were merged, and a directed signed signaling network was reconstructed (Supplementary file 1, section 1 and Supplementary file 4). Eligible edges (see section 2.3) were then selected, and correlation analysis was undertaken on eligible gene pairs. The *pypath* python module (13) was also used to do the same and create an edge list based on the OmniPath database (see Supplementary file 4). This edge list (n=20853) was also imported into the R environment for the downstream statistical analysis on the gene pairs.

### 2.2 Gene expression profiles extraction

The standard GEO query format (GEO Profiles) were used to identify all up- and down-expressed or differentially expressed genes (DEG) representing within the KEGG and/or OmniPath edge lists. Gene expression profiles available in GDSC were downloaded for both edge lists, followed by preprocessing and outlier detection. Finally, based on GEO and GDCS, four expression matrices were created using the genes which make up of KEGG and OmniPath edge list (Supplementary file1 sections 2 and 3. Supplementary files 5). For more details about the total number of gene expression profiles, DEGs, and total number of samples which were downloaded from GEO and GDSC, see Figure 1 and Table 2.

**Table 2:** General properties and date retrieved of the signaling networks. The number of DEGs are also given, which are those commons between the edge list genes and gene expression profile genes and identified by the GEO/G DSC database either up- or down-regulated. Samples are total number of samples in GEO and GDSC databases for which expression data were available for the given gene pair. The node number of the giant component, the diameter of the network and the ratio of shared genes between edge list genes and gene-expression-profile genes are presented in the last three columns, respectively.

| KEGG |  |  |  |  |  |  | OmniPath |  |  |  |  |  |  |
| --- | --- | --- | --- | --- | --- | --- | --- | --- | --- | --- | --- | --- | --- |
|  | Date Retrieved | DEGs | Samples | Giant Component | Diameter | Ratio |  | Date Retrieved | DEGs | Samples | Giant Component | Diameter | Ratio |
| GEO | 2017.08 | 3047 | 40903 | 2549 | 17 | 0.95 |  | 2019.05 | 4724 | 40774 | 3848 | 17 | 0.95 |
| GDSC | 2017.10. | 2745 | 1018 | 2583 | 17 | 0.16 |  | 2019.05 | 4402 | 1018 | 4045 | 15 | 0.25 |

### 2.3 Mutual association analysis

In the next step, correlation on the expression profiles of each gene pair were statistically tested. For computing correlation coefficients for any gene pair, including Pearson, Spearman’s rank, and Kendall rank coefficients, we only considered gene pairs having more than two samples. These gene pairs were considered as eligible edges for downstream statistical analysis (Supplementary file 1, section 4). Samples with expression data for the gene pairs may have come from different datasets with multiple organism and tissues, therefore should be separated and analyzed independently. Figure 2A represents the effect of this preprocessing on an exemplary gene pair in our dataset. In this study, the gene expression profiles were considered dataset-specific to avoid any inconsistency among the samples collected from diverse datasets. In other words, sample heterogeneity can easily affect any pairwise relationship. An edge is therefore considered as homogeneous if the correlation sign is consistent across all. These homogeneous edges were used for correlation analysis. Then, according to the statistical significance and the sign of the correlation coefficient, the coherent and incoherent gene pairs were inferred (Supplementary file 1, sections 5). Note that the p-value less than 0.05 was considered to represent as cutoff in Figure 4 and 5.

**Figure 2:**
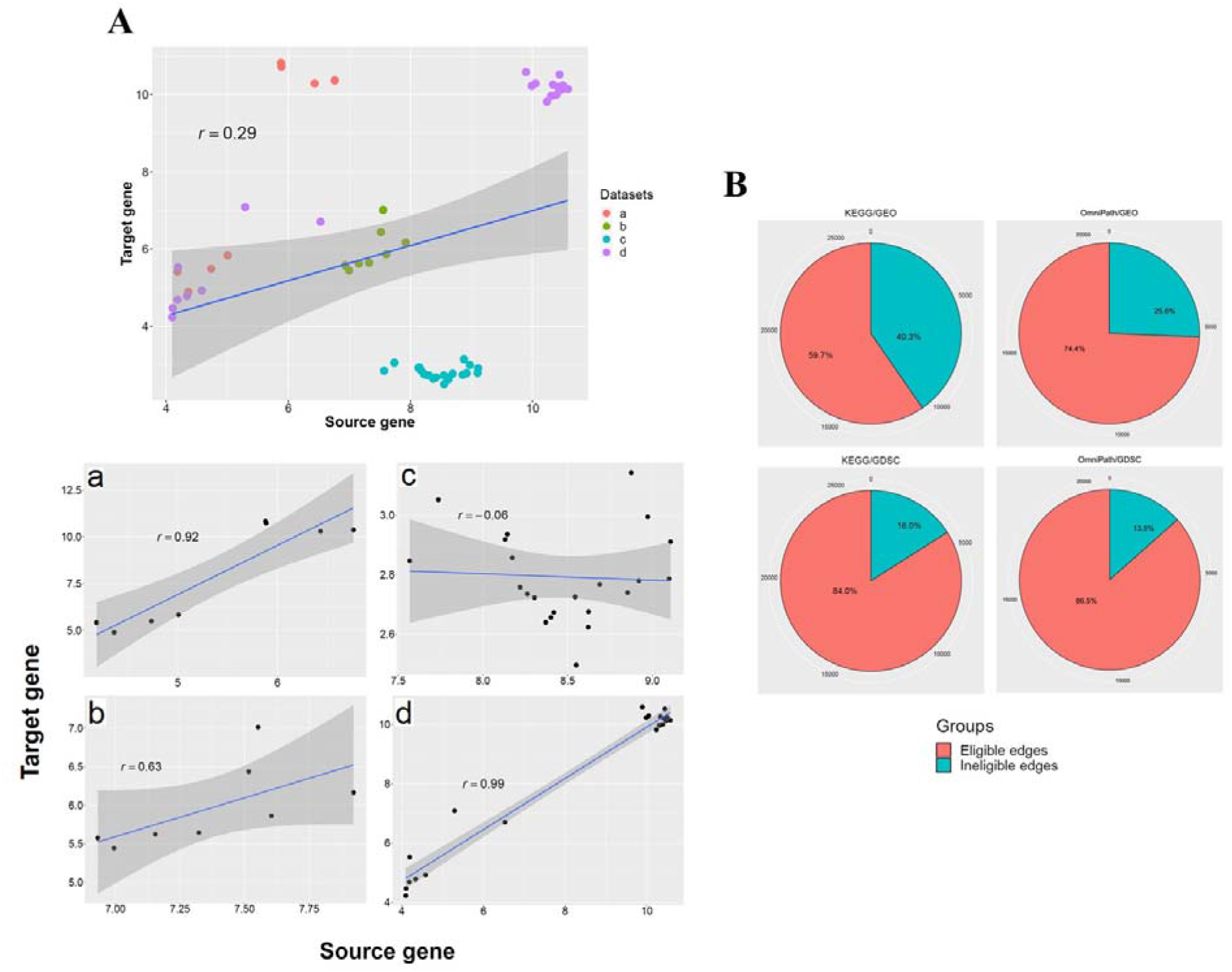
(A) An exemplary relationship between gene pair expression. These scatter plots contain the Pearson coefficient correlations and fitted linear regression line. The X-axis and Y-axis values differ according to the expression profile of this gene pair in different gene expression dataset. It is depicted the gene expression profiles of this exemplary gene pairs in the edge list before pre-processing. The same gene pair’s expression profiles separated to the four relevant datasets. (B) The proportion of eligible and ineligible edges in the four parallel analyses. The numbers around each chart represent the number of edges at that point.

### 2.4 Randomly selected unconnected gene pairs

The edge lists obtained in the previous step were converted into adjacency matrices using igraph package in R (16). Then, the adjacency matrix was self-multiplied more than (e.g., n>17) the diameter of the network (Table 2), After that, we randomly selected 1000 unconnected gene pairs ten times for which the corresponding elements in the matrix were zero (gene pairs with no direct immediate and non-immediate interactions). For these gene pairs, that we called unconnected gene pairs (UGPs), the same downstream analyses, i.e., pre-processing and correlation analysis, were implemented to compare significance and sign of correlation coefficients to connected gene pairs (Supplementary file 1, section 6, and Supplementary files 7).

### 2.5 Complex subgraphs

Since gene pairs are not isolated within the whole signaling networks, the larger subnetworks, consisted of gene pairs relationships, also need to be considered. So, we extracted specific subgraphs from the signaling networks to investigate any association between gene expression profiles and the complex structures of gene pairs relationships. DNFBL, DPFBL1, and DPFBL2 are subgraphs of gene pairs which influence each other directly twice (see Table 1 for the full names of subgraphs). These pairs are readily found by checking the source and target nodes in the edge lists (or upper and lower triangles in adjacency matrices). We then focused on the connected gene pairs, which also influence each other indirectly by a sequence of intermediate nodes. Following matrix self-multiplication, the weighted and unweighted adjacency matrices of the giant component of eligible edges in signaling network were powered by the network radius magnitude. Considering that the network is directed and the adjacency matrix is not symmetric, the feed-forward and feedback loops i.e., MNFBL1-2, MPFBL1-2, MFFL1-2, and MNFFL1-2 are determined (Table 1). For a more detailed explanation, see Supplementary File 1, sections 7 and 8.

## 3. Results

The overall details of the four parallel coherency analyses were presented including the dimension of the expression matrices generated from whole-transcriptome expression profiles, and the size and diameter of the giant component in each analysis (Table 2). Of note, the number of DEGs was higher in OmniPath than KEGG even though the size of KEGG network is 1.8-fold of the OmniPath network. |The ratio of eligible edges to all edges was calculated for all four analyses (see Figure 2B). The ratio of eligible edges in the OmniPath edge list also was higher than KEGG based on both GDSC and GEO databases. In addition, the ratio of eligible edges was higher in GDSC compared with GEO. These comparison indicates the higher quality of gathered and annotated data in OmniPath and GDSC databases.

### 3.1 The ratio of coherency for gene pairs

After filtering out heterogeneous edges, an extensive list of homogeneous edges was constructed (Supplementary file 1 sections 3.5 - 3.7 and Supplementary files 6) for correlation analysis. The violin plots of Pearson correlation coefficients for each analysis are shown in Figure 3A. The distribution of the coefficients shows a nearly uniform distribution with a little left skewness for KEGG/GEO and OmniPath/GEO while for KEGG/GDSC and OmniPath/GDSC, it follows a normal distribution with the median at approximately zero. In addition to the issue of different sample size in GEO and GDSC, this suggests that for GDSC-based edges, correlations between the expression profiles of the gene pairs do not tend to show a high positive or negative correlation. In other words, for a given gene pair (I, II), over-expression or under-expression of gene I does not have a substantial effect on the expression of gene II regardless of the edge sign.

**Figure 3:**
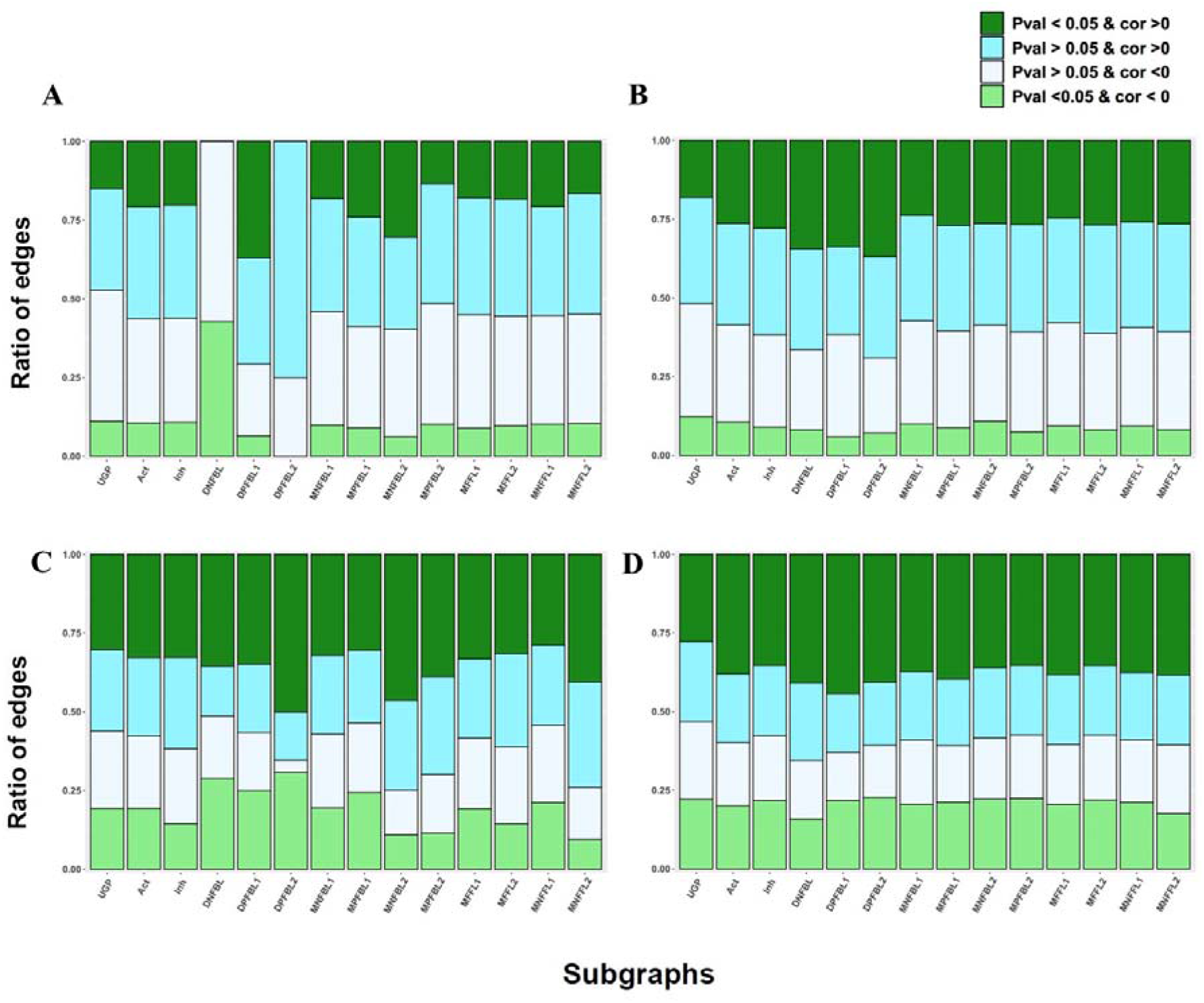
(A) Distribution of Pearson correlation-coefficient values for the four parallel coherency analyses. (B) The ratio of coherent, incoherent and NA gene pairs. The values around each pie chart represent the exact numbers.

**Figure 4:**
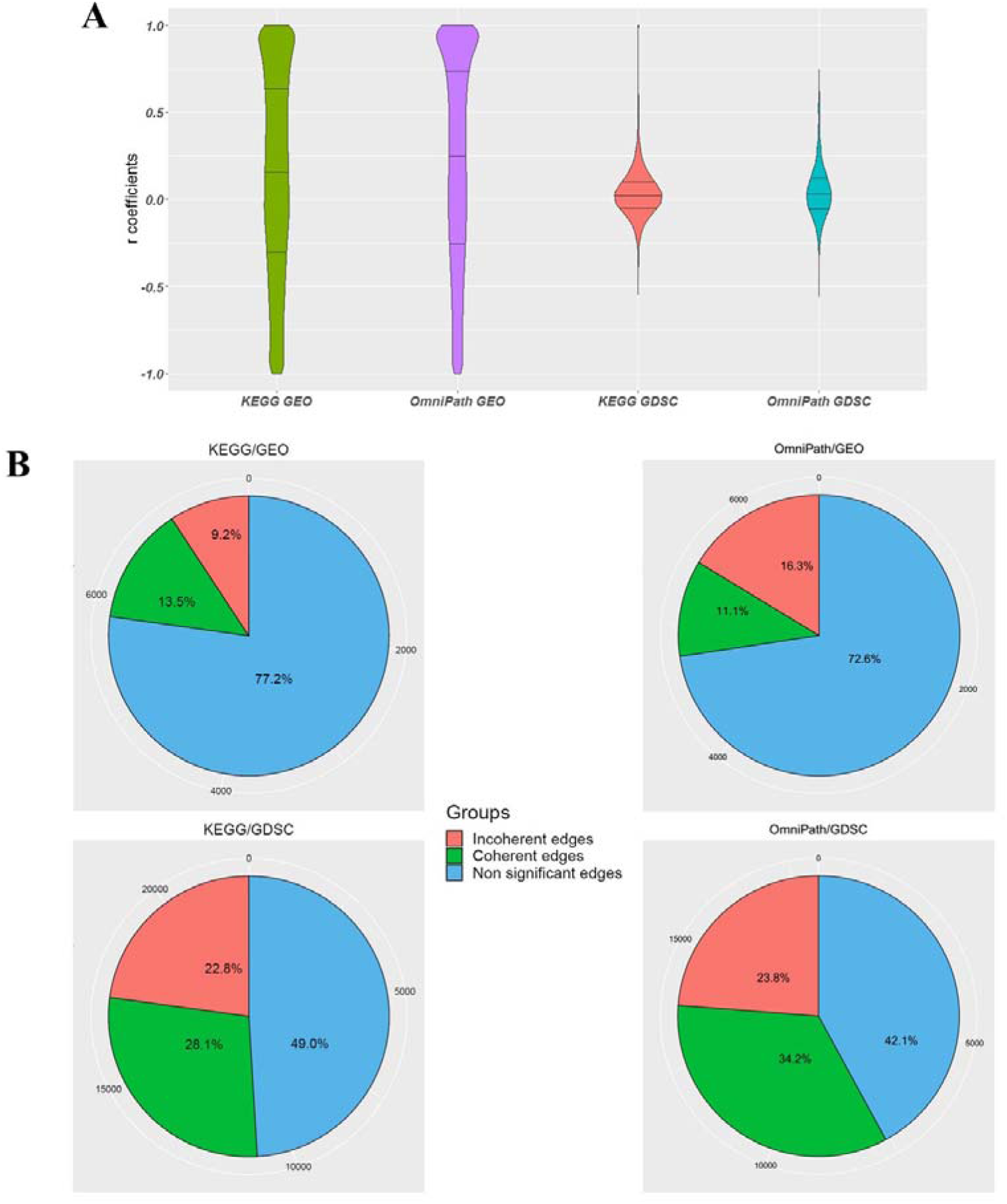
The volcano plots of the activation and inhibition edges. The horizontal axis is the Pearson correlation coefficient, and the vertical axis shows log transformed FDR-adjusted p-values. The threshold line (bluej represents the significance cut-off value of 0.05. (A), (B), (C) and (D) plots correspond to KEGG/GEO, OmniPath/GEO, KEGG/GDSC, and OmniPath/GDSC analyses, respectively.

**Figure 5:**
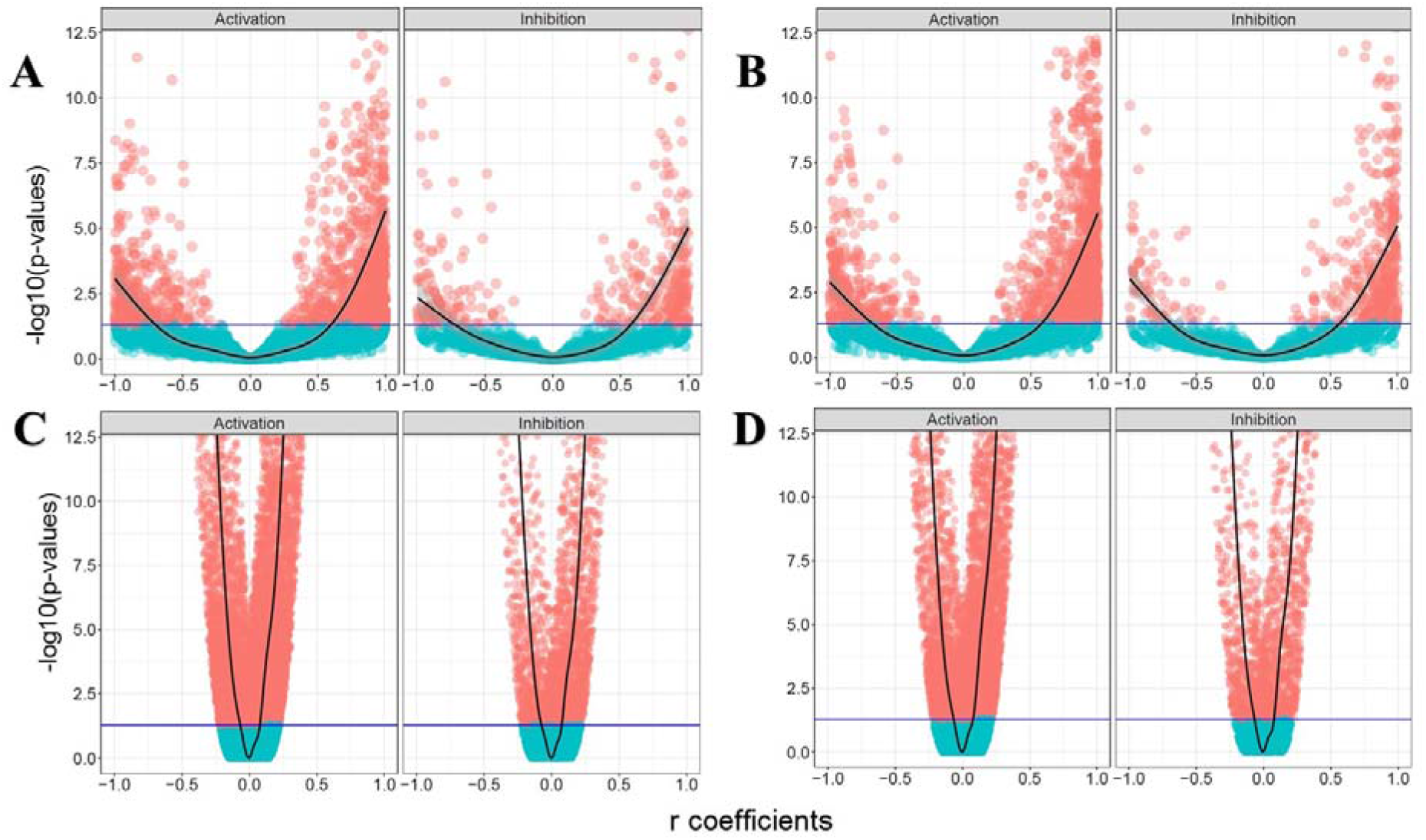
The ratio of eligible and homogeneous edges involved in different subgraphs are represented by stacker bar plots for all four analyses. (A) and (B) are KEGG/GEO and OmniPath/GEO plots, and (C) and (D) plots correspond to KEGG/GDSC and OmniPath/GDSC. Note that the UGP ratios are the average ratio over of all UGP set. See Table 1 for the full names of subgraphs.

Figure 3B depicts the ratios of coherent and incoherent gene pairs along with the number of non-significant edges which have the FDR-adjusted p-values larger than 0.05, and we could not declare about the coherency status by a likelihood greater than or equal to 95%. In addition, the sum of the ratio of incoherent gene pairs and nonsignificant edges were more than the ratio of coherent gene pairs in all four analyses. The ratio of coherent gene pairs in OmniPath is in general more than KEGG. Also, the ratio of coherent gene pairs in GDSC database is more than GEO. In the following, we represented the results based on the Pearson correlation coefficient However, note that the ratio of coherency for gene pairs is almost similar using Spearman and Kendall rank correlation coefficients too (Supplementary file 1, section 4),

Figure 4 shows the FDR-adjusted p-values versus correlation coefficients of activation and inhibition edges in all four analysis. The symmetric pattern of coefficients is recognizable for both activation and inhibition edges in the four analyses. It suggests that the correlation between a given gene pair is not predominantly affected by the sign of the interaction. In other words, although activation edges illustrate an overrepresentation of strong positively correlated gene pairs in all four analysis, the inhibition edges do not display any enrichment in the strong negative side of plots compared to the strong positive side. It also shows that the majority of coherent gene pairs are related to activation, not inhibition edges. In the next step, we tried to explore and provide reasoning more about the incoherent gene pairs by focusing on more complex subgraphs in the signaling network.

### 3.2 The ratio of coherency on subgraphs

In this step, we explored whether complex subgraphs are coherent comparing with considering single edges. Otherwise speaking, we assumed that observing some incoherency of activation and inhibition edges depend on the complex structure of signaling network and logical behavior of larger subgraphs should be considered to infer coherency (Figure 5). Similar to the simple activation or inhibition edges, correlations are computed and categorized, considering the correlation sign for each mentioned subgraph and the calculated adjusted p-values. The details are also available in Supplementary file 2 for all the four analyses. For example, we expected that the portion of significant positive correlations are more than negative correlations in DNFBL as a negative feedback loop comparing with DPFBL1 and DPFBL2. Because the two edges of the DNFBL do not have the same sign and overexpression of one protein entail the underexpression of the other one in a negative feedback loop. However, this expectation only fulfilled in KEGG/GEO analysis partially.

To statistically compare, the correlation analysis was also implemented on multiple sets of 1,000 randomly unconnected gene pairs (UGP) and binomial proportion test was then computed to compare all of the proportions illustrated in Figure 5. Most of the comparison (more than 60%) showed the significant statistical difference between each pairwise proportions of UGP and the connected gene pairs, i.e., Act, Inh, DNFBL, etc. for all four combinations of databases (Supplementary File 3). It would be possible that the connected genes are affected by each other in respect to UGP, but it may happen in a more complex way that it is not inferred by correlation analysis (Figure 5). We also aimed to continue our search to check coherency in larger subgraph structure. We, therefore, identified subgraphs which contain more than two edges, i.e., MNFBLs, MPFBLs, MFFLs, and MNFFLs (See Table 1). However, we did not observe any strong coherent relationship among the gene pairs again, suggesting that, independent of the structure of the subgraph, gene expression profiles do not match the logic of signaling circuits.

## 4. Discussion

High-throughput technologies, such as mRNA microarray, CHIP-seq and mass spectrometry proteomics, has uncovered the amount of expression in mRNA and protein levels (17–20). There is an apparent correspondence between mRNA and protein concentrations. Nonetheless, more than fifty percent of protein variation cannot be explained by variation in mRNA concentration (21–23). These unexplained variations might come from organism-specific translational and post-translational regulations, including protein degradation and gene sequence features (24). The correlation between mRNA and protein concentrations are considerable for some house-keeping genes, but in many eukaryotes, there is no strong correlation for genes of signal transduction or transcriptional regulation. While, these proteins are often involved in different signaling networks, and they determine the cell’s fate and behavior of the system (21). Although the regulation of gene expression results in a particular concentration of proteins, it is not sufficient to completely describe protein abundances (25, 26). The roles of other mechanism such as post-transcriptional, post-translational, and protein degradation regulations has been reported in controlling steady states of protein abundances and activity (25, 27). These modifications had shown their impacts in this study when we illustrated that there is a poor coherency in transducing the signals with the gene expression. These findings are valid for multicellular eukaryotes like the worm and fly. In contrast, yeast genes engaged in signal transduction have high correlations between mRNA and protein concentrations (22). However, Larsen et al. recently demonstrated that there is not any causal relationship between the expression of transcription factors and their targets in the gene regulatory network of E. coli and thereupon the transcriptional regulation cannot be adequately addressed by the current static gene regulatory networks (28). As a result, inferring a gene regulatory or signaling relationships from transcript data is a challenging because this data is not as a proxy of molecular activity. Only in some cases, the results are acceptable for constructing logical circuit of biological elements, e.g., if the components of the system are all, kinases and transition of the signals are related to the phosphorylation process (8, 29, 30). Previously, we also demonstrated the association of altered expressions with the signaling circuits on chronic obstructive pulmonary disease (31) and rabies infection (32) as case studies.

Based on the correlation results in Figure 3A, the volcano plots in Figure 4 which exhibit no significant difference between activation and inhibition edges, the ratios in Figure 5, causal correlation can be inferred poorly at the transcript level at least in a multicellular eukaryotic such as *Homo sapiens.* Proportional binomial tests suggested that there are statistical differences between UGP and other subgraphs (supplementary file 3), and this demonstrates that the structure of subgraphs affects the coherency. It is also strongly advocated to use information in signaling networks, or define relationships between the genes, assess the gene expression at both transcript and protein level or look for the direct interactions.

## 5. Conclusion

Although there is a general assumption that the expression level could strengthen or weaken the signal to transduce in the signaling pathway, but we illustrated that in many instances, there is not a noticeable coherency between the mRNA level of gene pairs and the way (i.e., logic) they manipulate one another (Figure 5). However, we also showed that there is a sort of association between the structure of the subgraphs and gene pair expression profiles. Expression profiles of the unconnected gene pairs were statistically more independent than connected ones. To support this idea, two signaling databases and two gene expression databases were used and the similar results acquired in the analysis of all four combinations.

In this study, we aimed to focus on the impact of the relationship logic on the destination of any given stimulated signaling pathway, which usually ignored in functional genomic studies. We demonstrated that DEGs have only a little information on the whole story of the associated mechanism. Most of these kinds of altered expression are disappeared gradually and ignored by the whole system of signaling network either stimulated endogenously or exogenously.

## Acknowledgent

The authors would like to thank Dr. Julio Saez-Rodriguez, Bence Szalai and Zahra Razaghi-Moghadam for their valuable comments and discussions.

## Supplementary file legends

**Supplementary file 1:** The experimental procedure based on KEGG/GEO analysis in detail. This file contains nine sections. The first section describes how the KEGG edge list with 26,490 edges was built. Next, in the second section, downloading and merging the up-down gene expression profiles was explained for KEGG genes. Section three walks you through the preprocessing of the expression profiles. In this step, an extensive list containing 1,969 experiments (GDS) was built. A large expression matrix called Exprtable with 40,903 samples in column and 3,187 genes in rows was constructed. From this matrix, a list called SignalingNet constructed having an element for each gene pair in the KEGG edge list. In the fourth section, each element of SignalingNet contains the expression values and correlation information for the source and the target genes. Section five includes the information for coherency of the edges and the number of activation and inhibition edges having specific p-value and correlation coefficient. Then, in the sixth section, ten sets of 1,000 unconnected node pairs were built in which the genes never reach one another (based on KEGG information). The correlation analysis was also performed on these node pairs. In the seventh section, the number of edges having specific p-value and correlation coefficient engaged in two-edge subgraphs was computed. Afterward, in the eighth section, the number of edges having specific p-value and correlation coefficient engaged in multiple-edge subgraphs were computed. Finally, in the ninth section, the results were summarized in some tables.

**Supplementary file 2:** Correlation analysis of all four analyses. Results are the number of edges having specific p-values and correlations in different subgraphs.

**Supplementary file 3:** The proportional statistical tests between the rows in the tables in supplementary file 2 for all four analysis in separate sheets.

**Supplementary file 4:** The KEGG and OmniPath edge lists.

**Supplementary file 5:** The large expression matrices constructed based on four analysis analyses.

**Supplementary file 6:** The SignalingNet list for the four analyses.

**Supplementary file 7:**The unconnected SignalingNet list for the four analyses.

