## Supplementary material for "Can we assume the gene expression profile as a proxy for signaling network activity?": 1

Investigating coherency of logical relationship and gene expression of each connected gene pair in the signaling network


### Investigating coherency of logical relationship and gene expression of each connected gene pair in the signaling network

###### *Mehran Piran, Reza Karbalaei, Mohieddin Jafari*

###### *December 12, 2018*

www.Jafarilab.com  


\(~\)  
\(~\)

### Introduction

Study the correlation between the gene expression profiles at both mRNA and protein level is common. Several studies have showed that mRNA and protein expression are poorly correlated. There are some reasons such as miRNA activity on mRNA transcripts or post-translational modifications which cause the variation in amounts of mRNAs and active/inactive Proteins. However, the association between mRNA and protein of housekeeping genes which is not mostly related to signaling processes has been also reported. Now, another major question is the association between mRNA or protein level with the logical relations inside the wiring diagram of the signaling networks. Considering the sign of interactions between components, does the gene expression or protein amount help the signal to be transduced or not? In two simple elements, there are eight possible states (figure 1). Four of these states conform to the direction of the signal transduction, i.e., if the interaction is activation, both of two related components are up-regulated or down-regulated; and if the interaction is inhibitory, the expression of two associated components is inverse (Panel A). The other four possible states are against to signal transduction’s direction (panel B). These two different clusters of states and counting of them in various biological conditions could be an interesting subject for the discussion. Figure 2 is the flowchart which depicts the different steps to reach the data needed to analyze the coherency between the gene pairs in KEGG signaling pathways.

\(~\)

Figure1

\(~\)  
\(~\)

### The Procedure

\(~\)

\(~\)

Figure2: Different steps to analyze the coherency between the gene pairs

\(~\)  
\(~\)

\(~\)

### 1.Creating an edgelist from KEGG database

An edgelist was created from all human KEGG pathways using “KEGGgraph” package.

\(~\)

##### 1.1.Downloading KEGG pathways

All the human signaling pathways were downloaded in form of KGML files from KEGG database KEGG Pathways. There were 206 human pathways on July 24th, 2017.

An example of these pathways is: Ras Signaling Pathway  
\(~\)

##### 1.2.Impoting pathways into R in form of graphs

Downloaded pathways were imported into R using the combination of “parseKGML2Graph” and “lapply” functions. The “parseKGML2Graph” is a function which creates a class of graph (graphNEL) from each pathway and is suitable for networks with few edges and high nodes. All 206 graphs were stored in KGMLGraphs object.

```
library(KEGGgraph)

#Define path
kgfiles<-list.files("F:/Science/Pasteur2/Projects/Project Coherency/KEGG GEO/kegg xml",full.names=T)

#Read kgml files
KGMLGraphs=lapply(kgfiles,parseKGML2Graph,genesOnly=F)
length(KGMLGraphs)
```

```
## [1] 206
```

```
head(KGMLGraphs)
```

```
## [[1]]
## A graphNEL graph with directed edges
## Number of Nodes = 94 
## Number of Edges = 247 
## 
## [[2]]
## A graphNEL graph with directed edges
## Number of Nodes = 115 
## Number of Edges = 311 
## 
## [[3]]
## A graphNEL graph with directed edges
## Number of Nodes = 64 
## Number of Edges = 18 
## 
## [[4]]
## A graphNEL graph with directed edges
## Number of Nodes = 84 
## Number of Edges = 79 
## 
## [[5]]
## A graphNEL graph with directed edges
## Number of Nodes = 93 
## Number of Edges = 282 
## 
## [[6]]
## A graphNEL graph with directed edges
## Number of Nodes = 267 
## Number of Edges = 856
```

\(~\)

##### 1.3.Extracting edge information

Edge information is derived from each pathway using “getKEGGedgeData” function, and they are stored in **l** object.

```
l = lapply(KGMLGraphs,getKEGGedgeData)
length(l)
```

```
## [1] 206
```

\(~\)

##### 1.4. Constructing the edgelists

The following codes construct the edgelists from all 206 signaling pathways and removw non-informative information. The loop contains two “if” conditions to exclude edges without complete data. Note that, nodes (e.g., hsa: 1950) are genes and Subtype is the relationship between genes, e.g., activation, inhibition and so on. A large list called **listedgelists** is created, and each element contains an edgelist.

```
# an example of complete informative edge:
l[[1]][[1]]
```

```
##   KEGG Edge (Type: PPrel):
## ------------------------------------------------------------
## [ Entry 1 ID ]: hsa:1950
## [ Entry 2 ID ]: hsa:1956
## [ Subtype ]: 
##   [ Subtype name ]: activation
##   [ Subtype value ]: -->
## ------------------------------------------------------------
```

```
# First exclusive condition: There is no edge between nodes. For example:
KGMLGraphs[[26]]
```

```
## A graphNEL graph with directed edges
## Number of Nodes = 133 
## Number of Edges = 0
```

```
# Second exclusive condition: There is no information about the edge. For example:
l[[10]][[251]]
```

```
##   KEGG Edge (Type: PCrel):
## ------------------------------------------------------------
## [ Entry 1 ID ]: hsa:6543
## [ Entry 2 ID ]: cpd:C01330
## [ Subtype ]: 
## ------------------------------------------------------------
```

\(~\)

```
a = NULL
class(a)
```

```
## [1] "NULL"
```

```
b = list()

listedgelists = list()

for(i in 1:length(l)){
  if(is.list(l[[i]]) & length(l[[i]]) == 0){
    listedgelists[[i]] = c(0,0,0)
  } else 
  {
  d = matrix(0,length(l[[i]]),3)
  d = as.data.frame(d)
  for(j in 1:length(l[[i]])){
    d[j,1] = l[[i]][[j]]@entry1ID
    d[j,2] = l[[i]][[j]]@entry2ID
    c1 = l[[i]][[j]]@subtype$subtype
    
    if(class(c1) == class(a)){
      d[j,3] =  "none"
    } else
    {
      d[j,3] = l[[i]][[j]]@subtype$subtype@name
    }
    
  }
  
    listedgelists[[i]] = d
  }
}
```

\(~\)

listedgelists object contains all 206 edgelists.

```
head(listedgelists[[1]]) #example of a graph in edge list format
```

```
##         V1       V2         V3
## 1 hsa:1950 hsa:1956 activation
## 2 hsa:7039 hsa:1956 activation
## 3 hsa:3082 hsa:4233 activation
## 4 hsa:4233 hsa:2065 activation
## 5 hsa:3479 hsa:3480 activation
## 6 hsa:7422 hsa:3791 activation
```

```
head(listedgelists[[157]]) #example of a graph in edge list format
```

```
##          V1        V2                  V3
## 1   hsa:718   hsa:727          activation
## 2  hsa:4153 hsa:10747 binding/association
## 3  hsa:4153  hsa:5648 binding/association
## 4 hsa:10747   hsa:720          activation
## 5  hsa:5648   hsa:720          activation
## 6 hsa:10747   hsa:721          activation
```

```
head(listedgelists[[206]]) #example of a graph in edge list format
```

```
##           V1        V2         V3
## 1 cpd:C20930 ko:K07637 activation
## 2  ko:K07637 ko:K07660 activation
## 3  ko:K07660 ko:K18514 activation
## 4  ko:K18514 ko:K12975 repression
## 5  ko:K07660 ko:K12973 expression
## 6  ko:K07660 ko:K07806 expression
```

\(~\)

##### 1.5.Merging all edgelists

All the edgelists are attached together as below, and a large edgelist is created.

```
table = do.call("rbind", listedgelists)
dim(table)
```

```
## [1] 68757     3
```

\(~\)

Duplicated rows are omitted as below.

```
table1 = table[!duplicated(table),]
```

\(~\)

##### 1.6. Preprocessing the edgelist

In the following code, we separated “activation” and “inhibition” interactions from the others.

```
idx1 = which(table1[,3] == "activation")

idx2 = which(table1[,3] == "inhibition")

idx = c(idx1,idx2)

table2 = table1[idx,]

dim(table2)
```

```
## [1] 28870     3
```

\(~\)

After that, only edges which are KEGG Ids, and they start with “hsa” are selected.

```
idx3 = grep("^hsa" , table2[,1])

idx4 = grep("^hsa" , table2[,2])

idx5 = intersect(idx3,idx4)

table3 = table2[idx5,]

dim(table3)
```

```
## [1] 26490     3
```

\(~\)

```
sum(!grepl("^hsa" , table3[,1]))
```

```
## [1] 0
```

```
sum(!grepl("^hsa" , table3[,2]))
```

```
## [1] 0
```

\(~\)

Next, all the KEGG Ids change into gene Ids.

```
kgid1 = as.character(table3[,1]) 

kgid2 = as.character(table3[,2])

geneid1 <- translateKEGGID2GeneID(kgid1)

geneid2 <- translateKEGGID2GeneID(kgid2)

any(is.na(geneid1))
```

```
## [1] FALSE
```

```
any(is.na(geneid1))
```

```
## [1] FALSE
```

\(~\)

Gene Ids are mapped to gene symbols using “org.Hs.eg.db” package which contains annotation for the human genome.

```
require(org.Hs.eg.db)

genesymbol1 <- sapply(mget(geneid1, org.Hs.egSYMBOL, 
                            ifnotfound=NA), "[[",1)
length(genesymbol1)
```

```
## [1] 26490
```

```
any(is.na(genesymbol1))
```

```
## [1] FALSE
```

```
genesymbol2 <- sapply(mget(geneid2, org.Hs.egSYMBOL, 
                           ifnotfound=NA), "[[",1)
length(genesymbol2)
```

```
## [1] 26490
```

```
any(is.na(genesymbol2))
```

```
## [1] FALSE
```

\(~\)

##### 1.7. Final edgelist

So far, there are three columns in the edgelist. The first two columns contain interacting genes and the last one shows interaction type: activation or inhibition

```
edgelist = table3 

edgelist[,1] = genesymbol1

edgelist[,2] = genesymbol2

dim(edgelist)
```

```
## [1] 26490     3
```

\(~\)

Finally, an ID column is added to the edgelist.

```
id1 = paste0("E0000" , 1:9 )
id2 = paste0("E000" , 10:99)
id3 = paste0("E00" , 100:999)
id4 = paste0("E0" , 1000:9999)
id5 = paste0("E" , 10000:length(edgelist[,1]))
ID = c(id1,id2,id3,id4,id5)
edgelist = cbind(ID,edgelist)
colnames(edgelist) = c("ID" , "Gene1" , "Gene2" , "Interaction type")
edgelist = apply(edgelist , 2 , as.character)
edgelist = as.data.frame(edgelist , stringsAsFactors = F)
class(edgelist)
```

```
## [1] "data.frame"
```

\(~\)

```
head(edgelist)
```

```
##       ID Gene1 Gene2 Interaction type
## 1 E00001   EGF  EGFR       activation
## 2 E00002  TGFA  EGFR       activation
## 3 E00003   HGF   MET       activation
## 4 E00004   MET ERBB3       activation
## 5 E00005  IGF1 IGF1R       activation
## 6 E00006 VEGFA   KDR       activation
```

\(~\)  
\(~\)

### 2.Downloading all up or down gene expression profiles

\(~\)  
\(~\)

All the genes are stored at **edgelistGenes** object.

```
edgelistGenes = unique(c(edgelist[,2],edgelist[,3]))
```

```
length(edgelistGenes)
```

```
## [1] 3187
```

\(~\)

There were 3187 unique genes in the KEGG edgelist. We downloaded all the human up-down gene expression profiles for these genes from GEO database (https://www.ncbi.nlm.nih.gov/geoprofiles/). This database contains gene expression profiles derived from curated GEO DataSets and presents them as a chart that displays the expression level of one gene across all Samples within a DataSet. Due to the limitation in NCBI query rules (importing less than 200 gene names in the query and downloading 500 gene profiles in each step), we downloaded 423 text files, each contains nearly 500 gene profiles. A query example for some of the genes is shown here: GEO Profile.

A query example:  
(“Homo sapiens”[Organism] OR homo sapiens[All Fields]) AND (“egf”[Gene Symbol] OR “tgfa”[Gene Symbol] OR “hgf”[Gene Symbol] OR “met”[Gene Symbol] OR “igf1”[Gene Symbol] OR “vegfa”[Gene Symbol] OR “pdgfa”[Gene Symbol] OR “pdgfb”[Gene Symbol] OR “pdgfc”[Gene Symbol] OR “pdgfd”[Gene Symbol] OR “fgf2”[Gene Symbol] OR “gas6”[Gene Symbol] OR “il6”[Gene Symbol] OR “egfr”[Gene Symbol] OR “igf1r”[Gene Symbol] OR “kdr”[Gene Symbol] OR “pdgfra”[Gene Symbol] OR “pdgfrb”[Gene Symbol] OR “fgfr3”[Gene Symbol] OR “fgfr2”[Gene Symbol] OR “axl”[Gene Symbol] OR “il6r”[Gene Symbol] OR “jak1”[Gene Symbol] OR “jak2”[Gene Symbol] OR “src”[Gene Symbol] OR “gab1”[Gene Symbol] OR “akt3”[Gene Symbol] OR “akt1”[Gene Symbol] OR “akt2”[Gene Symbol] OR “mtor”[Gene Symbol] AND “up down genes”[filter])

\(~\)

##### 2.1.Merging all gene expression profiles into one file

All the downloaded gene profiles are merged into Geneprofiles.txt as below (figure3):  
1) all the text files are placed in a folder in drive C.  
2) In the Windows7 terminal window, the following codes were run.  
. cd C: Geneprofiles (Enter)  
. dir (Enter) (Shows all the files in the directory)  
. copy \*.txt Geneprofiles.txt (Enter)

\(~\)

Figure3

\(~\)  
\(~\)

##### 2.2.Importing Geneprofiles file into R

```
library(readxl)
Geneprofiles = read_excel("Geneprofiles.xlsx", col_names = F)
dim(Geneprofiles)
Geneprofiles = as.data.frame(Geneprofiles)
```

\(~\)

\(~\)  
\(~\)

### 3.Processing gene expression profiles

##### 3.1.Creating a large list called listGeneprofiles which contains each part of Geneprofiles dataframe in one of its elements

The following codes are to create a large list called **listGeneprofiles**. each element of the list contains one row for the GDS name, one row for ID\_REF, GSMs, Gene ids and Gene symbol, and multiple rows for probe sets and their values.

An empty list called listGeneprofiles was created. To extract the information of each GDS part, indices of rows containing “GDS”" are found. A “loop” was used to exert some operations on each GDS part. v1 is a vector which contains the row after GDS i (row which start with ID\_REF). “index” is a vector holding all the elements of v1 which start with GSM letters. “l” shows how many GSMs are present in row idx[i] + 1. “d” is a data frame as the same dimention as each GDS part. The number of rows of matrix d, equals to The number of rows which start from GDS[i] to GDS[i+1] minus two (“length((idx[i]):(idx[i+1] - 2))”). The number of columns of object d, equals to the sum of column one, all the GSM columns and two columns after GSM columns (length(c(1,index,index[l]+c(1,2))))). Whenever d is completed for GDS i, extra rows are deleted. That is, row one and rows which contain probesets (have “at” or numbers at the end) are kept.

```
listGeneprofiles = list()
idx = grep("^GDS" , Geneprofiles[,1])
for(i in 1:(length(idx)-1)){
  
  v1 =  data[idx[i]+1,] 
  index = grep("^GSM" , v1)
  l = length(index)
  
  d = as.data.frame(matrix(0,length((idx[i]):(idx[i+1] - 2)) , length(c(1,index,index[l]+c(1,2)))))
  for(j in 1:length((idx[i]):(idx[i+1] - 2))) {
    d[j,] = Geneprofiles[idx[i]+j-1,c(1,index,index[l]+c(2,3))]
  }
  
  colnames(d) = d[2,]
  idxx = grep(paste(paste(c(0:9,'_at'),"$",sep=""),collapse="|"),d[,1])
  listGeneprofiles[[i]] = d[idxx,]
  
  #print(i)
}
```

```
length(listGeneprofiles)
```

```
## [1] 73460
```

\(~\)

listGeneprofiles contains 73460 elements. each element has a similar structure as below.

```
listGeneprofiles[[1]]
```

```
##        ID_REF GSM575 GSM576             GSM577             GSM578 GSM579
## 1       GDS53   <NA>   <NA>               <NA>               <NA>   <NA>
## 4   M22877_at   4419 4528.3 4732.6000000000004 2399.3000000000002 1757.6
## 5 M27968_s_at 2204.9   2417 2456.6999999999998             5088.2 4615.3
##   Gene symbol Gene ID
## 1        <NA>    <NA>
## 4        CYCS   54205
## 5        FGF2    2247
```

```
listGeneprofiles[[57777]]
```

```
##        ID_REF          GSM136055          GSM136056          GSM136057
## 1     GDS2494               <NA>               <NA>               <NA>
## 5 200065_s_at            3630.35             3685.9            3662.81
## 6 202512_s_at 48.658799999999999 49.640799999999999              50.71
## 7 221497_x_at 162.16499999999999            128.107 138.99700000000001
##            GSM136064          GSM136065          GSM136066
## 1               <NA>               <NA>               <NA>
## 5            3622.76            3423.19            3399.93
## 6 72.458799999999997 55.823999999999998 71.829800000000006
## 7 90.802700000000002 82.813800000000001 78.484999999999999
##            GSM136058          GSM136059          GSM136060    Gene symbol
## 1               <NA>               <NA>               <NA>           <NA>
## 5            3083.94            3125.62            3177.01 MIR3620///ARF1
## 6 38.535200000000003 38.714700000000001 43.994900000000001           ATG5
## 7 142.18299999999999            156.648 143.34800000000001          EGLN1
##           Gene ID
## 1            <NA>
## 5 100500810///375
## 6            9474
## 7           54583
```

\(~\)  
\(~\)

##### 3.2.Creating a list called preListGDS which contains each GDS (dataset) in one of its elements

The following codes show how many GDSes (datasets) does exist in the “listGeneprofiles” object.

```
gds = c()
for(i in 1:length(listGeneprofiles)){
  gds[i] = listGeneprofiles[[i]][1,1]
}
gds = unique(gds)
```

```
length(gds)
```

```
## [1] 73460
```

\(~\)

Same datasets were merged as below.

```
preListGDS = list()
for(i in 1:length(gds)){
  idx = which(gds1[i]==gds)
  preListGDS[[i]] = do.call("rbind",listGeneprofiles[idx])
}
length(preListGDS)
```

```
## [1] 1693
```

```
# There are 1693 datasets
```

Figure5

\(~\)

\(~\)

Some datasets does not contain gene symbols or gene ids. We delete them as below.

```
v = c()
for(i in 1:length(preListGDS)){
  v[i]="Gene symbol" %in% colnames(preListGDS[[i]]) & "Gene ID" %in% colnames(preListGDS[[i]])
}

index = which(!v)
index
```

```
## [1]  933 1062 1229 1332
```

```
l = preListGDS[index]

c(l[[1]][1,1],l[[2]][1,1],l[[3]][1,1],l[[4]][1,1])
```

```
## [1] "GDS2771" "GDS3795" "GDS3268" "GDS4206"
```

```
colnames(l[[2]])
```

```
##   [1] "ID_REF"    "GSM483301" "GSM483302" "GSM483303" "GSM483305"
##   [6] "GSM483307" "GSM483312" "GSM483313" "GSM483317" "GSM483318"
##  [11] "GSM483319" "GSM483322" "GSM483327" "GSM483328" "GSM483330"
##  [16] "GSM483332" "GSM483333" "GSM483336" "GSM483337" "GSM483339"
##  [21] "GSM483351" "GSM483352" "GSM483354" "GSM483358" "GSM483384"
##  [26] "GSM483386" "GSM483388" "GSM483390" "GSM483391" "GSM483396"
##  [31] "GSM483399" "GSM483400" "GSM483401" "GSM483412" "GSM483418"
##  [36] "GSM483420" "GSM483421" "GSM483426" "GSM483428" "GSM483431"
##  [41] "GSM483436" "GSM483442" "GSM483443" "GSM483444" "GSM483447"
##  [46] "GSM483448" "GSM483450" "GSM483455" "GSM483458" "GSM483461"
##  [51] "GSM483462" "GSM483464" "GSM483466" "GSM483468" "GSM483476"
##  [56] "GSM483477" "GSM483300" "GSM483308" "GSM483310" "GSM483311"
##  [61] "GSM483323" "GSM483338" "GSM483353" "GSM483361" "GSM483363"
##  [66] "GSM483364" "GSM483366" "GSM483368" "GSM483371" "GSM483372"
##  [71] "GSM483373" "GSM483374" "GSM483379" "GSM483380" "GSM483381"
##  [76] "GSM483389" "GSM483404" "GSM483405" "GSM483410" "GSM483411"
##  [81] "GSM483413" "GSM483416" "GSM483417" "GSM483419" "GSM483427"
##  [86] "GSM483433" "GSM483434" "GSM483445" "GSM483459" "GSM483465"
##  [91] "GSM483470" "GSM483473" "GSM483478" "GSM483304" "GSM483315"
##  [96] "GSM483320" "GSM483325" "GSM483329" "GSM483331" "GSM483334"
## [101] "GSM483341" "GSM483343" "GSM483344" "GSM483347" "GSM483348"
## [106] "GSM483349" "GSM483350" "GSM483356" "GSM483362" "GSM483365"
## [111] "GSM483367" "GSM483369" "GSM483370" "GSM483375" "GSM483376"
## [116] "GSM483377" "GSM483378" "GSM483385" "GSM483402" "GSM483403"
## [121] "GSM483406" "GSM483407" "GSM483408" "GSM483414" "GSM483415"
## [126] "GSM483424" "GSM483437" "GSM483439" "GSM483440" "GSM483446"
## [131] "GSM483449" "GSM483454" "GSM483456" "GSM483460" "GSM483463"
## [136] "GSM483471" "GSM483297" "GSM483298" "GSM483299" "GSM483306"
## [141] "GSM483309" "GSM483314" "GSM483316" "GSM483321" "GSM483324"
## [146] "GSM483326" "GSM483335" "GSM483340" "GSM483342" "GSM483345"
## [151] "GSM483346" "GSM483355" "GSM483357" "GSM483359" "GSM483360"
## [156] "GSM483382" "GSM483383" "GSM483387" "GSM483392" "GSM483393"
## [161] "GSM483394" "GSM483395" "GSM483397" "GSM483398" "GSM483409"
## [166] "GSM483422" "GSM483423" "GSM483425" "GSM483429" "GSM483430"
## [171] "GSM483432" "GSM483435" "GSM483438" "GSM483441" "GSM483451"
## [176] "GSM483452" "GSM483453" "GSM483457" "GSM483467" "GSM483469"
## [181] "GSM483472" "GSM483474" "GSM483475" "GSM483479" "GSM483480"
## [186] "GSM483481" "GSM483482" "GSM483483" "GSM483484" "GSM483485"
## [191] "GSM483486" "GSM483487" "GSM483488" "GSM483489" "0"        
## [196] "0"
```

\(~\)

```
preListGDS = preListGDS[-index]
length(preListGDS)
```

```
## [1] 1689
```

\(~\)

GDS rows are deleted from each element of preListGDS as below.

```
for(i in 1:length(preListGDS)){
  preListGDS[[i]] = preListGDS[[i]][!grepl("^GDS",preListGDS[[i]][,1]),-1]
}
```

\(~\)

##### 3.3.Editing preListGDS object into ListGDS

Datasets of **preListGDS** need to be edited; so, a “for loop” along with 4 conditions are created to edit preListGDS. edited preListGDS is called **ListGDS**. In the gene symbol column of most of GDSes, there are multiple gene symbols for one value. In that case, the gene symbol which is in the edgelist is selected.

```
ListGDS = preListGDS
```

```
# The first condition determines if there is a need for editing

# The second condition is for when there are multiple gene symbols for one value and 
# more than one of them is mapped to the edgelist genes.

# Thr third condition is for when there are multiple gene symbols for one value,
# but none of them mapped to the gene set "Gene"

# The forth condition is for when there are multiple gene symbols for one value and 
# only one of them is mapped to the gene set "Gene""    


for(i in 1:length(ListGDS)){
  
  l1 = length(ListGDS[[i]][1,]) - 1
  
  a = ListGDS[[i]][,l1]
  
  a = strsplit(a,"///",fixed = T)
  
  l = unlist(lapply(a,length))
  
  idxx = which(l>1)
  
    if(length(idxx>0)){
    a1 = a[idxx]
    for(j in 1:length(a1)){
      idxx1 = which(a1[[j]] %in% edgelistGenes)
      if(length(idxx1>1)){
        ListGDS[[i]][,l1][idxx[j]] = a[idxx][[j]][idxx1[1]]
      } else if(length(idxx1)==0){
        ListGDS[[i]][,l1][idxx[j]] = a[idxx][[j]][1]
      } else {
        ListGDS[[i]][,l1][idxx[j]] = a[idxx][[j]][idxx1]
      }
    }
  }
  
  # Deleting regular [A] characters next to the some values
  # changing class of values to numeric
  # changing order of columns
  
  ListGDS[[i]] = cbind(ListGDS[[i]][,l1,drop=F],ListGDS[[i]][,1:(l1-1),drop=F])
  ListGDS[[i]] = as.data.frame(ListGDS[[i]],stringsAsFactors=F)
  
  for(j in 2:l1){
    ListGDS[[i]][,j] = sub("\\[A]","",ListGDS[[i]][,j])
    ListGDS[[i]][,j] = as.numeric(ListGDS[[i]][,j])
  }
   
   #print(i)
}
```

```
head(ListGDS[[1]] , 20)
```

```
##     Gene symbol  GSM575  GSM576  GSM577  GSM578  GSM579
## 4          CYCS  4419.0  4528.3  4732.6  2399.3  1757.6
## 5          FGF2  2204.9  2417.0  2456.7  5088.2  4615.3
## 41         GAS6   765.9   562.4   667.5   364.8   395.6
## 51         GRB2   226.2   212.6   198.2    10.3    34.3
## 42        IL2RB   367.8   508.4   328.2    57.9    34.3
## 52           F3   573.5   607.5   356.9    79.9    21.6
## 6        CASP10  1765.1  1400.3  1843.8   660.6   479.1
## 7         PTPRM   676.4   622.7   611.0   309.4   298.9
## 43       IQGAP1   508.0   482.1   480.6   323.9   328.9
## 44       SREBF1  8303.8  9411.2  8260.5  3804.7  4592.9
## 53         FGD1  2637.9  2035.4  1447.3  5435.8  4638.0
## 61          RHO  6781.8  6509.0  7001.6 10833.1 11141.8
## 45         THRA 21144.2 45869.4 52175.2  9517.2  6752.6
## 46       PLA2R1   721.7   375.6   462.5  5271.3  6598.9
## 47        RAB7A  4117.5  4042.4  2926.3 10662.4 11609.5
## 48         SDC2  1846.2  1794.0  2030.2   656.4   776.9
## 54           C5   167.4    41.9    33.5   326.9   295.8
## 49      PPP2R5A  6597.4  7377.2  8328.5  3951.5  3474.7
## 410        IRF9   445.3   315.8   283.7  2369.0  3024.7
## 411   SERPINB10   570.1   478.5   465.0   123.3   109.5
```

```
head(preListGDS[[1]] , 20)
```

```
##                 GSM575             GSM576             GSM577
## 4                 4419             4528.3 4732.6000000000004
## 5               2204.9               2417 2456.6999999999998
## 41               765.9              562.4              667.5
## 51            226.2[A]           212.6[A]           198.2[A]
## 42               367.8              508.4              328.2
## 52               573.5              607.5              356.9
## 6               1765.1             1400.3             1843.8
## 7                676.4 622.70000000000005                611
## 43              508[A]           482.1[A]           480.6[A]
## 44  8303.7999999999993 9411.2000000000007             8260.5
## 53              2637.9             2035.4             1447.3
## 61              6781.8               6509             7001.6
## 45             21144.2            45869.4 52175.199999999997
## 46               721.7              375.6              462.5
## 47              4117.5             4042.4             2926.3
## 48              1846.2               1794             2030.2
## 54            167.4[A]            41.9[A]            33.5[A]
## 49              6597.4             7377.2             8328.5
## 410              445.3              315.8              283.7
## 411              570.1              478.5                465
##                 GSM578             GSM579 Gene symbol Gene ID
## 4   2399.3000000000002             1757.6        CYCS   54205
## 5               5088.2             4615.3        FGF2    2247
## 41               364.8              395.6        GAS6    2621
## 51             10.3[A]            34.3[A]        GRB2    2885
## 42                57.9            34.3[A]       IL2RB    3560
## 52             79.9[A]               21.6          F3    2152
## 6             660.6[A]           479.1[A]      CASP10     843
## 7             309.4[A] 298.89999999999998       PTPRM    5797
## 43            323.9[A]              328.9      IQGAP1    8826
## 44              3804.7 4592.8999999999996      SREBF1    6720
## 53              5435.8               4638        FGD1    2245
## 61             10833.1            11141.8         RHO    6010
## 45  9517.2000000000007             6752.6        THRA    7067
## 46              5271.3             6598.9      PLA2R1   22925
## 47             10662.4            11609.5       RAB7A    7879
## 48               656.4              776.9        SDC2    6383
## 54  326.89999999999998           295.8[A]          C5     727
## 49              3951.5             3474.7     PPP2R5A    5525
## 410               2369             3024.7        IRF9   10379
## 411           123.3[A]              109.5   SERPINB10    5273
```

```
class(ListGDS[[1]][,4])
```

```
## [1] "numeric"
```

```
# We keep gene symbols and delete columns related to gene Ids

dim(ListGDS[[1]])
```

```
## [1] 49  6
```

```
dim(preListGDS[[1]])
```

```
## [1] 49  7
```

\(~\)

The following code, shows the indices of GDSes which have repetitious GSMs.

```
indexx = list()
for(i in 1:length(ListGDS)){
  
  logic=c()
  
  for(j in (1:length(ListGDS))[-i]){
    logic[j]=any(colnames(ListGDS[[i]])[-1] %in% colnames(ListGDS[[j]]))
  }
  index = which(logic)
  if(length(index) > 0){
    indexx[i] = index
  }
  print(i)
}

index.of.GDSes.having.repetitious.GSMs = which(unlist(lapply(indexx , length)) > 0)
```

\(~\)

In the code below, repetitious GSMs are deleted in different GDSes.

```
for(i in 1:length(ListGDS)){
  
  logic=c()
  for(j in (1:length(ListGDS))[-i]){
    logic[j]=any(colnames(ListGDS[[i]])[-1] %in% colnames(ListGDS[[j]]))
  }
  index = which(logic)
  if(length(index) > 0){
    for(z in 1:length(index)){
      logicc = colnames(ListGDS[[index[z]]]) %in% colnames(ListGDS[[i]])[-1]
      indexx = which(logicc)
      ListGDS[[index[z]]]=ListGDS[[index[z]]][,-indexx]
    }
  }
  # print(i)
}
```

\(~\)

Indices of ListGDS for which there is no GSM left are omitted.

```
index = which(unlist(lapply(lapply(lapply(ListGDS , dim) , function(x) x[2]) , function(x) length(x) == 0)))
ListGDS = ListGDS[-index]
```

```
# All the datasets have more than two GSMs
which(unlist(lapply(lapply(lapply(ListGDS , dim) , function(x) x[2]), function(x) x < 2)))
```

```
## integer(0)
```

\(~\)

Indices of ListGDS for which there is no gene left are removed.

```
index = which(unlist(lapply(lapply(lapply(ListGDS , dim) , function(x) x[1]) , function(x) x < 2)))
ListGDS = ListGDS[-index]
```

\(~\)

All the gene names that are in the ListGDS object, were stored at the object called **profilesGenes**.

```
profilesGenes = c()

for(i in 1:length(ListGDS)){
  profilesGenes = c(profilesGenes,ListGDS[[i]][,1])
}

profilesGenes = unique(profilesGenes)

any(is.na(profilesGenes))

profilesGenes = profilesGenes[-which(is.na(profilesGenes))]
```

```
length(profilesGenes)
```

```
## [1] 3207
```

\(~\)

The first row of the KEGG edgelist contains “EGF” gene. Check how many datasets contain this gene.

```
which(profilesGenes == "EGF")
```

```
## [1] 1679
```

```
logic=c()
for(i in 1:length(ListGDS)){
  logic[i] =  profilesGenes[1679] %in% ListGDS[[i]][,1]
}
idxng1679 = which(logic)
length(which(logic))
```

```
## [1] 44
```

```
head(ListGDS[[idxng1679[20]]],10)
```

```
##    Gene symbol GSM1200171 GSM1200172 GSM1200173 GSM1200174
## 4       PRKACB  3762.2700  3588.7100  5354.8300  5143.6300
## 5          FAS   204.5730   195.6560   507.2090   473.8230
## 6          FAS   178.3230   169.3150   401.8230   366.3830
## 7          EGF    72.6697    66.5497    22.1397    22.2993
## 8         CYCS  8361.6400  8924.8400  5821.4400  6205.0000
## 9          HGF    36.7687    40.6940    85.0087    88.0442
## 10         HGF    51.4297    47.0476    83.9657    83.3615
## 11         FAS   107.4120   103.9990   317.9370   290.5730
## 12         FAS    60.8529    55.7002   161.3550   140.7340
## 13        CYCS    40.3911    42.7323    20.5259    17.2582
```

```
length(profilesGenes) # 3207 is the number of genes extracted from all the gene profiles
```

```
## [1] 3207
```

```
length(edgelistGenes) # 3187 is the number of genes at the edgelist
```

```
## [1] 3187
```

\(~\)

common genes between gene profiles and edgelist were stored in **Genes** object.

```
sum(profilesGenes %in% edgelistGenes)     

Genes = profilesGenes[which(profilesGenes %in% edgelistGenes)]
```

```
length(Genes)
```

```
## [1] 3047
```

```
# There are 3047 common genes between the gene profiles and the edgelist
```

\(~\)

In the folowing code all the GSM names are stored at **gsms** object.

```
list3=list()
for(i in 1:(length(ListGDS))){
  index = grep("^GSM" , colnames(ListGDS[[i]]))
  list3[[i]] = colnames(ListGDS[[i]])[index]
}
length(list3)

gsm = unlist(list3)

gsms = unique(gsm)
```

```
length(gsms)
```

```
## [1] 40903
```

\(~\)  
\(~\)

##### 3.4.Constructing an expression matrix called Exprtable which contains all the gene values amongst the GSMs

First of all, an NA matrix called Exprtable is created which has genes in row and GSMs in column. for each gene name, a sublist is created by a for-loop. For gene one, we check which GDSes of ListGDS object contain it. Then, we store these GDSes at a sublist object. After that, for each element of the sublist, the rows indicating the gene i are stored at **idxx** object. If there exists just one row, we put that row in Exprtable object. Otherwise, the row with larger IQR is located in Exprtable expression matrix.

```
Exprtable = matrix(NA,length(Genes),length(gsms))

Exprtable = as.data.frame(Exprtable)

rownames(Exprtable) = Genes

colnames(Exprtable) = gsms
```

```
dim(Exprtable)
```

```
## [1]  3047 40903
```

```
for(i in 1:length(Genes)){
  
  
  logic = c()
  for(j in 1:length(ListGDS)){
    logic[j] = Genes[i] %in% ListGDS[[j]][,1]
  }
  sublist = ListGDS[logic]
  
 
  for(j in 1:length(sublist)){
    
    gsm = colnames(sublist[[j]])[-1]
      
      idxx = which(sublist[[j]][,1] == Genes[i])
      
      if(length(idxx) == 1){
        Exprtable[Genes[i],gsm] = sublist[[j]][idxx,-1]
      } else {
            d = sublist[[j]][idxx,-1]
            IQRs = c()
           for(z in 1:length(d[,1])){
            a=as.numeric(na.omit(as.numeric(d[z,])))
            IQRs[z] = quantile(a, 0.75 ) - quantile(a, 0.25)
            Exprtable[Genes[i],gsm] = d[which.max(IQRs),]
          }
          
        }
      }
      #print(i)
    }
```

```
sum(is.na(Exprtable))
```

```
## [1] 120876590
```

```
sum(!(is.na(Exprtable)))
```

```
## [1] 3754851
```

\(~\)  
\(~\)

##### 3.5.designing a list called preSignalingNet1 which contains expression values for each edge

In this step, both **Exprtable** and **edglist** are used. For each edge of the edgelist, a data frame is created. First of all, an empty list called preSignalongNet is created. source gene of row i of the edgelist is stored at gene1 object. Target gene of the same row is stored at gene2 object. The source and target gene rows in Exprtable are stored at a and b objects respectively. If there exist more than two non-NA GSMs in both source and target genes, these values are saved as a dataframe in element i of preSignalingNet1 object. If not, we put “empty”" in preSignalingNet1[i].

```
preSignalingNet1 = list()
for(i in 1:length(edgelist[,1])){
  gene1 = edgelist[,2:3][i,1]
  gene2 = edgelist[,2:3][i,2]
  
  idx1 = which(gene1 == rownames(Exprtable))
  idx2 = which(gene2 == rownames(Exprtable))
  a = Exprtable[idx1,]
  b = Exprtable[idx2,]
  l1 = !is.na(a)
  l2 = !is.na(b)
  
  if(length(intersect(which(l1),which(l2))) > 0){
    d = rbind(a,b)
    idx = intersect(which(l1),which(l2))
    preSignalingNet1[[i]] = d[,idx]
    names(preSignalingNet1)[i] = edgelist[i,1]
  } else {
    preSignalingNet1[[i]] = "empty"
    names(preSignalingNet1)[i] = edgelist[i,1]
  }
  #print(i)
}
```

\(~\)

The information of the edge number 10000 is diepicted as below. THBS4 is the source gene and CD47 is the target gene.

```
edgelist[10000,]
```

```
##           ID Gene1 Gene2 Interaction type
## 10000 E10000 THBS4  CD47       activation
```

```
preSignalingNet1[[10000]]
```

```
##       GSM183695 GSM185526 GSM185527 GSM185528 GSM185529 GSM185530
## THBS4   86.5063   80.9516  13.80250   285.017  310.7550   39.8476
## CD47    20.8250   33.1526   3.27527    11.985   32.0414   24.6110
##       GSM185531 GSM185532 GSM185533 GSM185534 GSM185535 GSM185536 GSM47867
## THBS4   521.398 254.00900 686.64900 525.22700  404.1640 733.72700     33.9
## CD47      3.422   7.69714   4.67268   7.08067   29.6316   9.12542     90.9
##       GSM47868 GSM47700 GSM47862 GSM47869 GSM47863 GSM47864 GSM47870
## THBS4     36.0     23.2     31.1     22.2      6.3      6.1     91.4
## CD47      26.7     38.5     50.4     74.3     75.0     44.1      4.8
##       GSM47871 GSM47865 GSM47866 GSM112271 GSM112272 GSM112273 GSM112274
## THBS4     58.3     61.7     56.3   -0.7179   -0.1888   -0.7167   -0.6063
## CD47      22.8      4.3      1.7    0.4725    0.3915    0.0521    0.1217
##       GSM112275 GSM112276 GSM112277 GSM112278 GSM112279 GSM112280
## THBS4   -0.9728   -0.8966   -1.1997   -2.0078   -0.9423   -0.6307
## CD47    -0.5224   -0.4888   -0.8899   -0.9997    0.0867    0.1363
##       GSM112281 GSM112282 GSM112283 GSM112284 GSM112285 GSM112286
## THBS4   -1.0251   -1.0449   -1.2765   -1.3115    0.4004    0.2038
## CD47    -0.5541   -0.5307   -1.0612   -1.1395    0.6933    0.6035
##       GSM112287 GSM112288 GSM112289 GSM112290 GSM112293 GSM112294
## THBS4    0.1565    0.3431   -0.5992   -0.6471    0.1637    0.1986
## CD47     0.2097    0.2109    0.0369    0.0715    0.4232    0.4921
##       GSM112295 GSM112296 GSM112297 GSM112298
## THBS4   -0.8115   -0.6926   -0.8596   -0.8864
## CD47    -0.0582   -0.2459   -0.5638   -0.7999
```

\(~\)

The Object **idx.NO.gsms** has the indices of the edges having less than three GSMs.

```
idx.NO.gsms = which(preSignalingNet1 %in% "empty" )
```

```
length(idx.NO.gsms)
```

```
## [1] 10650
```

\(~\)

##### 3.7.Creating preSignalingNet2

All the values in each element of preSignalingNet1 were separated based on the source of datasets and they are stored at **preSignalingNet2** object. Furthermore, for some indices of preSignalingNet1, source and target genes are the same and they have exactly the same values. These indices are stored at **index.of.not.matching.preSignalingNet.and.ListGDS** object and they are assigned empty in preSignalingNet2 object. **‘These indices are related to loops in the edgelist.’**

```
preSignalingNet2 = preSignalingNet1

index.of.not.matching.preSignalingNet.and.ListGDS=c()

for(i in (1:length(preSignalingNet1))[-idx.NO.gsms]){
  
   
  # Here the index object get indices of all the GDSes which have created signalingnet[[i]] and have the source and target genes 
  logic=c()
  for(j in 1:length(ListGDS)){
    logic[j]=any(colnames(preSignalingNet1[[i]]) %in% colnames(ListGDS[[j]])) & all(rownames(preSignalingNet1[[i]]) %in% ListGDS[[j]][,1])
  }
  index = which(logic)
  
  if(length(index) > 1){
    coexpr = ListGDS[index]
    coEXpr = list()
    colNAMES = lapply(coexpr,colnames)
    for(z in 1:length(coexpr)){
      coEXpr[[z]]=preSignalingNet1[[i]][ , colNAMES[[z]][which( colNAMES[[z]] %in% colnames(preSignalingNet1[[i]]))]]
      
    }
    preSignalingNet2[[i]] = coEXpr
    
  } else if(length(index) == 1){
    preSignalingNet2[[i]] = preSignalingNet1[[i]]
    
  } else { 
    preSignalingNet2[[i]] = "empty"
    index.of.not.matching.preSignalingNet.and.ListGDS[i] = i
  }
  
}

index.of.not.matching.preSignalingNet.and.ListGDS = which(!is.na(unlist(index.of.not.matching.preSignalingNet.and.ListGDS)))
```

\(~\)

```
index.of.not.matching.preSignalingNet.and.ListGDS
```

```
##  [1]   213  5245  5258  5271  5284  5297  5310  5323  5336  5349  5362
## [12]  5375  5388  8711  8715  8719 10057 10060 10077 10080 10081 10082
## [23] 10083 10084 10134 16235 17298 18206 18445 18447 18574 21348 21352
## [34] 21356 25757
```

```
edgelist[index.of.not.matching.preSignalingNet.and.ListGDS[1:4],]
```

```
##          ID   Gene1   Gene2 Interaction type
## 213  E00213   HBEGF   HBEGF       activation
## 5245 E05245 ANAPC10 ANAPC10       activation
## 5258 E05258   CDC26   CDC26       activation
## 5271 E05271 ANAPC13 ANAPC13       activation
```

```
preSignalingNet1[[index.of.not.matching.preSignalingNet.and.ListGDS[2]]][,1:6]
```

```
##          GSM1305903 GSM1305905 GSM1305907 GSM1305909 GSM1305902 GSM1305904
## ANAPC10       49.98      52.17      56.98      54.39      33.34      27.26
## ANAPC101      49.98      52.17      56.98      54.39      33.34      27.26
```

\(~\)

```
Eligibles =  (1:length(preSignalingNet2))[-c(index.of.not.matching.preSignalingNet.and.ListGDS,idx.NO.gsms)]
length(Eligibles)
# There are 15805 edges for down stream analysis
```

\(~\)

Correlation analysis was performed on each edge (gene pair) after preprocessing step. Samples having value for the gene pairs, may come from different datasets and they should be separated and analyzed independently. Figure 6 represents the effect of this preprocessing on a gene pairs in our dataset. In panel A, because all the values are related to multiple datasets in preSignalingNet1[[4]], we can see the very dispersion of the data. However, in the preSignalingNe2[[4]] all of these values are separated into 14 datasets. in panel B, six of these separated datasets are plotted.

Figure 6

\(~\)

\(~\)

```
## null device 
##           1
```

\(~\)  
\(~\)

###### Getting rid of outliers

Values more than 1.5 times of IQR are considered as outliers and are taken out as below.

```
for(i in Eligibles){

  if(class(preSignalingNet2[[i]])=="list"){
for(j in 1:length(preSignalingNet2[[i]]))
  outliers1 = boxplot.stats(as.numeric(preSignalingNet2[[i]][[j]][1,]))$out
  outliers2 = boxplot.stats(as.numeric(preSignalingNet2[[i]][[j]][2,]))$out
  
  index1 = which(as.numeric(preSignalingNet2[[i]][[j]][2,]) %in% outliers1)
  index2 = which(as.numeric(preSignalingNet2[[i]][[j]][2,]) %in% outliers2)
  index = unique(c(index1,index2))
  if(length(index) > 0){
  preSignalingNet2[[i]][[j]] = preSignalingNet2[[i]][[j]][,-index]
  }
  
  } else {
  outliers1 = boxplot.stats(as.numeric(preSignalingNet2[[i]][1,]))$out
  outliers2 = boxplot.stats(as.numeric(preSignalingNet2[[i]][2,]))$out
  
  index1 = which(as.numeric(preSignalingNet2[[i]][1,]) %in% outliers1)
  index2 = which(as.numeric(preSignalingNet2[[i]][2,]) %in% outliers2)
  index = unique(c(index1,index2))
  if(length(index) > 0){
    preSignalingNet2[[i]] = preSignalingNet2[[i]][,-index]}
  
  }
}
```

\(~\)

The following code proves that all the elements of preSignalingNet2 which is in the class dataframe, have more than two samples. The code must return a TRUE value.

```
CLASS = lapply(preSignalingNet2 , class)
all(lapply(lapply(preSignalingNet2[which(CLASS == "data.frame")] , dim), function(x) x[2]) > 2)
```

```
## [1] FALSE
```

\(~\)

The following code presents the number of datasets which have two or fewer datasets.

```
sum(unlist(lapply(preSignalingNet2[which(CLASS == "list")] , function(x) lapply(x,function(x) dim(x)[2] <= 2))))
```

```
## [1] 43
```

\(~\)

The following code shows the indices of preSignalingNet2 which are in the class list and have datasets with less than two samples.

```
index = which(unlist(lapply(preSignalingNet2[which(CLASS == "list")] , function(x) any(unlist(lapply(x,function(x) dim(x)[2] < 3))))))
index
```

```
## E00598 E00868 E00877 E01073 E01081 E01430 E01439 E04464 E04490 E06633 
##    393    550    556    683    689    914    921   2509   2527   3398 
## E06640 E06648 E06655 E06657 E06665 E06672 E06858 E06865 E06873 E06880 
##   3405   3413   3420   3422   3430   3436   3569   3576   3583   3589 
## E06882 E06890 E06897 E07488 E07495 E07503 E07510 E07512 E07520 E07527 
##   3590   3596   3602   4081   4087   4091   4096   4098   4103   4109 
## E09571 E09777 E09842 E13198 E16423 E16428 E17413 E17743 E18042 E18711 
##   4999   5144   5193   6901   7581   7586   8053   8216   8389   8787 
## E25851 E25910 E25914 
##  11007  11050  11054
```

\(~\)

```
preSignalingNet2[which(CLASS == "list")][[index[1]]]
```

```
## [[1]]
##        GSM4312 GSM4317
## TGFB3  296.045 109.721
## TGFBR2  24.545 136.243
## 
## [[2]]
##         GSM7621  GSM7622 GSM7623 GSM7624  GSM6681  GSM6682 GSM6683 GSM6684
## TGFB3   848.063  827.429 3403.71 2631.38  739.542  606.972 3023.23 2892.95
## TGFBR2 5545.760 5872.560 8368.38 7563.18 5431.500 5991.290 7298.31 6956.94
##         GSM6685  GSM6686 GSM6687 GSM6688
## TGFB3   715.008  824.526 3184.55 2850.80
## TGFBR2 4540.660 4318.670 7407.48 6698.04
```

\(~\)

We delete the ineligible datasets from the preSignalingNet2 through the following code. Ineligible datasets are those which contain less than three samples.

```
for(i in 1:length(index)){
  idx = which(unlist(lapply(lapply(preSignalingNet2[which(CLASS == "list")][index][[i]] , dim) , function(x) x[2]<= 2)))
  preSignalingNet2[which(CLASS == "list")][index][[i]] = preSignalingNet2[which(CLASS == "list")][index][[i]][-idx]
}
```

\(~\)

the following code isto check whether there is any element of preSignalingNet2 in class of list and has the length of one.

```
index = which(lapply(preSignalingNet2 , class) == "list" & lapply(preSignalingNet2 , length) == 1)
index
```

```
## E00598 
##    598
```

```
preSignalingNet2[[index]] = as.data.frame(preSignalingNet2[[index]])
```

\(~\)

Because some changes may be imposed on some elements, another checking is done to make sure all the datasets have more than two GSMs.

```
CLASS = lapply(preSignalingNet2 , class)
all(lapply(lapply(preSignalingNet2[which(CLASS == "data.frame")] , dim), function(x) x[2]) > 2)
```

```
## [1] FALSE
```

```
all(unlist(lapply(preSignalingNet2[which(CLASS == "list")] , function(x) any(unlist(lapply(x,function(x) dim(x)[2] >= 3))))))
```

```
## [1] TRUE
```

\(~\)

The elements which contain less than three GSMs are assigned empty.

```
index = which(lapply(lapply(preSignalingNet2[which(CLASS == "data.frame")] , dim), function(x) x[2]) < 3)
index
```

```
## E12586 
##   2460
```

```
if(length(index)>0){
preSignalingNet2[which(CLASS == "data.frame")][[index]]
preSignalingNet2[which(CLASS == "data.frame")][[index]] = "empty"
}
```

\(~\)

Eligible edges are updated.

```
Eligibles =  which(lapply(preSignalingNet2 , class) == "list" | lapply(preSignalingNet2 , class) == "data.frame")
length(Eligibles)
```

```
## [1] 15804
```

\(~\)

Figure 7

### 4.Correlation analysis of each edge

##### 4.1.Creating a large list called SignalingNet which contains expression values and correlation information for each edge

The next step is computing the correlation between the two interacting genes. Pearson, Spearman and Kendall correlation tests are calculated for each edge and a list called **SignalingNet** is created.

```
SignalingNet = preSignalingNet2

for(i in Eligibles){
  
  if(class(SignalingNet[[i]])=="list"){
  corAnalysis = list()
  dim = list()
  for(j in 1:length(SignalingNet[[i]])){
    
  p = cor.test(as.numeric(SignalingNet[[i]][[j]][1,]) , as.numeric(SignalingNet[[i]][[j]][2,]) , method = "pearson")
  s = cor.test(as.numeric(SignalingNet[[i]][[j]][1,]) , as.numeric(SignalingNet[[i]][[j]][2,]) , method = "spearman")
  k = cor.test(as.numeric(SignalingNet[[i]][[j]][1,]) , as.numeric(SignalingNet[[i]][[j]][2,]) , method = "kendall")
  pearson = as.data.frame(cbind(pearsoncor = unname(p$estimate) , pearsonpval = p$p.value))
  spearman = as.data.frame(cbind(spearmancor = unname(s$estimate) , spearmanpval = s$p.value))
  kendall = as.data.frame(cbind(kendallcor = unname(k$estimate) , kendallpval = k$p.value))
  corAnalysis[[j]] = list(pearson = pearson,spearman = spearman,kendall = kendall)
  dim[[j]] = dim(SignalingNet[[i]][[j]])
  }

  SignalingNet[[i]] = list(Source = unname(edgelist[i,2]) , Target = unname(edgelist[i,3])  , length = length(SignalingNet[[i]]) ,dim = dim , coExpr =  SignalingNet[[i]] ,
                             corAnalysis = corAnalysis , Interactiontype = unname(edgelist[i,4]) , Coherency = "empty")

  } else {
    p = cor.test(as.numeric(SignalingNet[[i]][1,]) , as.numeric(SignalingNet[[i]][2,]) , method = "pearson")
    s = cor.test(as.numeric(SignalingNet[[i]][1,]) , as.numeric(SignalingNet[[i]][2,]) , method = "spearman")
    k = cor.test(as.numeric(SignalingNet[[i]][1,]) , as.numeric(SignalingNet[[i]][2,]) , method = "kendall")
    pearson = as.data.frame(cbind(pearsoncor = unname(p$estimate) , pearsonpval = p$p.value))
    spearman = as.data.frame(cbind(spearmancor = unname(s$estimate) , spearmanpval = s$p.value))
    kendall = as.data.frame(cbind(kendallcor = unname(k$estimate) , kendallpval = k$p.value))
    
    
    SignalingNet[[i]] = list(Source = unname(edgelist[i,2]) , Target = unname(edgelist[i,3]) , length = 1 ,dim = dim(SignalingNet[[i]]) , coExpr =  SignalingNet[[i]] ,  
                               corAnalysis = list(pearson = pearson,spearman = spearman,kendall = kendall) , 
                               Interactiontype = unname(edgelist[i,4]) , Coherency = "empty")
  }

}


for(i in (1:length(SignalingNet))[-Eligibles]){
  SignalingNet[[i]] = list(Source = unname(edgelist[i,2]) , Target = unname(edgelist[i,3]) , length = 0 ,dim = 0 , coExpr = NA , corAnalysis = NA , Interactiontype = unname(edgelist[i,4]) , Coherency = "empty")
}
```

\(~\)

###### An example of eligible edge:

```
SignalingNet[[70]]
```

```
## $Source
## [1] "PDGFRB"
## 
## $Target
## [1] "PLCG2"
## 
## $length
## [1] 4
## 
## $dim
## $dim[[1]]
## [1] 2 8
## 
## $dim[[2]]
## [1] 2 8
## 
## $dim[[3]]
## [1]  2 10
## 
## $dim[[4]]
## [1]  2 30
## 
## 
## $coExpr
## $coExpr[[1]]
##        GSM490979 GSM490980 GSM490981 GSM490982 GSM490983 GSM490984
## PDGFRB   8.13064   8.35757   7.80769   8.03344   9.01956   8.54397
## PLCG2    6.11117   6.29475   6.50859   6.23847   5.81704   5.82973
##        GSM490985 GSM490986
## PDGFRB   9.12372   9.15383
## PLCG2    5.92684   6.08239
## 
## $coExpr[[2]]
##        GSM244647 GSM244649 GSM244651 GSM244653 GSM244648 GSM244650
## PDGFRB   36.1238   39.6828   6.70644   13.8024   21.5159   34.3139
## PLCG2   360.3680  373.0510  85.62970   68.4027  428.7340  424.1310
##        GSM244652 GSM244654
## PDGFRB   11.9610   12.7781
## PLCG2    59.5258  104.2860
## 
## $coExpr[[3]]
##        GSM102789 GSM102785 GSM102787 GSM102790 GSM102786 GSM102788
## PDGFRB   176.298   124.255   107.709   352.027   239.746   95.4943
## PLCG2    565.279   259.747   554.512  1922.380   365.583 2091.8100
##        GSM102681 GSM102783 GSM102782 GSM102784
## PDGFRB   1298.46   780.864   951.725   846.224
## PLCG2    1613.89  1324.060  1713.440  1369.790
## 
## $coExpr[[4]]
##        GSM1143676 GSM1143677 GSM1143678 GSM1143679 GSM1143680 GSM1143681
## PDGFRB    7.31532    7.23638    7.54621    7.48674    7.67071    7.35978
## PLCG2     7.30048    7.22943    7.15666    6.98803    7.30929    6.96447
##        GSM1143682 GSM1143683 GSM1143684 GSM1143685 GSM1143686 GSM1143687
## PDGFRB    7.54044    7.39747    7.56700    7.40331    7.44638    7.82191
## PLCG2     7.11817    7.32129    7.08797    6.95786    7.06090    7.56741
##        GSM1143688 GSM1143689 GSM1143690 GSM1143691 GSM1143692 GSM1143693
## PDGFRB    7.54010    7.59168    7.78653    7.47773    7.25271    7.41038
## PLCG2     7.36197    7.49527    7.60299    7.37531    7.32986    7.18851
##        GSM1143694 GSM1143695 GSM1143696 GSM1143697 GSM1143698 GSM1143699
## PDGFRB    7.41699    7.27599    7.48022    7.45763    7.52638    7.78671
## PLCG2     7.19516    7.25258    7.05503    7.03926    7.35168    7.10651
##        GSM1143700 GSM1143701 GSM1143702 GSM1143703 GSM1143704 GSM1143705
## PDGFRB    7.52223    7.06359    7.76968    7.32825    7.41585    7.43055
## PLCG2     7.10267    7.24621    7.28328    7.25294    7.31378    7.46169
## 
## 
## $corAnalysis
## $corAnalysis[[1]]
## $corAnalysis[[1]]$pearson
##   pearsoncor pearsonpval adjusted.pearsonpval
## 1 -0.7402235  0.03573155            0.0790027
## 
## $corAnalysis[[1]]$spearman
##   spearmancor spearmanpval
## 1  -0.7142857   0.05758929
## 
## $corAnalysis[[1]]$kendall
##   kendallcor kendallpval
## 1       -0.5   0.1086806
## 
## 
## $corAnalysis[[2]]
## $corAnalysis[[2]]$pearson
##   pearsoncor pearsonpval adjusted.pearsonpval
## 1  0.8483993 0.007750156           0.02304932
## 
## $corAnalysis[[2]]$spearman
##   spearmancor spearmanpval
## 1   0.6666667   0.08308532
## 
## $corAnalysis[[2]]$kendall
##   kendallcor kendallpval
## 1  0.4285714    0.178869
## 
## 
## $corAnalysis[[3]]
## $corAnalysis[[3]]$pearson
##   pearsoncor pearsonpval adjusted.pearsonpval
## 1  0.4567883   0.1844641            0.2898644
## 
## $corAnalysis[[3]]$spearman
##   spearmancor spearmanpval
## 1   0.2606061    0.4696753
## 
## $corAnalysis[[3]]$kendall
##   kendallcor kendallpval
## 1  0.2444444   0.3807198
## 
## 
## $corAnalysis[[4]]
## $corAnalysis[[4]]$pearson
##   pearsoncor pearsonpval adjusted.pearsonpval
## 1  0.2661583   0.1551258            0.2530683
## 
## $corAnalysis[[4]]$spearman
##   spearmancor spearmanpval
## 1   0.1764182    0.3495161
## 
## $corAnalysis[[4]]$kendall
##   kendallcor kendallpval
## 1  0.1310345   0.3206742
## 
## 
## 
## $Interactiontype
## [1] "activation"
## 
## $Coherency
## [1] "empty"
```

###### An example of ineligible edge:

```
SignalingNet[[43]]
```

```
## $Source
## [1] "IGF1R"
## 
## $Target
## [1] "PLCG1"
## 
## $length
## [1] 0
## 
## $dim
## [1] 0
## 
## $coExpr
## [1] NA
## 
## $corAnalysis
## [1] NA
## 
## $Interactiontype
## [1] "activation"
## 
## $Coherency
## [1] "empty"
```

\(~\)

Using the following code, all the Pearson, Spearman and Kendall correlation coefficients and p-values are saved in the following objects.

```
pearsonpvalues = c()
spearmanpvalues = c()
kendallpvalues = c()
pearsoncors = c()
spearmancors = c()
kendallcors = c()


for(i in Eligibles){
  if(SignalingNet[[i]]$length > 1){
    for(j in 1:length(SignalingNet[[i]]$corAnalysis)){
      
        a = SignalingNet[[i]]$corAnalysis[[j]]$pearson$pearsonpval
        pearsonpvalues = c(pearsonpvalues,a)
        b = SignalingNet[[i]]$corAnalysis[[j]]$spearman$spearmanpval
        spearmanpvalues = c(spearmanpvalues,b)
        c = SignalingNet[[i]]$corAnalysis[[j]]$kendall$kendallpval
        kendallpvalues = c(kendallpvalues,c)
        d = SignalingNet[[i]]$corAnalysis[[j]]$pearson$pearsoncor
        pearsoncors = c(pearsoncors,d)
        e = SignalingNet[[i]]$corAnalysis[[j]]$spearman$spearmancor
        spearmancors = c(spearmancors,e)
        f = SignalingNet[[i]]$corAnalysis[[j]]$kendall$kendallcor
        kendallcors = c(kendallcors,f)
        
      
    }
  }else {
    
      a = SignalingNet[[i]]$corAnalysis$pearson$pearsonpval
      pearsonpvalues = c(pearsonpvalues,a)
      b = SignalingNet[[i]]$corAnalysis$spearman$spearmanpval
      spearmanpvalues = c(spearmanpvalues,b)
      c = SignalingNet[[i]]$corAnalysis$kendall$kendallpval
      kendallpvalues = c(kendallpvalues,c)
      d = SignalingNet[[i]]$corAnalysis$pearson$pearsoncor
      pearsoncors = c(pearsoncors,d)
      e = SignalingNet[[i]]$corAnalysis$spearman$spearmancor
      spearmancors = c(spearmancors,e)
      f = SignalingNet[[i]]$corAnalysis$kendall$kendallcor
      kendallcors = c(kendallcors,f)
    }
    
}
```

```
length(pearsonpvalues)
```

```
## [1] 76897
```

```
length(spearmanpvalues)
```

```
## [1] 76897
```

```
length(kendallpvalues)
```

```
## [1] 76897
```

\(~\)  
\(~\)

Using the codes below, p-values are adjusted using “fdr” method.

```
adjusted.pearsonpvalues = p.adjust(pearsonpvalues , method = "fdr")
```

```
all(lapply(SignalingNet[Eligibles] , function(x) x$length) > 0)
```

```
## [1] TRUE
```

\(~\)

Adjusted p-values are added to the **SignalingNet** object. After completing this operation, adjusted.pearsonpvalues object must be NULL.

```
for(i in Eligibles){
  l = SignalingNet[[i]]$length
  if(l > 1){
for(j in 1:l){
SignalingNet[[i]]$corAnalysis[[j]]$pearson$adjusted.pearsonpval = adjusted.pearsonpvalues[j]
}
 adjusted.pearsonpvalues = adjusted.pearsonpvalues[-1:-l] 
 }else{
    SignalingNet[[i]]$corAnalysis$pearson$adjusted.pearsonpval = adjusted.pearsonpvalues[1]
    adjusted.pearsonpvalues = adjusted.pearsonpvalues[-1]
  }
}
```

Figure8

In the histograms, most of the p-values are under 0.1. The number of positive coefficients are larger than negative coefficients. so, histograms of coefficients are left-skewed.

\(~\)

The codes below show that how many edges have a p-value less than a specified value.

```
cdfpearsonpval = c()
length(pearsonpvalues)
```

```
## [1] 76897
```

```
pvalue = seq(0.01 , 0.99 , by = 0.01)
for(i in 1:99){
  cdfpearsonpval[i] = sum(na.omit(pearsonpvalues < pvalue[i]))
}
cdfpearsonpval = cdfpearsonpval/length(pearsonpvalues)
cdfpearsonpval = rbind(pvalue,cdfpearsonpval)
cdfpearsonpval[,1:5]
```

```
##                     [,1]      [,2]      [,3]      [,4]      [,5]
## pvalue         0.0100000 0.0200000 0.0300000 0.0400000 0.0500000
## cdfpearsonpval 0.3521854 0.4038779 0.4370782 0.4620987 0.4842452
```

```
# For example in the codes below, 35 percent of edges had a pvalue less than 0.01
```

\(~\)

```
cdfspearmanpval = c()
pvalue = seq(0.01 , 0.99 , by = 0.01)
for(i in 1:99){
  cdfspearmanpval[i] = sum(na.omit(spearmanpvalues < pvalue[i]))
}
cdfspearmanpval = cdfspearmanpval/length(spearmanpvalues)
cdfspearmanpval = rbind(pvalue,cdfspearmanpval)
cdfspearmanpval[,1:5]
```

```
##                     [,1]      [,2]      [,3]      [,4]     [,5]
## pvalue          0.010000 0.0200000 0.0300000 0.0400000 0.050000
## cdfspearmanpval 0.259477 0.3160461 0.3524975 0.3870892 0.410094
```

```
cdfkendallpval = c()
pvalue = seq(0.01 , 0.99 , by = 0.01)
for(i in 1:99){
  cdfkendallpval[i] = sum(na.omit(kendallpvalues < pvalue[i]))
}
cdfkendallpval = cdfkendallpval/length(kendallpvalues)
cdfkendallpval = rbind(pvalue,cdfkendallpval)
cdfkendallpval[,1:5]
```

```
##                     [,1]      [,2]      [,3]      [,4]      [,5]
## pvalue         0.0100000 0.0200000 0.0300000 0.0400000 0.0500000
## cdfkendallpval 0.2406726 0.2939256 0.3237578 0.3505208 0.3763086
```

\(~\)

```
cdfpearsoncorr = c()
length(pearsoncors)
```

```
## [1] 76897
```

```
positiveCorr = seq(0 , 0.9 , by = 0.1)
for(i in 1:10){
  cdfpearsoncorr[i] = sum(na.omit(pearsoncors > positiveCorr[i]))
}
cdfpearsoncorr = cdfpearsoncorr/length(pearsoncors)
cdfpearsoncorr = rbind(positiveCorr,cdfpearsoncorr)
# about 10% of edges have pearson corr larger than 0.9
cdfpearsoncorr
```

```
##                     [,1]      [,2]      [,3]      [,4]      [,5]      [,6]
## positiveCorr   0.0000000 0.1000000 0.2000000 0.3000000 0.4000000 0.5000000
## cdfpearsoncorr 0.5919607 0.5378103 0.4845313 0.4329558 0.3831749 0.3341223
##                     [,7]      [,8]      [,9]     [,10]
## positiveCorr   0.6000000 0.7000000 0.8000000 0.9000000
## cdfpearsoncorr 0.2848486 0.2310753 0.1724905 0.1018245
```

\(~\)

```
cdfpearsoncorr1 = c()
length(pearsoncors)
```

```
## [1] 76897
```

```
negativeCorr = seq(0 , -0.9 , by = -0.1)
for(i in 1:10){
  cdfpearsoncorr1[i] = sum(na.omit(pearsoncors < negativeCorr[i]))
}
cdfpearsoncorr1 = cdfpearsoncorr1/length(pearsoncors)
cdfpearsoncorr1 = rbind(negativeCorr,cdfpearsoncorr1)
# about 3.9% of edges have pearson corr less than -0.9
cdfpearsoncorr1
```

```
##                      [,1]       [,2]       [,3]       [,4]       [,5]
## negativeCorr    0.0000000 -0.1000000 -0.2000000 -0.3000000 -0.4000000
## cdfpearsoncorr1 0.4064528  0.3499096  0.2969817  0.2505169  0.2071992
##                      [,6]       [,7]        [,8]        [,9]       [,10]
## negativeCorr    -0.500000 -0.6000000 -0.70000000 -0.80000000 -0.90000000
## cdfpearsoncorr1  0.168212  0.1320208  0.09957476  0.06936551  0.03939035
```

\(~\)

```
cdfspearmancorr = c()
positiveCorr = seq(0 , 0.9 , by = 0.1)
for(i in 1:10){
  cdfspearmancorr[i] = sum(na.omit(spearmancors > positiveCorr[i]))
}
cdfspearmancorr = cdfspearmancorr/length(spearmancors)
cdfspearmancorr = rbind(positiveCorr,cdfspearmancorr)
# about 3.5% of edges have spearman corr larger than 0.9
cdfspearmancorr
```

```
##                      [,1]      [,2]      [,3]      [,4]      [,5]
## positiveCorr    0.0000000 0.1000000 0.2000000 0.3000000 0.4000000
## cdfspearmancorr 0.6002185 0.5422188 0.4857667 0.4329688 0.3782202
##                      [,6]      [,7]      [,8]      [,9]      [,10]
## positiveCorr    0.5000000 0.6000000 0.7000000 0.8000000 0.90000000
## cdfspearmancorr 0.3209748 0.2557057 0.1897083 0.1024227 0.03524195
```

\(~\)

```
cdfspearmancorr1 = c()
negativeCorr = seq(0 , -0.9 , by = -0.1)
for(i in 1:10){
  cdfspearmancorr1[i] = sum(na.omit(spearmancors < negativeCorr[i]))
}
cdfspearmancorr1 = cdfspearmancorr1/length(spearmancors)
cdfspearmancorr1 = rbind(negativeCorr,cdfspearmancorr1)
# about 1.2% of edges have spearman core less than -0.9
cdfspearmancorr1
```

```
##                     [,1]      [,2]       [,3]       [,4]       [,5]
## negativeCorr     0.00000 -0.100000 -0.2000000 -0.3000000 -0.4000000
## cdfspearmancorr1 0.39523  0.339155  0.2878396  0.2420901  0.2003329
##                        [,6]       [,7]        [,8]        [,9]       [,10]
## negativeCorr     -0.5000000 -0.6000000 -0.70000000 -0.80000000 -0.90000000
## cdfspearmancorr1  0.1601883  0.1183401  0.08026321  0.03743969  0.01204208
```

\(~\)

```
cdfkendallcorr = c()
positiveCorr = seq(0 , 0.9 , by = 0.1)
for(i in 1:10){
  cdfkendallcorr[i] = sum(na.omit(kendallcors > positiveCorr[i]))
}
cdfkendallcorr = cdfkendallcorr/length(kendallcors)
cdfkendallcorr = rbind(positiveCorr,cdfkendallcorr)
# about 0.8% of edges have kendall corr larger than 0.9
cdfkendallcorr
```

```
##                    [,1]      [,2]      [,3]      [,4]     [,5]      [,6]
## positiveCorr   0.000000 0.1000000 0.2000000 0.3000000 0.400000 0.5000000
## cdfkendallcorr 0.595966 0.5151306 0.4342172 0.3588176 0.277228 0.1983823
##                     [,7]       [,8]       [,9]       [,10]
## positiveCorr   0.6000000 0.70000000 0.80000000 0.900000000
## cdfkendallcorr 0.1231387 0.06464491 0.02395412 0.008517888
```

\(~\)

```
cdfkendallcorr1 = c()
negativeCorr = seq(0 , -0.9 , by = -0.1)
for(i in 1:10){
  cdfkendallcorr1[i] = sum(na.omit(kendallcors < negativeCorr[i]))
}
cdfkendallcorr1 = cdfkendallcorr1/length(kendallcors)
cdfkendallcorr1 = rbind(negativeCorr,cdfkendallcorr1)
# about 0.32% of edges have kendall core less than -0.9
cdfkendallcorr1
```

```
##                      [,1]       [,2]       [,3]       [,4]       [,5]
## negativeCorr    0.0000000 -0.1000000 -0.2000000 -0.3000000 -0.4000000
## cdfkendallcorr1 0.3903793  0.3138874  0.2426232  0.1856639  0.1293549
##                        [,6]        [,7]        [,8]        [,9]
## negativeCorr    -0.50000000 -0.60000000 -0.70000000 -0.80000000
## cdfkendallcorr1  0.08337126  0.04706295  0.02379807  0.00923313
##                        [,10]
## negativeCorr    -0.900000000
## cdfkendallcorr1  0.004200424
```

\(~\)

Here we create a function for our database “SignalingNet” to check the number of its elements having less than n GSMs amonge eligible elements.

```
GSMnumber = function(x){logic = c()

for(i in Eligibles){
  if(SignalingNet[[i]]$length > 1){
  logic[i] = dim(do.call("cbind" , SignalingNet[[i]]$coExpr))[2] < x
  }else{logic[i] = dim(SignalingNet[[i]]$coExpr)[2] < x}
}
length(which(logic))
}

# to show the number of edges that have less than 10 GSMs:
GSMnumber(10)
```

```
## [1] 1565
```

\(~\)

### 5.One edge Subgraphs

##### 5.1.Getting the number of Activation and Inhibition edges having p-values and correlations of interest.

\(~\)  
In this section the number of edges having the specific pvalue and correlation are computed. The first condition determines if SignalingNet[[i]]$coranalysis has one element or multiple elements. If there is more than one element, all the elements should be non-NA. Then we check that all the elements of SignalingNet[[i]] have p-value < 0.05 and correlation > 0 . If all the conditions are met, the index of the edge is stored at **index.pval.cor1** object.

```
index.pval.cor1 = c()

for(i in Eligibles){
  logic =c()
  if(SignalingNet[[i]]$length > 1){
    
    logic1=c()
    for(j in 1:SignalingNet[[i]]$length){
      logic1[j] = !any(unlist(lapply(SignalingNet[[i]]$corAnalysis[[j]],is.na)))
    }
    if(all(logic1)){
      
  for(j in 1:SignalingNet[[i]]$length){
    
  logic[j] = SignalingNet[[i]]$corAnalysis[[j]]$pearson$adjusted.pearsonpval < 0.05 & SignalingNet[[i]]$corAnalysis[[j]]$pearson$pearsoncor > 0 
  }
  if(all(logic)) {
    index.pval.cor1 = c(index.pval.cor1,i)
  }
  }
  } else { if(!any(unlist(lapply(SignalingNet[[i]]$corAnalysis,is.na)))){
    a = SignalingNet[[i]]$corAnalysis$pearson$adjusted.pearsonpval < 0.05 & SignalingNet[[i]]$corAnalysis$pearson$pearsoncor > 0 
    if(a){
      index.pval.cor1 = c(index.pval.cor1,i)
      }
  }
  }
 
}
```

\(~\)  
\(~\)

The first condition determines if there are more than one GDS for each edge. The second condition tells we need edges for which all the Pearson p-values are NA or > 0.05. If all the conditions are met, the index of the edge is stored at **index.pval.cor.NA** object.

```
index.pval.cor.NA = c()

for(i in Eligibles){
  logic = c()
  if(SignalingNet[[i]]$length > 1){
    
    for(j in 1:SignalingNet[[i]]$length){
      
      logic[j]=any(unlist(lapply(SignalingNet[[i]]$corAnalysis[[j]],is.na))) | SignalingNet[[i]]$corAnalysis[[j]]$pearson$adjusted.pearsonpval > 0.05
    }
    if(all(logic)){ index.pval.cor.NA = c(index.pval.cor.NA,i) }
    
  }else{ if(any(unlist(lapply(SignalingNet[[i]]$corAnalysis,is.na)))| SignalingNet[[i]]$corAnalysis$pearson$adjusted.pearsonpval > 0.05){
    index.pval.cor.NA = c(index.pval.cor.NA,i)}
  }
}
```

\(~\)  
\(~\)

The first condition determines whether SignalingNet[[i]]$coranalysis has one element or multiple elements. If there are several elements, all of them should be non-NA. Then we check that all elements of SignalingNet[[i]] have the p-value < 0.05 or correlation < 0 . If all the conditions are met, the index of the edge will be stored at **index.pval.cor2** object.

```
index.pval.cor2 = c()

for(i in Eligibles){
  logic =c()
  if(SignalingNet[[i]]$length > 1){
    
    logic1=c()
    for(j in 1:SignalingNet[[i]]$length){
      logic1[j] = !any(unlist(lapply(SignalingNet[[i]]$corAnalysis[[j]],is.na)))
    }
    if(all(logic1)){
      
      for(j in 1:SignalingNet[[i]]$length){
        
        logic[j] = SignalingNet[[i]]$corAnalysis[[j]]$pearson$adjusted.pearsonpval < 0.05 & SignalingNet[[i]]$corAnalysis[[j]]$pearson$pearsoncor < 0 
      }
      if(all(logic)) {
        index.pval.cor2 = c(index.pval.cor2,i)
      }
    }
  } else { if(!any(unlist(lapply(SignalingNet[[i]]$corAnalysis,is.na)))){
    a = SignalingNet[[i]]$corAnalysis$pearson$adjusted.pearsonpval < 0.05 & SignalingNet[[i]]$corAnalysis$pearson$pearsoncor < 0 
    if(a){
      index.pval.cor2 = c(index.pval.cor2,i)
    }
  }
  }
  
}
```

\(~\)  
\(~\)

##### 5.2.Determining coherency of edges

If an edge is activation, and p-value < 0.05 and Pearson correlation > 0, or If an edge is inhibition, and p-value < 0.05 and pearson correlation < 0, the edge is coherent. If edge is activation, and p-value < 0.05 and pearson correlation < 0, or If edge is inhibition, and p-value < 0.05 and Pearson correlation > 0, the edge is incoherent. If p-value > 0.05 or p-value == NA, the edge is NA.

\(~\)  
\(~\)

```
logic = edgelist[index.pval.cor1,4] == "activation"
length(which(logic))
```

```
## [1] 896
```

```
idx.Act1 = which(logic)


logic = edgelist[index.pval.cor1,4] == "inhibition"
length(which(logic))
```

```
## [1] 243
```

```
idx.Inh1 = which(logic)
```

\(~\)

```
for(i in index.pval.cor1){
  if(SignalingNet[[i]][[7]] == "activation"){
    SignalingNet[[i]][[8]] = "Coherent"
  } else {SignalingNet[[i]][[8]] = "Incoherent"}
}
```

\(~\)

```
logic = edgelist[index.pval.cor2,4] == "activation"
length(which(logic))
```

```
## [1] 457
```

```
idx.Act2 = which(logic)


logic = edgelist[index.pval.cor2,4] == "inhibition"
length(which(logic))
```

```
## [1] 130
```

```
idx.Inh2 = which(logic)
```

\(~\)

```
for(i in index.pval.cor2){
  if(SignalingNet[[i]][[7]] == "activation"){
    SignalingNet[[i]][[8]] = "Incoherent"
  } else {SignalingNet[[i]][[8]] = "Coherent"}
}
```

\(~\)

```
logic = edgelist[index.pval.cor.NA,4] == "activation"
length(which(logic))
```

```
## [1] 4602
```

```
idx.Act.NA = which(logic)

logic = edgelist[index.pval.cor.NA,4] == "inhibition"
length(which(logic))
```

```
## [1] 1246
```

```
idx.Inh.NA = which(logic)
```

\(~\)

```
for(i in index.pval.cor.NA){
  SignalingNet[[i]][[8]] = NA
}
```

\(~\)  
\(~\)

```
number.of.coherent.edges=0
number.of.incoherent.edges=0
number.of.NA.edges=0

for(i in c(index.pval.cor1,index.pval.cor2)){
  if(SignalingNet[[i]][[8]] == "Coherent"){number.of.coherent.edges = number.of.coherent.edges + 1
  }else if(SignalingNet[[i]][[8]] == "Incoherent"){number.of.incoherent.edges = number.of.incoherent.edges + 1
  }
}


for(i in index.pval.cor.NA){
   if(is.na(SignalingNet[[i]][[8]])){number.of.NA.edges = number.of.NA.edges + 1}
}
```

```
number.of.coherent.edges
```

```
## [1] 1026
```

```
number.of.incoherent.edges
```

```
## [1] 700
```

```
number.of.NA.edges
```

```
## [1] 5848
```

\(~\)

\(~\)

In the following code, some eligible edges which are heterogeneous are stored at the object called **index.heterogeneous**. Heterogeneous edges are those for which there exist multiple datasets with dissimilar p-values and correlations (Table 2).

```
index.heterogeneous = Eligibles[!(Eligibles %in%  index)]

length(index.heterogeneous )
```

```
## [1] 15803
```

```
length(c(index.pval.cor1 , index.pval.cor2 , index.pval.cor.NA , index.heterogeneous ))
```

```
## [1] 23377
```

```
length(unique(c(index.pval.cor1 , index.pval.cor2 , index.pval.cor.NA , index.heterogeneous )))
```

```
## [1] 15805
```

```
length(Eligibles)
```

```
## [1] 15804
```

```
logic = edgelist[index.heterogeneous,4] == "activation"
length(which(logic))
```

```
## [1] 12693
```

```
logic = edgelist[index.heterogeneous,4] == "inhibition"
length(which(logic))
```

```
## [1] 3110
```

\(~\)

\(~\)

\(~\)

\(~\)

\(~\)

\(~\)

\(~\)

\(~\)

### 6. Unconnected gene pairs

\(~\)

For all the four independent analyses, we randomly select 1000 unconnected gene pairs through adjacency matrix self-multiplication. After that, correlation analysis is done on these gene pair. To do that, a graph is created from the eligible edges in the edgelist, and a non-weighted adjacency matrix is created from the giant component of the graph. For more information refer to the section 8. If an entry between two genes in an adjacency matrix stays zero in all the multiplications, there is no path which could connect these two genes.

```
matpower2 = AdjMatrix  %*% AdjMatrix

matpower3 = matpower2  %*% AdjMatrix

matpower4 = matpower3  %*% AdjMatrix

matpower5 = matpower4  %*% AdjMatrix

matpower6 = matpower5 %*% AdjMatrix

matpower7 = matpower6 %*% AdjMatrix

matpower8 = matpower7 %*% AdjMatrix

matpower9 = matpower8  %*% AdjMatrix

matpower10 = matpower9  %*% AdjMatrix

matpower11 = matpower10  %*% AdjMatrix

matpower12 = matpower11  %*% AdjMatrix

matpower13 = matpower12 %*% AdjMatrix

matpower14 = matpower13 %*% AdjMatrix

matpower15 = matpower14  %*% AdjMatrix

matpower16 = matpower15  %*% AdjMatrix

matpower17 = matpower16  %*% AdjMatrix

matpower18 = matpower17  %*% AdjMatrix

matpower19 = matpower18  %*% AdjMatrix

matpower20 = matpower19  %*% AdjMatrix

matpower21 = matpower20  %*% AdjMatrix

matpower22 = matpower21  %*% AdjMatrix

matpower23 = matpower22  %*% AdjMatrix

matpower24 = matpower23  %*% AdjMatrix

matpower25 = matpower24  %*% AdjMatrix

matpower26 = matpower25  %*% AdjMatrix

matpower27 = matpower26  %*% AdjMatrix

matpower28 = matpower27  %*% AdjMatrix

matpower29 = matpower28  %*% AdjMatrix

matpower30 = matpower29  %*% AdjMatrix

matpower31 = matpower30  %*% AdjMatrix

matpower32 = matpower31  %*% AdjMatrix

matpower33 = matpower32  %*% AdjMatrix

matpower34 = matpower33  %*% AdjMatrix

matpower35 = matpower34  %*% AdjMatrix

matpower36 = matpower35  %*% AdjMatrix

matpower37 = matpower36  %*% AdjMatrix

matpower38 = matpower37  %*% AdjMatrix

matpower39 = matpower38  %*% AdjMatrix

matpower40 = matpower39  %*% AdjMatrix
```

\(~\)

```
rn = rownames(AdjMatrix)
cn = colnames(AdjMatrix)
lzeros = list()
```

\(~\)

In the following code, gene pairs for which the value is zero in all 40 non-weighted adjacency matrices are chosen.

```
for(i in 1:length(matpower40[1,])){
  a = cbind(1,2)
  for(j in 1:length(matpower40[,1])){
    if(matpower2[i,j] == 0 & matpower3[i,j] == 0 & matpower4[i,j] == 0 & matpower5[i,j] == 0 & 
       matpower6[i,j] == 0 & matpower7[i,j] == 0 & matpower8[i,j] == 0 & matpower9[i,j] == 0 &
       matpower10[i,j] == 0 & matpower11[i,j] == 0 & matpower12[i,j] == 0 & matpower13[i,j] == 0 &
       matpower14[i,j] == 0 & matpower15[i,j] == 0 & matpower16[i,j] == 0 & matpower17[i,j] == 0 &
       matpower18[i,j] == 0 & matpower19[i,j] == 0 & matpower20[i,j] == 0 & matpower21[i,j] == 0 &
       matpower22[i,j] == 0 & matpower23[i,j] == 0 & matpower24[i,j] == 0 & matpower25[i,j] == 0 &
       matpower26[i,j] == 0 & matpower27[i,j] == 0 & matpower28[i,j] == 0 & matpower29[i,j] == 0 &
       matpower30[i,j] == 0 & matpower31[i,j] == 0 & matpower32[i,j] == 0 & matpower33[i,j] == 0 &
       matpower34[i,j] == 0 & matpower35[i,j] == 0 & matpower36[i,j] == 0 & matpower37[i,j] == 0 &
       matpower38[i,j] == 0 & matpower39[i,j] == 0 & matpower40[i,j] == 0 & AdjMatrix[i,j] == 0 ){
      
      a = rbind(cbind(rn[i],cn[j]),a)
    }
  }
  
  lzeros[[i]] = a
  print(i)
}


unconnected.edgelist = do.call("rbind" , lzeros)
index = which(unconnected.edgelist[,1]=="1")
unconnected.edgelist=unconnected.edgelist[-index,]
```

\(~\)  
\(~\)

There should be no common edges between the main edgelist and unconnected-gene-pair edgelist. because matpower40 adjacency matrix was driven from the giant component of the graph which was created from eligible-edge edgelist, there may exist some common edges between unconnected.edgelist and the eligible-edge edgelist. The reason why this happens is that some eligible edges are not included in the giant component. These edges should be omitted from the unconnected.edgelist. After that, we randomly selected 2000 edges from this edgelist.

```
e = edgelist[Eligibles,2:3]


index.of.rows.in.x.that.are.in.y  <- function(x,y)
{
  x.vec <- apply(x, 1, paste, collapse = "")
  y.vec <- apply(y, 1, paste, collapse = "")
  index = x.vec %in% y.vec
  return(which(index))
}
Index = index.of.rows.in.x.that.are.in.y(unconnected.edgelist,e)
Index

# unconnected.edgelist = unconnected.edgelist[-Index,]

short.unconnected.edgelist = unconnected.edgelist[seq(1,4000000,2000),]
```

\(~\)  
\(~\)

The loop-form gene pairs are removed through the following code.

```
# 
logic=c()
for(i in 1:2000){
  logic[i] = short.unconnected.edgelist[i,1]==short.unconnected.edgelist[i,2]
}
idxloop = which(logic)
idxloop
short.unconnected.edgelist[idxloop,]
short.unconnected.edgelist = short.unconnected.edgelist[-idxloop,]
```

\(~\)

All the gene names of each dataset are stored as an element of **list.gene** object.

```
list.genes = lapply(ListGDS , function(x) unname(x[,1]))
```

\(~\)

Here, a large list called **pre.unconnected.SignalingNet1** is created. Each element of this list is representative of each gene pair in **short.unconnected.edgelist**, and contains gene expression profiles for that gene pair. Some gene pairs are in multiple datasets and some of them are in one dataset. So, some elements of pre.unconnected.SignalingNet1 contain a list of multiple datasets and some of them contain just one dataset. To construct this object the following instruction is applied using **ListGDS** object:

The first condition is to see which datasets contain gene pairs in edge i of the short.unconnected.edgelist. The second condition tells if there are more than one datasets which satisfy the previous condition. This condition separates each element of pre.unconnected.SignalingNet1 into multiple datasets. The third condition tells that in dataset j, just two rows are related to the gene pairs. If the third condition is not satisfied, then between the similar-gene-named rows, we select the one with the largest IQR.

```
pre.unconnected.SignalingNet1 = list()
```

\(~\)

```
for(i in 1:length(short.unconnected.edgelist[,1])){
  index = which(unlist(lapply(list.genes , function(x) all(short.unconnected.edgelist[i,] %in% x))))
  if(length(index) > 0){
    
    small.list = list()
    for(j in 1:length(index)){
    d = ListGDS[[index[j]]][which(list.genes[[index[j]]] %in% short.unconnected.edgelist[i,]) , ]
    
    if(length(d[,1])==2){
      rownames(d) = d[,1]
      d = d[,-1]
      small.list[[j]] = d
      
    }else{
      n = d[,1]
      n1 = unique(n)
      
      index1 = which(n %in% n1[1])
      index2 = which(n %in% n1[2])
      
      if(length(index1)>1){
        d1 = d[index1,]
       
        IQRs = c()
        for(z in 1:length(d1[,1])){
          a=as.numeric(na.omit(as.numeric(d1[z,-1])))
          IQRs[z] = quantile(a, 0.75 ) - quantile(a, 0.25)
        }
        row.one = d1[which.max(IQRs),]
        
      }else{
        row.one = d[which(d[,1] == n1[1]),]
      }
      
      
      if(length(index2)>1){
        d2 = d[index2,]
        
        IQRs = c()
        for(z in 1:length(d2[,1])){
          a=as.numeric(na.omit(as.numeric(d2[z,-1])))
          IQRs[z] = quantile(a, 0.75 ) - quantile(a, 0.25)
        }
        row.two = d2[which.max(IQRs),]
        
      }else{
        row.two = d[which(d[,1] == n1[2]),]
      }
      
      dNEW = rbind(row.one,row.two)
      rownames(dNEW) = dNEW[,1]
      dNEW = dNEW[,-1]
      small.list[[j]] = dNEW
      
    }
    }
    
    if(j > 1){
    pre.unconnected.SignalingNet1[[i]] = small.list
    }else{ pre.unconnected.SignalingNet1[[i]] = small.list[[1]]}
    
  }
  #print(i)
}
```

\(~\)  
\(~\)

Some elements of pre.unconnected.SignalingNet1 are a list of multiple datasets and are in class list, some elements of pre.unconnected.SignalingNet1 are a dataset and are in class dataframe, and some elements of pre.unconnected.SignalingNet1 are NULL. All the elements of this object which contain datasets are stored at **pre.unconnected.SignalingNet2** object.

```
CLASS = lapply(pre.unconnected.SignalingNet1 , class)
index = which(CLASS == "list" |  CLASS == "data.frame")


pre.unconnected.SignalingNet2 = pre.unconnected.SignalingNet1[index]
```

\(~\)  
\(~\)

##### Getting rid of outliers

```
for(i in 1:length(pre.unconnected.SignalingNet2)){
  
  if(class(pre.unconnected.SignalingNet2[[i]])=="list"){
    for(j in 1:length(pre.unconnected.SignalingNet2[[i]]))
      outliers1 = boxplot.stats(as.numeric(pre.unconnected.SignalingNet2[[i]][[j]][1,]))$out
    outliers2 = boxplot.stats(as.numeric(pre.unconnected.SignalingNet2[[i]][[j]][2,]))$out
    
    index1 = which(as.numeric(pre.unconnected.SignalingNet2[[i]][[j]][2,]) %in% outliers1)
    index2 = which(as.numeric(pre.unconnected.SignalingNet2[[i]][[j]][2,]) %in% outliers2)
    index = unique(c(index1,index2))
    if(length(index) > 0){
      pre.unconnected.SignalingNet2[[i]][[j]] = pre.unconnected.SignalingNet2[[i]][[j]][,-index]
    }
    
  } else {
    outliers1 = boxplot.stats(as.numeric(pre.unconnected.SignalingNet2[[i]][1,]))$out
    outliers2 = boxplot.stats(as.numeric(pre.unconnected.SignalingNet2[[i]][2,]))$out
    
    index1 = which(as.numeric(pre.unconnected.SignalingNet2[[i]][1,]) %in% outliers1)
    index2 = which(as.numeric(pre.unconnected.SignalingNet2[[i]][2,]) %in% outliers2)
    index = unique(c(index1,index2))
    if(length(index) > 0){
      pre.unconnected.SignalingNet2[[i]] = pre.unconnected.SignalingNet2[[i]][,-index]}
    
  }
}
```

\(~\)  
\(~\)

```
CLASS = lapply(pre.unconnected.SignalingNet2 , class)


# The following code show that all the elements of pre.unconnected.SignalingNet2 which is data frame, have more than two samples.       
all(lapply(lapply(pre.unconnected.SignalingNet2[which(CLASS == "data.frame")] , dim), function(x) x[2]) > 2)


lapply(pre.unconnected.SignalingNet2[which(CLASS == "list")] , function(x) lapply(x,dim))


# The following code present the number of datasets with less than three samples. 
sum(unlist(lapply(pre.unconnected.SignalingNet2[which(CLASS == "list")] , function(x) lapply(x,function(x) dim(x)[2] <= 2))))


# The following code shows the indices of pre.unconnected.SignalingNet2 which are in the class list having datasets with less than three samples.        
index = which(!(unlist(lapply(pre.unconnected.SignalingNet2[which(CLASS == "list")] , function(x) all(unlist(lapply(x,function(x) dim(x)[2] > 2)))))))
index
# 92 216 491 582


# We deleted the ineligible datasets from the pre.unconnected.SignalingNet2 through the following code
for(i in 1:length(index)){
  idx = which(unlist(lapply(lapply(pre.unconnected.SignalingNet2[which(CLASS == "list")][index][[i]] , dim) , function(x) x[2]<= 2)))
  pre.unconnected.SignalingNet2[which(CLASS == "list")][index][[i]] = pre.unconnected.SignalingNet2[which(CLASS == "list")][index][[i]][-idx]
  }


# Through the code below we check whether or not there exist elements of pre.unconnected.SignalingNet2 which is in the class list and have the length of one (have just one dataset)
any(lapply(pre.unconnected.SignalingNet2[which(CLASS == "list")] , length) == 1)
```

\(~\)  
\(~\)

##### Correlation analysis

\(~\)

A new list called **unconnected.SignalingNet** is created by the following code which contains the correlation results.

```
unconnected.SignalingNet = pre.unconnected.SignalingNet2


for(i in (1:length(unconnected.SignalingNet))){
  
  if(class(unconnected.SignalingNet[[i]])=="list"){
    corAnalysis = list()
    dim = list()
    for(j in 1:length(unconnected.SignalingNet[[i]])){
      
      p = cor.test(as.numeric(unconnected.SignalingNet[[i]][[j]][1,]) , as.numeric(unconnected.SignalingNet[[i]][[j]][2,]) , method = "pearson")
      s = cor.test(as.numeric(unconnected.SignalingNet[[i]][[j]][1,]) , as.numeric(unconnected.SignalingNet[[i]][[j]][2,]) , method = "spearman")
      k = cor.test(as.numeric(unconnected.SignalingNet[[i]][[j]][1,]) , as.numeric(unconnected.SignalingNet[[i]][[j]][2,]) , method = "kendall")
      pearson = as.data.frame(cbind(pearsoncor = unname(p$estimate) , pearsonpval = p$p.value))
      spearman = as.data.frame(cbind(spearmancor = unname(s$estimate) , spearmanpval = s$p.value))
      kendall = as.data.frame(cbind(kendallcor = unname(k$estimate) , kendallpval = k$p.value))
      corAnalysis[[j]] = list(pearson = pearson,spearman = spearman,kendall = kendall)
      dim[[j]] = dim(unconnected.SignalingNet[[i]][[j]])
    }
    
    unconnected.SignalingNet[[i]] = list(Source = rownames(unconnected.SignalingNet[[i]][[1]])[1] , Target = rownames(unconnected.SignalingNet[[i]][[1]])[2]  , length = length(unconnected.SignalingNet[[i]]) ,dim = dim , coExpr =  unconnected.SignalingNet[[i]] ,
                             corAnalysis = corAnalysis , Interactiontype = "None" , Coherency = "empty")
    
  } else {
    p = cor.test(as.numeric(unconnected.SignalingNet[[i]][1,]) , as.numeric(unconnected.SignalingNet[[i]][2,]) , method = "pearson")
    s = cor.test(as.numeric(unconnected.SignalingNet[[i]][1,]) , as.numeric(unconnected.SignalingNet[[i]][2,]) , method = "spearman")
    k = cor.test(as.numeric(unconnected.SignalingNet[[i]][1,]) , as.numeric(unconnected.SignalingNet[[i]][2,]) , method = "kendall")
    pearson = as.data.frame(cbind(pearsoncor = unname(p$estimate) , pearsonpval = p$p.value))
    spearman = as.data.frame(cbind(spearmancor = unname(s$estimate) , spearmanpval = s$p.value))
    kendall = as.data.frame(cbind(kendallcor = unname(k$estimate) , kendallpval = k$p.value))
    
    
    unconnected.SignalingNet[[i]] = list(Source = unconnected.SignalingNet[[i]])[1] , Target = unconnected.SignalingNet[[i]])[2] , length = 1 ,dim = dim(unconnected.SignalingNet[[i]]) , coExpr =  unconnected.SignalingNet[[i]] ,  
                             corAnalysis = list(pearson = pearson,spearman = spearman,kendall = kendall) , 
                             Interactiontype = "None" , Coherency = "empty")
  }
  #print(i)
}
```

\(~\)

```
# The first 1000 elements of unconnected.SignalingNet object are selected.
unconnected.SignalingNet = unconnected.SignalingNet[1:1000]
```

```
unconnected.SignalingNet[[1]]
```

```
## $Source
## [1] "EGF"
## 
## $Target
## [1] "ZEB2"
## 
## $length
## [1] 2
## 
## $dim
## $dim[[1]]
## [1] 2 8
## 
## $dim[[2]]
## [1]  2 27
## 
## 
## $coExpr
## $coExpr[[1]]
##      GSM1214636 GSM1214637 GSM1214638 GSM1214639 GSM1214640 GSM1214641
## EGF     3.44964    3.40186    3.54159    3.46314     5.0123    5.72753
## ZEB2    7.82780    7.62326    7.01536    7.88351     4.0871    3.76693
##      GSM1214642 GSM1214643
## EGF     5.00763    5.37050
## ZEB2    4.07289    4.38288
## 
## $coExpr[[2]]
##      GSM63318 GSM63321 GSM63326 GSM63331 GSM63333 GSM63334 GSM63316
## EGF     0.445    2.103    0.361    0.531    0.511    0.484    0.945
## ZEB2    0.619    0.733   -0.477   -0.774    0.224    0.663    0.171
##      GSM63329 GSM63324 GSM63339 GSM63323 GSM63322 GSM63313 GSM63314
## EGF     0.067    1.305    2.024    0.872    1.229    0.224    1.196
## ZEB2   -0.668    0.736    0.797    0.916    1.617    0.275    0.992
##      GSM63315 GSM63319 GSM63320 GSM63325 GSM63327 GSM63328 GSM63338
## EGF     1.215       NA    0.479    0.964       NA    0.132    0.540
## ZEB2    0.583    0.682    0.208   -0.239    0.152    0.926    0.573
##      GSM63330 GSM63317 GSM63332 GSM63336 GSM63340 GSM63335
## EGF     0.450    1.846    0.961    1.150    1.162    0.826
## ZEB2    1.878    0.686   -0.467    0.562    1.058    1.113
## 
## 
## $corAnalysis
## $corAnalysis[[1]]
## $corAnalysis[[1]]$pearson
##   pearsoncor  pearsonpval adjusted.pearsonpval
## 1 -0.9728855 4.882808e-05         0.0005027198
## 
## $corAnalysis[[1]]$spearman
##   spearmancor spearmanpval
## 1  -0.8095238   0.02177579
## 
## $corAnalysis[[1]]$kendall
##   kendallcor kendallpval
## 1 -0.5714286   0.0610119
## 
## 
## $corAnalysis[[2]]
## $corAnalysis[[2]]$pearson
##   pearsoncor pearsonpval adjusted.pearsonpval
## 1  0.3031217    0.140771            0.2516881
## 
## $corAnalysis[[2]]$spearman
##   spearmancor spearmanpval
## 1        0.34   0.09677333
## 
## $corAnalysis[[2]]$kendall
##   kendallcor kendallpval
## 1  0.2266667   0.1183434
## 
## 
## 
## $Interactiontype
## [1] "None"
## 
## $Coherency
## [1] "empty"
```

\(~\)

All the p-values and correlations are stored at these objects.

```
pearsonpvalues = c()
spearmanpvalues = c()
kendallpvalues = c()
pearsoncors = c()
spearmancors = c()
kendallcors = c()


for(i in 1:1000){
  if(unconnected.SignalingNet[[i]]$length > 1){
    for(j in 1:length(unconnected.SignalingNet[[i]]$corAnalysis)){
      
      a = unconnected.SignalingNet[[i]]$corAnalysis[[j]]$pearson$pearsonpval
      pearsonpvalues = c(pearsonpvalues,a)
      b = unconnected.SignalingNet[[i]]$corAnalysis[[j]]$spearman$spearmanpval
      spearmanpvalues = c(spearmanpvalues,b)
      c = unconnected.SignalingNet[[i]]$corAnalysis[[j]]$kendall$kendallpval
      kendallpvalues = c(kendallpvalues,c)
      d = unconnected.SignalingNet[[i]]$corAnalysis[[j]]$pearson$pearsoncor
      pearsoncors = c(pearsoncors,d)
      e = unconnected.SignalingNet[[i]]$corAnalysis[[j]]$spearman$spearmancor
      spearmancors = c(spearmancors,e)
      f = unconnected.SignalingNet[[i]]$corAnalysis[[j]]$kendall$kendallcor
      kendallcors = c(kendallcors,f)
      
      
    }
  }else {
    
    a = unconnected.SignalingNet[[i]]$corAnalysis$pearson$pearsonpval
    pearsonpvalues = c(pearsonpvalues,a)
    b = unconnected.SignalingNet[[i]]$corAnalysis$spearman$spearmanpval
    spearmanpvalues = c(spearmanpvalues,b)
    c = unconnected.SignalingNet[[i]]$corAnalysis$kendall$kendallpval
    kendallpvalues = c(kendallpvalues,c)
    d = unconnected.SignalingNet[[i]]$corAnalysis$pearson$pearsoncor
    pearsoncors = c(pearsoncors,d)
    e = unconnected.SignalingNet[[i]]$corAnalysis$spearman$spearmancor
    spearmancors = c(spearmancors,e)
    f = unconnected.SignalingNet[[i]]$corAnalysis$kendall$kendallcor
    kendallcors = c(kendallcors,f)
  }
  
  #print(i)
}
```

\(~\)

All the adjusted Pearson p-values are stored at the **adjusted.pearsonpvalues** object.

```
adjusted.pearsonpvalues = p.adjust(pearsonpvalues , method = "fdr")
adjusted.pearsonpvalues

all(lapply(unconnected.SignalingNet , function(x) x$length) > 0)
```

\(~\)

Adjusted Pearson p-values are added to the unconnected.SignalingNet object. After completing the process, adjusted.pearsonpvalues object should be empty or NULL.

```
for(i in 1:1000){
  l = unconnected.SignalingNet[[i]]$length
  if(l > 1){
    for(j in 1:l){
      unconnected.SignalingNet[[i]]$corAnalysis[[j]]$pearson$adjusted.pearsonpval = adjusted.pearsonpvalues[j]
    }
    adjusted.pearsonpvalues = adjusted.pearsonpvalues[-1:-l] 
  }else{
    unconnected.SignalingNet[[i]]$corAnalysis$pearson$adjusted.pearsonpval = adjusted.pearsonpvalues[1]
    adjusted.pearsonpvalues = adjusted.pearsonpvalues[-1]
  }
}
```

\(~\)

```
adjusted.pearsonpvalues
```

```
## numeric(0)
```

\(~\)

The next codes are for getting the number of edges which are engaged in unconnected gene pairs (Table 2). The indices of gene pairs having p-values > 0.05 or p-values == NA are stored at **index.pval.cor.NA** object.

```
index.pval.cor.NA = c()

for(i in 1:1000){
  logic = c()
  if(unconnected.SignalingNet[[i]]$length > 1){
    
    for(j in 1:unconnected.SignalingNet[[i]]$length){
      
      logic[j]=any(unlist(lapply(unconnected.SignalingNet[[i]]$corAnalysis[[j]],is.na))) | unconnected.SignalingNet[[i]]$corAnalysis[[j]]$pearson$adjusted.pearsonpval > 0.05
    }
    if(all(logic)){ index.pval.cor.NA = c(index.pval.cor.NA,i) }
    
  }else{ if(any(unlist(lapply(unconnected.SignalingNet[[i]]$corAnalysis,is.na)))| unconnected.SignalingNet[[i]]$corAnalysis$pearson$adjusted.pearsonpval > 0.05){
    index.pval.cor.NA = c(index.pval.cor.NA,i)}
  }
}
```

```
length(index.pval.cor.NA)
```

```
## [1] 392
```

\(~\)  
\(~\)

The indices of gene pairs which have p-values < 0.05 and correlation > 0 are stored at **index.pval.cor1** object.

```
index.pval.cor1 = c()

for(i in 1:1000){
  logic =c()
  if(unconnected.SignalingNet[[i]]$length > 1){
    
    logic1=c()
    for(j in 1:unconnected.SignalingNet[[i]]$length){
      logic1[j] = !any(unlist(lapply(unconnected.SignalingNet[[i]]$corAnalysis[[j]],is.na)))
    }
    if(all(logic1)){
      
      for(j in 1:unconnected.SignalingNet[[i]]$length){
        
        logic[j] = unconnected.SignalingNet[[i]]$corAnalysis[[j]]$pearson$adjusted.pearsonpval < 0.05 & unconnected.SignalingNet[[i]]$corAnalysis[[j]]$pearson$pearsoncor > 0 
      }
      if(all(logic)) {
        index.pval.cor1 = c(index.pval.cor1,i)
      }
    }
  } else { if(!any(unlist(lapply(unconnected.SignalingNet[[i]]$corAnalysis,is.na)))){
    a = unconnected.SignalingNet[[i]]$corAnalysis$pearson$adjusted.pearsonpval < 0.05 & unconnected.SignalingNet[[i]]$corAnalysis$pearson$pearsoncor > 0 
    if(a){
      index.pval.cor1 = c(index.pval.cor1,i)
    }
  }
  }
  
}
```

```
length(index.pval.cor1)
```

```
## [1] 52
```

\(~\)  
\(~\)

The indices of gene pairs which have p-values < 0.05 and correlation < 0 are stored at **index.pval.cor1** object.

```
index.pval.cor2 = c()

for(i in 1:1000){
  logic =c()
  if(unconnected.SignalingNet[[i]]$length > 1){
    
    logic1=c()
    for(j in 1:unconnected.SignalingNet[[i]]$length){
      logic1[j] = !any(unlist(lapply(unconnected.SignalingNet[[i]]$corAnalysis[[j]],is.na)))
    }
    if(all(logic1)){
      
      for(j in 1:unconnected.SignalingNet[[i]]$length){
        
        logic[j] = unconnected.SignalingNet[[i]]$corAnalysis[[j]]$pearson$adjusted.pearsonpval < 0.05 & unconnected.SignalingNet[[i]]$corAnalysis[[j]]$pearson$pearsoncor < 0 
      }
      if(all(logic)) {
        index.pval.cor2 = c(index.pval.cor2,i)
      }
    }
  } else { if(!any(unlist(lapply(unconnected.SignalingNet[[i]]$corAnalysis,is.na)))){
    a = unconnected.SignalingNet[[i]]$corAnalysis$pearson$adjusted.pearsonpval < 0.05 & unconnected.SignalingNet[[i]]$corAnalysis$pearson$pearsoncor < 0 
    if(a){
      index.pval.cor2 = c(index.pval.cor2,i)
    }
  }
  }
  
}
```

```
length(index.pval.cor2)
```

```
## [1] 51
```

\(~\)  
\(~\)

```
1000 - 392 - 52 - 51
```

```
## [1] 505
```

\(~\)

\(~\)

\(~\)

\(~\)

\(~\)

\(~\)

\(~\)

\(~\)

### 7.Two Edge Subgraphs

\(~\)  
\(~\)

```
load("Two Edge Subgraphs KEGG GEO.RData")
```

The following code shows if there is any loop among the eligible edges.

```
logic=c()
for(i in Eligibles){
  logic[i] = edgelist[i,2]==edgelist[i,3]
}
idxloop = which(logic)
edgelist[idxloop,]
```

```
## [1] ID               Gene1            Gene2            Interaction type
## <0 rows> (or 0-length row.names)
```

```
# There is no loop
```

\(~\)

Edges which has been participated in two-edge subgraphs are saved at **twoEdges** edgelist. the indices of these edges are stored at **idxTWOedges** object.

```
idxTWOedges = list()
for(i in Eligibles){
  a = edgelist[i,2]
  b = edgelist[i,3]
  
  index = which(edgelist[,2] == b & edgelist[,3] == a)
  if(length(index) > 0){idxTWOedges[[i]] = c(i,index)} 
  
}
idxTWOedges = unlist(idxTWOedges)

length(idxTWOedges)
twoEdges=edgelist[idxTWOedges,]
sum(duplicated(twoEdges))
sum(duplicated(edgelist[2:4]))

twoEdges = twoEdges[!duplicated(twoEdges),]
idxTWOedges = rownames(twoEdges)
idxTWOedges = as.numeric(idxTWOedges)
```

\(~\)

```
length(idxTWOedges)
```

```
## [1] 314
```

```
tail(twoEdges)
```

```
##           ID  Gene1  Gene2 Interaction type idxTWOedges
## 20272 E20272 CDKN1B  CCND2       inhibition       20272
## 25769 E25769  CCND2 CDKN1B       inhibition       25769
## 20273 E20273 CDKN1B  CCND3       inhibition       20273
## 25770 E25770  CCND3 CDKN1B       inhibition       25770
## 21710 E21710 CSNK1E  WWTR1       inhibition       21710
## 21726 E21726  WWTR1 CSNK1E       inhibition       21726
```

```
head(twoEdges)
```

```
##           ID  Gene1  Gene2 Interaction type idxTWOedges
## 21    E00021   EGFR    SRC       activation          21
## 16566 E16566    SRC   EGFR       activation       16566
## 264   E00264   PTK2 PIK3CA       activation         264
## 4387  E04387 PIK3CA   PTK2       activation        4387
## 265   E00265   PTK2 PIK3CB       activation         265
## 4388  E04388 PIK3CB   PTK2       activation        4388
```

\(~\)

Two-edge subgraphs are stored at **two.edge.subgraphs** object. This table is a **bisection edgelist**, and it contains columns which determine the indices of edges that has been participated in two edge subgraphs. Each row of the table presents a two-edge subgraph (Figure9).

```
twoEdges$idxTWOedges = idxTWOedges

which(duplicated(edgelist[,2:4]))

index = seq(1 , length(idxTWOedges) , by = 2 )

two.edge.subgraphs = cbind(twoEdges[index,] , twoEdges[index + 1 ,])
```

Figure 9: The bisection edgelist

\(~\)

DNFBL-two-edge subgrphs are stored at the bisection edgelist called **DNFBL**, DPFBL1-two-edge subgrphs are stored at the bisection edgellist called **DPFBL1** and DPFBL2-two-edge subgrphs are stored at the bisection edgellist called **DPFBL2** (Tables 1 and 2).

```
idxDNFBL.1 = c()
for(i in 1:length(two.edge.subgraphs[,1])){
  idxDNFBL.1[i]= two.edge.subgraphs[i,4]=="activation" & two.edge.subgraphs[i,9]=="inhibition"
}
idxDNFBL.1 = which(idxDNFBL.1)

DNFBL.1 = two.edge.subgraphs[idxDNFBL.1,]
```

```
length(DNFBL.1[,1])
```

```
## [1] 20
```

```
head(DNFBL.1)
```

```
##          ID   Gene1     Gene2 Interaction type idxTWOedges     ID
## 1741 E01741    RAC1      RHOA       activation        1741 E25247
## 1742 E01742    RAC2      RHOA       activation        1742 E25248
## 1743 E01743    RAC3      RHOA       activation        1743 E25249
## 2900 E02900  PRKACB   PPP1R1B       activation        2900 E25642
## 3107 E03107 TNFSF11 TNFRSF11B       activation        3107 E21675
## 4885 E04885    INSR    PTPN11       activation        4885 E25571
##          Gene1   Gene2 Interaction type idxTWOedges
## 1741      RHOA    RAC1       inhibition       25247
## 1742      RHOA    RAC2       inhibition       25248
## 1743      RHOA    RAC3       inhibition       25249
## 2900   PPP1R1B  PRKACB       inhibition       25642
## 3107 TNFRSF11B TNFSF11       inhibition       21675
## 4885    PTPN11    INSR       inhibition       25571
```

```
idxDNFBL.2 = c()
for(i in 1:length(two.edge.subgraphs[,1])){
  idxDNFBL.2[i]=two.edge.subgraphs[i,4]=="inhibition" & two.edge.subgraphs[i,9]=="activation"
}
idxDNFBL.2 = which(idxDNFBL.2)
```

```
idxDNFBL.2
```

```
## integer(0)
```

```
DNFBL = DNFBL.1
```

\(~\)  
\(~\)

```
idxDPFBL1 = c()
for(i in 1:length(two.edge.subgraphs[,1])){
  idxDPFBL1[i]=two.edge.subgraphs[i,4]=="activation" & two.edge.subgraphs[i,9]=="activation"
}
idxDPFBL1 = which(idxDPFBL1)

DPFBL1 = two.edge.subgraphs[idxDPFBL1,]
```

```
length(DPFBL1[,1])
```

```
## [1] 125
```

```
head(DPFBL1)
```

```
##         ID Gene1  Gene2 Interaction type idxTWOedges     ID  Gene1 Gene2
## 21  E00021  EGFR    SRC       activation          21 E16566    SRC  EGFR
## 264 E00264  PTK2 PIK3CA       activation         264 E04387 PIK3CA  PTK2
## 265 E00265  PTK2 PIK3CB       activation         265 E04388 PIK3CB  PTK2
## 266 E00266  PTK2 PIK3CD       activation         266 E04389 PIK3CD  PTK2
## 267 E00267  PTK2 PIK3R1       activation         267 E04390 PIK3R1  PTK2
## 268 E00268  PTK2 PIK3R2       activation         268 E04391 PIK3R2  PTK2
##     Interaction type idxTWOedges
## 21        activation       16566
## 264       activation        4387
## 265       activation        4388
## 266       activation        4389
## 267       activation        4390
## 268       activation        4391
```

\(~\)  
\(~\)

```
idxDPFBL2 = c()
for(i in 1:length(two.edge.subgraphs[,1])){
  idxDPFBL2[i]=two.edge.subgraphs[i,4]=="inhibition" & two.edge.subgraphs[i,4]=="inhibition"
}
idxDPFBL2 = which(idxDPFBL2)

DPFBL2 = two.edge.subgraphs[idxDPFBL2,]
```

```
length(DPFBL2[,1])
```

```
## [1] 12
```

```
head(DPFBL2)
```

```
##           ID  Gene1  Gene2 Interaction type idxTWOedges     ID  Gene1
## 21354 E21354   GLI1   GLI3       inhibition       21354 E21350   GLI3
## 19192 E19192   BCL2    BAX       inhibition       19192 E26437    BAX
## 19202 E19202   BCL2    BAD       inhibition       19202 E19216    BAD
## 19217 E19217 BCL2L1   BAK1       inhibition       19217 E25805   BAK1
## 19841 E19841   CDK2 CDKN1B       inhibition       19841 E19850 CDKN1B
## 19842 E19842   CDK2 CDKN1C       inhibition       19842 E19851 CDKN1C
##        Gene2 Interaction type idxTWOedges
## 21354   GLI1       inhibition       21350
## 19192   BCL2       inhibition       26437
## 19202   BCL2       inhibition       19216
## 19217 BCL2L1       inhibition       25805
## 19841   CDK2       inhibition       19850
## 19842   CDK2       inhibition       19851
```

\(~\)  
\(~\)

##### 7.1.Computing the number of edges constructing in three kinds of two-edge subgraphs

\(~\)

##### DNFBL

```
index = c(DNFBL[,5],DNFBL[,10])


DNFBL.pval.cor1 = 0
DNFBL.pval.cor.NA = 0
DNFBL.pval.cor2 = 0


for(i in index){
  logic =c()
  if(SignalingNet[[i]]$length > 1){
    
    for(j in 1:SignalingNet[[i]]$length){
      logic[j]= !any(unlist(lapply(SignalingNet[[i]]$corAnalysis[[j]],is.na)))
    }
    if(all(logic)){
      logic1 = c()
      logic2 = c()
      for(j in 1:SignalingNet[[i]]$length){
        
        logic1[j] = SignalingNet[[i]]$corAnalysis[[j]]$pearson$adjusted.pearsonpval < 0.05 & SignalingNet[[i]]$corAnalysis[[j]]$pearson$pearsoncor > 0 
        logic2[j] = SignalingNet[[i]]$corAnalysis[[j]]$pearson$adjusted.pearsonpval < 0.05 & SignalingNet[[i]]$corAnalysis[[j]]$pearson$pearsoncor < 0
      }
      if(all(logic1)) {DNFBL.pval.cor1=DNFBL.pval.cor1+1}
      if(all(logic2)) {DNFBL.pval.cor2=DNFBL.pval.cor2+1}
    }
  } else{
    if(!any(unlist(lapply(SignalingNet[[i]]$corAnalysis,is.na)))){
      a = SignalingNet[[i]]$corAnalysis$pearson$adjusted.pearsonpval < 0.05 & SignalingNet[[i]]$corAnalysis$pearson$pearsoncor > 0 
      b = SignalingNet[[i]]$corAnalysis$pearson$adjusted.pearsonpval < 0.05 & SignalingNet[[i]]$corAnalysis$pearson$pearsoncor < 0
      if(a){DNFBL.pval.cor1=DNFBL.pval.cor1+1}
      if(b){DNFBL.pval.cor2=DNFBL.pval.cor2+1}
    }
  }
}


for(i in index){
  logic = c()
  if(SignalingNet[[i]]$length > 1){
    
    for(j in 1:SignalingNet[[i]]$length){
      
      logic[j]=any(unlist(lapply(SignalingNet[[i]]$corAnalysis[[j]],is.na))) | SignalingNet[[i]]$corAnalysis[[j]]$pearson$adjusted.pearsonpval > 0.05
    }
    if(all(logic)){ DNFBL.pval.cor.NA = DNFBL.pval.cor.NA + 1 }
    
  }else{ if(any(unlist(lapply(SignalingNet[[i]]$corAnalysis,is.na)))| SignalingNet[[i]]$corAnalysis$pearson$adjusted.pearsonpval > 0.05){
    index.pval.cor.NA = DNFBL.pval.cor.NA + 1}
  }
}
```

```
DNFBL.pval.cor1
```

```
## [1] 0
```

```
DNFBL.pval.cor.NA
```

```
## [1] 8
```

```
DNFBL.pval.cor2
```

```
## [1] 3
```

```
length(DNFBL[,1]) * 2 - c(DNFBL.pval.cor1 + DNFBL.pval.cor.NA + DNFBL.pval.cor2)
```

```
## [1] 29
```

\(~\)  
\(~\)

##### DPFBL1

```
index = c(DPFBL1[,5] , DPFBL1[,10])


DPFBL1.pval.cor1 = 0
DPFBL1.pval.cor.NA = 0
DPFBL1.pval.cor2 = 0


for(i in index){
  logic =c()
  if(SignalingNet[[i]]$length > 1){
    
    for(j in 1:SignalingNet[[i]]$length){
      logic[j]=(!(class(SignalingNet[[i]]$corAnalysis[[j]]) == "logical" | any(is.na(unlist(SignalingNet[[i]]$corAnalysis[[j]])))))
    }
    if(all(logic)){
      logic1 = c()
      logic2 = c()
      for(j in 1:SignalingNet[[i]]$length){
        
        logic1[j] = SignalingNet[[i]]$corAnalysis[[j]]$pearson$adjusted.pearsonpval < 0.05 & SignalingNet[[i]]$corAnalysis[[j]]$pearson$pearsoncor > 0 
        logic2[j] = SignalingNet[[i]]$corAnalysis[[j]]$pearson$adjusted.pearsonpval < 0.05 & SignalingNet[[i]]$corAnalysis[[j]]$pearson$pearsoncor < 0
      }
      if(all(logic1)) {DPFBL1.pval.cor1=DPFBL1.pval.cor1+1}
      if(all(logic2)) {DPFBL1.pval.cor2=DPFBL1.pval.cor2+1}
    }
  } else if(SignalingNet[[i]]$length == 1) {
    if(!(class(SignalingNet[[i]]$corAnalysis) == "logical" | any(is.na(unlist(SignalingNet[[i]]$corAnalysis))))){
      a = SignalingNet[[i]]$corAnalysis$pearson$adjusted.pearsonpval < 0.05 & SignalingNet[[i]]$corAnalysis$pearson$pearsoncor > 0 
      b = SignalingNet[[i]]$corAnalysis$pearson$adjusted.pearsonpval < 0.05 & SignalingNet[[i]]$corAnalysis$pearson$pearsoncor < 0
      if(a){DPFBL1.pval.cor1=DPFBL1.pval.cor1+1}
      if(b){DPFBL1.pval.cor2=DPFBL1.pval.cor2+1}
    }
  }
}


for(i in index){
  logic =c()
  if(SignalingNet[[i]]$length > 1){
    for(j in 1:SignalingNet[[i]]$length){
      logic[j]=!class(SignalingNet[[i]]$corAnalysis[[j]]) == "logical"
    }
    if(all(logic)){
      logic1 =c()
      for(j in 1:SignalingNet[[i]]$length){
        logic1[j]= is.na(SignalingNet[[i]]$corAnalysis[[j]]$pearson$adjusted.pearsonpval) | SignalingNet[[i]]$corAnalysis[[j]]$pearson$adjusted.pearsonpval > 0.05
      }
      if(all(logic1)){DPFBL1.pval.cor.NA = DPFBL1.pval.cor.NA+1}
    }    
    
  } else {
    if(!class(SignalingNet[[i]]$corAnalysis) == "logical"){
      a = is.na(SignalingNet[[i]]$corAnalysis$pearson$adjusted.pearsonpval) | SignalingNet[[i]]$corAnalysis$pearson$adjusted.pearsonpval > 0.05
      if(a){DPFBL1.pval.cor.NA=DPFBL1.pval.cor.NA+1}
    }
  }
}
```

```
DPFBL1.pval.cor1
```

```
## [1] 34
```

```
DPFBL1.pval.cor.NA
```

```
## [1] 85
```

```
DPFBL1.pval.cor2
```

```
## [1] 6
```

```
length(DPFBL1[,1]) * 2 - c(DPFBL1.pval.cor1 + DPFBL1.pval.cor.NA + DPFBL1.pval.cor2)
```

```
## [1] 125
```

\(~\)  
\(~\)

##### DPFBL2

```
index = c(DPFBL2[,5] , DPFBL2[,10])


DPFBL2.pval.cor1 = 0
DPFBL2.pval.cor.NA = 0
DPFBL2.pval.cor2 = 0


for(i in index){
  logic =c()
  if(SignalingNet[[i]]$length > 1){
    
    for(j in 1:SignalingNet[[i]]$length){
      logic[j]=(!(class(SignalingNet[[i]]$corAnalysis[[j]]) == "logical" | any(is.na(unlist(SignalingNet[[i]]$corAnalysis[[j]])))))
    }
    if(all(logic)){
      logic1 = c()
      logic2 = c()
      for(j in 1:SignalingNet[[i]]$length){
        
        logic1[j] = SignalingNet[[i]]$corAnalysis[[j]]$pearson$adjusted.pearsonpval < 0.05 & SignalingNet[[i]]$corAnalysis[[j]]$pearson$pearsoncor > 0 
        logic2[j] = SignalingNet[[i]]$corAnalysis[[j]]$pearson$adjusted.pearsonpval < 0.05 & SignalingNet[[i]]$corAnalysis[[j]]$pearson$pearsoncor < 0
      }
      if(all(logic1)) {DPFBL2.pval.cor1=DPFBL2.pval.cor1+1}
      if(all(logic2)) {DPFBL2.pval.cor2=DPFBL2.pval.cor2+1}
    }
  } else if(SignalingNet[[i]]$length == 1) {
    if(!(class(SignalingNet[[i]]$corAnalysis) == "logical" | any(is.na(unlist(SignalingNet[[i]]$corAnalysis))))){
      a = SignalingNet[[i]]$corAnalysis$pearson$adjusted.pearsonpval < 0.05 & SignalingNet[[i]]$corAnalysis$pearson$pearsoncor > 0 
      b = SignalingNet[[i]]$corAnalysis$pearson$adjusted.pearsonpval < 0.05 & SignalingNet[[i]]$corAnalysis$pearson$pearsoncor < 0
      if(a){DPFBL2.pval.cor1=DPFBL2.pval.cor1+1}
      if(b){DPFBL2.pval.cor2=DPFBL2.pval.cor2+1}
    }
  }
}


for(i in index){
  logic =c()
  if(SignalingNet[[i]]$length > 1){
    for(j in 1:SignalingNet[[i]]$length){
      logic[j]=!class(SignalingNet[[i]]$corAnalysis[[j]]) == "logical"
    }
    if(all(logic)){
      logic1 =c()
      for(j in 1:SignalingNet[[i]]$length){
        logic1[j]= is.na(SignalingNet[[i]]$corAnalysis[[j]]$pearson$adjusted.pearsonpval) | SignalingNet[[i]]$corAnalysis[[j]]$pearson$adjusted.pearsonpval > 0.05
      }
      if(all(logic1)){DPFBL2.pval.cor.NA = DPFBL2.pval.cor.NA+1}
    }    
    
  } else {
    if(!class(SignalingNet[[i]]$corAnalysis) == "logical"){
      a = is.na(SignalingNet[[i]]$corAnalysis$pearson$adjusted.pearsonpval) | SignalingNet[[i]]$corAnalysis$pearson$adjusted.pearsonpval > 0.05
      if(a){DPFBL2.pval.cor.NA=DPFBL2.pval.cor.NA+1}
    }
  }
}
```

```
DPFBL2.pval.cor1
```

```
## [1] 0
```

```
DPFBL2.pval.cor.NA
```

```
## [1] 10
```

```
DPFBL2.pval.cor2
```

```
## [1] 0
```

```
length(DPFBL2[,1]) * 2 - (DPFBL2.pval.cor1 + DPFBL2.pval.cor.NA + DPFBL2.pval.cor2)
```

```
## [1] 14
```

\(~\)

\(~\)

\(~\)

\(~\)

\(~\)

\(~\)

\(~\)

\(~\)

### 8.Multiple Edge subgraphs

To compute the number of edges which participate in multiple-edge subgraphs, an edgelist was created from eligible edges called **shortEdgelist**.

```
library(igraph)
```

```
shortEdgelist = edgelist[Eligibles,]

shortEdgelist[which(shortEdgelist[,4] == "activation"),4] = 1
shortEdgelist[which(shortEdgelist[,4] == "inhibition"),4] = -1
```

\(~\)

A weighted graph was created from shortEdgelist object, and -1 was assigned to inhibitions and 1 was assigned to activations as weights. Then, the largest component of the eligibile edges in the graph was extracted called **weighted.giant.component**. We changed this object into an adjacency matrix called **WadjaMatrix**. The same processes were done for non-weighted adjacency matrix and an object called **none.weighted.giant.component** was created. Then, **AdjaMatrix** object was created from that object. The diameter of the two giant components was computed and using matrix self-multiplication, the number of edges which are engaged in 8 kinds of multiple-edge subgraphs was computed (Table1 and Table2).

\(~\)  
\(~\)

If the edge i in the edgelist is activation and the entity between the gene pair in one of the weighted matrices equals to their unweighted counterparts while the entities are not zero or NA, this edge is engaged in MFFL1 or MPFBL1 subgraphs.

If the edge i in the edgelist is inhibition and the entity between the gene pair in one of the weighted matrices equals to their unweighted counterparts while the entities are not zero or NA, this edge is engaged in MNFFL2 or MNFBL2 subgraphs.

If the edge i in the edgelist is activation and the entity between the gene pair in one of the weighted matrices does not equal to their unweighted counterparts while the entities are not zero or NA, this edge is engaged in MNFBL1 or MNFFL1 subgraphs.

If the edge i in the edgelist is inhibition and the entity between the gene pair in one of the weighted matrices does not equal to their unweighted counterparts while the entities are not zero or NA, this edge is engaged in MFFL2 or MPFBL2 subgraphs.

```
w = as.numeric(shortEdgelist[,4])
shortEdgelist = as.matrix(shortEdgelist)

g = graph_from_edgelist(shortEdgelist[,2:3] , directed = T)
E(g)$weights = w
```

\(~\)

```
shortEdgelistGenes = unique(c(shortEdgelist[,2] , c(shortEdgelist[,3])))

c= components(g)

weighted.giant.component = induced.subgraph(g , which(c$membership ==  1))
```

```
weighted.giant.component
```

```
## IGRAPH f0c4f7d DN-- 2549 15731 -- 
## + attr: name (v/c), weights (e/n)
## + edges from f0c4f7d (vertex names):
##  [1] EGF  ->EGFR   TGFA ->EGFR   HGF  ->MET    MET  ->ERBB3  IGF1 ->IGF1R 
##  [6] VEGFA->KDR    PDGFA->PDGFRA PDGFB->PDGFRA PDGFC->PDGFRA PDGFD->PDGFRA
## [11] PDGFA->PDGFRB PDGFB->PDGFRB PDGFC->PDGFRB PDGFD->PDGFRB FGF2 ->FGFR3 
## [16] FGF2 ->FGFR2  GAS6 ->AXL    IL6  ->IL6R   EGFR ->JAK1   EGFR ->JAK2  
## [21] EGFR ->SRC    EGFR ->GAB1   EGFR ->PLCG1  EGFR ->PLCG2  EGFR ->SHC2  
## [26] EGFR ->SHC4   EGFR ->SHC3   EGFR ->SHC1   MET  ->JAK1   MET  ->JAK2  
## [31] MET  ->SRC    MET  ->GAB1   MET  ->PLCG1  MET  ->PLCG2  MET  ->SHC2  
## [36] MET  ->SHC4   MET  ->SHC3   MET  ->SHC1   IGF1R->JAK1   IGF1R->JAK2  
## + ... omitted several edges
```

```
is.connected(weighted.giant.component)
```

```
## [1] TRUE
```

```
WadjMatrix = as_adjacency_matrix(weighted.giant.component , attr = "weights")

WadjMatrix = as.matrix(WadjMatrix)
```

```
diameter(weighted.giant.component)
```

```
## [1] 17
```

\(~\)  
\(~\)

```
Wmatpower2 = WadjMatrix  %*% WadjMatrix

Wmatpower3 = Wmatpower2  %*% WadjMatrix

Wmatpower4 = Wmatpower3  %*% WadjMatrix

Wmatpower5 = Wmatpower4  %*% WadjMatrix

Wmatpower6 = Wmatpower5 %*% WadjMatrix

Wmatpower7 = Wmatpower6 %*% WadjMatrix

Wmatpower8 = Wmatpower7 %*% WadjMatrix

Wmatpower9 = Wmatpower8  %*% WadjMatrix

Wmatpower10 = Wmatpower9  %*% WadjMatrix

Wmatpower11 = Wmatpower10  %*% WadjMatrix

Wmatpower12 = Wmatpower11  %*% WadjMatrix

Wmatpower13 = Wmatpower12 %*% WadjMatrix

Wmatpower14 = Wmatpower13 %*% WadjMatrix

Wmatpower15 = Wmatpower14  %*% WadjMatrix

Wmatpower16 = Wmatpower15  %*% WadjMatrix

Wmatpower17 = Wmatpower16  %*% WadjMatrix
```

\(~\)  
\(~\)

```
g1 = graph_from_edgelist(shortEdgelist[,2:3] , directed = T)

c= components(g1)

none.weighted.giant.component = induced.subgraph(g1 , which(c$membership ==  1))
```

```
none.weighted.giant.component
```

```
## IGRAPH 1232f1c DN-- 2549 15731 -- 
## + attr: name (v/c)
## + edges from 1232f1c (vertex names):
##  [1] EGF  ->EGFR   TGFA ->EGFR   HGF  ->MET    MET  ->ERBB3  IGF1 ->IGF1R 
##  [6] VEGFA->KDR    PDGFA->PDGFRA PDGFB->PDGFRA PDGFC->PDGFRA PDGFD->PDGFRA
## [11] PDGFA->PDGFRB PDGFB->PDGFRB PDGFC->PDGFRB PDGFD->PDGFRB FGF2 ->FGFR3 
## [16] FGF2 ->FGFR2  GAS6 ->AXL    IL6  ->IL6R   EGFR ->JAK1   EGFR ->JAK2  
## [21] EGFR ->SRC    EGFR ->GAB1   EGFR ->PLCG1  EGFR ->PLCG2  EGFR ->SHC2  
## [26] EGFR ->SHC4   EGFR ->SHC3   EGFR ->SHC1   MET  ->JAK1   MET  ->JAK2  
## [31] MET  ->SRC    MET  ->GAB1   MET  ->PLCG1  MET  ->PLCG2  MET  ->SHC2  
## [36] MET  ->SHC4   MET  ->SHC3   MET  ->SHC1   IGF1R->JAK1   IGF1R->JAK2  
## + ... omitted several edges
```

```
is.connected(none.weighted.giant.component)
```

```
## [1] TRUE
```

```
AdjMatrix = as_adjacency_matrix(none.weighted.giant.component)

AdjMatrix = as.matrix(AdjMatrix)
```

```
diameter(none.weighted.giant.component)
```

```
## [1] 17
```

\(~\)

```
matpower2 = AdjMatrix  %*% AdjMatrix

matpower3 = matpower2  %*% AdjMatrix

matpower4 = matpower3  %*% AdjMatrix

matpower5 = matpower4  %*% AdjMatrix

matpower6 = matpower5 %*% AdjMatrix

matpower7 = matpower6 %*% AdjMatrix

matpower8 = matpower7 %*% AdjMatrix

matpower9 = matpower8  %*% AdjMatrix

matpower10 = matpower9  %*% AdjMatrix

matpower11 = matpower10  %*% AdjMatrix

matpower12 = matpower11  %*% AdjMatrix

matpower13 = matpower12 %*% AdjMatrix

matpower14 = matpower13 %*% AdjMatrix

matpower15 = matpower14  %*% AdjMatrix

matpower16 = matpower15  %*% AdjMatrix

matpower17 = matpower16  %*% AdjMatrix
```

\(~\)  
\(~\)

**listAdj1** object is a large list containing matrices as the same dimension as the adjacency matrices. This list contains 16 matrices, and Each matrix is representative of one of the powered matrices.

```
Apower = matrix(0 , 2549 , 2549)
colnames(Apower) = colnames(AdjMatrix)
rownames(Apower) = rownames(AdjMatrix)
listAdj1 = list(Apower2 = Apower,
Apower3 = Apower,
Apower4 = Apower,
Apower5 = Apower,
Apower6 = Apower,
Apower7 = Apower,
Apower8 = Apower,
Apower9 = Apower,
Apower10 = Apower,
Apower11 = Apower,
Apower12 = Apower,
Apower13 = Apower,
Apower14 = Apower,
Apower15 = Apower,
Apower16 = Apower,
Apower17 = Apower
)
```

\(~\)

Before computing the number of edges which participate in multiple-edge subgraphs, the following algorithms are required to be applied: If WmatpowerX[i,j] == matpowerX[i,j], we put the value of WmatpowerX[i,j] at the ApowerX[i,j] in **listAdj1** list. Otherwise, we put NA. After that, the number of edges which take part in “MPFBL1”, “MNFBL2”, “MFFL1” and “MNFFL2” multiple-edge subgraphs are computed. For the other multiple-edge subgraphs called “MNFBL1”, “MPFBL2”, “MFFL2” and “MNFFL1”, If WmatpowerX[i,j] != matpowerX[i,j], we put the value of WmatpowerX[i,j] at the ApowerX[i,j] in **listAdj2** list. Otherwise, we put NA.

```
for(i in 1:length(AdjMatrix[,1])){
  
for(j in 1:length(AdjMatrix[1,])){

if(Wmatpower2[i,j] == matpower2[i,j]){
  listAdj1$Apower2[i,j] = Wmatpower2[i,j]
} else { listAdj1$Apower2[i,j] = NA }


if(Wmatpower3[i,j] == matpower3[i,j]){
  listAdj1$Apower3[i,j] = Wmatpower3[i,j]
} else { listAdj1$Apower3[i,j] = NA }


if(Wmatpower4[i,j] == matpower4[i,j]){
  listAdj1$Apower4[i,j] = Wmatpower4[i,j]
} else { listAdj1$Apower4[i,j] = NA }


if(Wmatpower5[i,j] == matpower5[i,j]){
  listAdj1$Apower5[i,j] = Wmatpower5[i,j]
} else { listAdj1$Apower5[i,j] = NA }


if(Wmatpower6[i,j] == matpower6[i,j]){
  listAdj1$Apower6[i,j] = Wmatpower6[i,j]
} else { listAdj1$Apower6[i,j] = NA }


if(Wmatpower7[i,j] == matpower7[i,j]){
  listAdj1$Apower7[i,j] = Wmatpower7[i,j]
} else { listAdj1$Apower7[i,j] = NA }


if(Wmatpower8[i,j] == matpower8[i,j]){
  listAdj1$Apower8[i,j] = Wmatpower8[i,j]
} else { listAdj1$Apower8[i,j] = NA }


if(Wmatpower9[i,j] == matpower9[i,j]){
  listAdj1$Apower9[i,j] = Wmatpower9[i,j]
} else { listAdj1$Apower9[i,j] = NA }


if(Wmatpower10[i,j] == matpower10[i,j]){
  listAdj1$Apower10[i,j] = Wmatpower10[i,j]
} else { listAdj1$Apower10[i,j] = NA }


if(Wmatpower11[i,j] == matpower11[i,j]){
  listAdj1$Apower11[i,j] = Wmatpower11[i,j]
} else { listAdj1$Apower11[i,j] = NA }


if(Wmatpower12[i,j] == matpower12[i,j]){
  listAdj1$Apower12[i,j] = Wmatpower12[i,j]
} else { listAdj1$Apower12[i,j] = NA }


if(Wmatpower13[i,j] == matpower13[i,j]){
  listAdj1$Apower13[i,j] = Wmatpower13[i,j]
} else { listAdj1$Apower13[i,j] = NA }


if(Wmatpower14[i,j] == matpower14[i,j]){
  listAdj1$Apower14[i,j] = Wmatpower14[i,j]
} else { listAdj1$Apower14[i,j] = NA }


if(Wmatpower15[i,j] == matpower15[i,j]){
  listAdj1$Apower15[i,j] = Wmatpower15[i,j]
} else { listAdj1$Apower15[i,j] = NA }


if(Wmatpower16[i,j] == matpower16[i,j]){
  listAdj1$Apower16[i,j] = Wmatpower16[i,j]
} else { listAdj1$Apower16[i,j] = NA }


if(Wmatpower17[i,j] == matpower17[i,j]){
  listAdj1$Apower17[i,j] = Wmatpower17[i,j]
} else { listAdj1$Apower17[i,j] = NA }

}
  # print(i)
}
```

\(~\)

```
Apower = matrix(0 , 2549 , 2549)
colnames(Apower) = colnames(AdjMatrix)
rownames(Apower) = rownames(AdjMatrix)
listAdj2 = list(Apower2 = Apower,
               Apower3 = Apower,
               Apower4 = Apower,
               Apower5 = Apower,
               Apower6 = Apower,
               Apower7 = Apower,
               Apower8 = Apower,
               Apower9 = Apower,
               Apower10 = Apower,
               Apower11 = Apower,
               Apower12 = Apower,
               Apower13 = Apower,
               Apower14 = Apower,
               Apower15 = Apower,
               Apower16 = Apower,
               Apower17 = Apower
)
```

```
for(i in 1:length(AdjMatrix[,1])){
  
  for(j in 1:length(AdjMatrix[1,])){
    
    if(Wmatpower2[i,j] != matpower2[i,j]){
      listAdj2$Apower2[i,j] = Wmatpower2[i,j]
    } else { listAdj2$Apower2[i,j] = NA }
    
    
    if(Wmatpower3[i,j] != matpower3[i,j]){
      listAdj2$Apower3[i,j] = Wmatpower3[i,j]
    } else { listAdj2$Apower3[i,j] = NA }
    
    
    if(Wmatpower4[i,j] != matpower4[i,j]){
      listAdj2$Apower4[i,j] = Wmatpower4[i,j]
    } else { listAdj2$Apower4[i,j] = NA }
    
    
    if(Wmatpower5[i,j] != matpower5[i,j]){
      listAdj2$Apower5[i,j] = Wmatpower5[i,j]
    } else { listAdj2$Apower5[i,j] = NA }
    
    
    if(Wmatpower6[i,j] != matpower6[i,j]){
      listAdj2$Apower6[i,j] = Wmatpower6[i,j]
    } else { listAdj2$Apower6[i,j] = NA }
    
    
    if(Wmatpower7[i,j] != matpower7[i,j]){
      listAdj2$Apower7[i,j] = Wmatpower7[i,j]
    } else { listAdj2$Apower7[i,j] = NA }
    
    
    if(Wmatpower8[i,j] != matpower8[i,j]){
      listAdj2$Apower8[i,j] = Wmatpower8[i,j]
    } else { listAdj2$Apower8[i,j] = NA }
    
    
    if(Wmatpower9[i,j] != matpower9[i,j]){
      listAdj2$Apower9[i,j] = Wmatpower9[i,j]
    } else { listAdj2$Apower9[i,j] = NA }
    
    
    if(Wmatpower10[i,j] != matpower10[i,j]){
      listAdj2$Apower10[i,j] = Wmatpower10[i,j]
    } else { listAdj2$Apower10[i,j] = NA }
    
    
    if(Wmatpower11[i,j] != matpower11[i,j]){
      listAdj2$Apower11[i,j] = Wmatpower11[i,j]
    } else { listAdj2$Apower11[i,j] = NA }
    
    
    if(Wmatpower12[i,j] != matpower12[i,j]){
      listAdj2$Apower12[i,j] = Wmatpower12[i,j]
    } else { listAdj2$Apower12[i,j] = NA }
    
    
    if(Wmatpower13[i,j] != matpower13[i,j]){
      listAdj2$Apower13[i,j] = Wmatpower13[i,j]
    } else { listAdj2$Apower13[i,j] = NA }
    
    
    if(Wmatpower14[i,j] != matpower14[i,j]){
      listAdj2$Apower14[i,j] = Wmatpower14[i,j]
    } else { listAdj2$Apower14[i,j] = NA }
    
    
    if(Wmatpower15[i,j] != matpower15[i,j]){
      listAdj2$Apower15[i,j] = Wmatpower15[i,j]
    } else { listAdj2$Apower15[i,j] = NA }
    
    
    if(Wmatpower16[i,j] != matpower16[i,j]){
      listAdj2$Apower16[i,j] = Wmatpower16[i,j]
    } else { listAdj2$Apower16[i,j] = NA }
    
    
    if(Wmatpower17[i,j] != matpower17[i,j]){
      listAdj2$Apower17[i,j] = Wmatpower17[i,j]
    } else { listAdj2$Apower17[i,j] = NA }
    
  }
  
}
```

\(~\)

Using the following code, the indices of the giant component edges in the **KEGG edgelist** are found, and they are stored at **commonIndex** object.

```
m = as_edgelist(weighted.giant.component)
m = as.data.frame(m , stringsAsFactors = F)
e = edgelist[,2:3]


index.of.rows.in.x.that.are.in.y  <- function(x,y)
{
  x.vec <- apply(x, 1, paste, collapse = "")
  y.vec <- apply(y, 1, paste, collapse = "")
  index = x.vec %in% y.vec
  return(which(index))
}
commonIndex = index.of.rows.in.x.that.are.in.y(e,m)
```

\(~\)  
\(~\)

##### 8.1.Computing the number of edges participating in MPFBL1, MNFBL2, MFFL1 and MNFFL2 multiple-edge subgraphs

\(~\)

The indices of edges taking part in MFFL1 subgraph are saved at **idxMFFL1** object through the following code.

```
MFFL1 = list()

idxMFFL1 = list()

for(i in commonIndex){

if(edgelist[i,4] == "1" & !(is.na(listAdj1$Apower2[edgelist[i,2],edgelist[i,3]]) |  listAdj1$Apower2[edgelist[i,2],edgelist[i,3]] == 0)){
  MFFL1[i]=listAdj1$Apower2[edgelist[i,2],edgelist[i,3]]
} else if(edgelist[i,4] == "1" & !(is.na(listAdj1$Apower3[edgelist[i,2],edgelist[i,3]]) |  listAdj1$Apower3[edgelist[i,2],edgelist[i,3]] == 0)){
  MFFL1[i]=listAdj1$Apower3[edgelist[i,2],edgelist[i,3]]
} else if(edgelist[i,4] == "1" & !(is.na(listAdj1$Apower4[edgelist[i,2],edgelist[i,3]]) |  listAdj1$Apower4[edgelist[i,2],edgelist[i,3]] == 0)){
  MFFL1[i]=listAdj1$Apower4[edgelist[i,2],edgelist[i,3]]
} else if(edgelist[i,4] == "1" & !(is.na(listAdj1$Apower5[edgelist[i,2],edgelist[i,3]]) |  listAdj1$Apower5[edgelist[i,2],edgelist[i,3]] == 0)){
  MFFL1[i]=listAdj1$Apower5[edgelist[i,2],edgelist[i,3]]
} else if(edgelist[i,4] == "1" & !(is.na(listAdj1$Apower6[edgelist[i,2],edgelist[i,3]]) |  listAdj1$Apower6[edgelist[i,2],edgelist[i,3]] == 0)){
  MFFL1[i]=listAdj1$Apower6[edgelist[i,2],edgelist[i,3]]
} else if(edgelist[i,4] == "1" & !(is.na(listAdj1$Apower7[edgelist[i,2],edgelist[i,3]]) |  listAdj1$Apower7[edgelist[i,2],edgelist[i,3]] == 0)){
  MFFL1[i]=listAdj1$Apower7[edgelist[i,2],edgelist[i,3]]
} else if(edgelist[i,4] == "1" & !(is.na(listAdj1$Apower8[edgelist[i,2],edgelist[i,3]]) |  listAdj1$Apower8[edgelist[i,2],edgelist[i,3]] == 0)){
  MFFL1[i]=listAdj1$Apower8[edgelist[i,2],edgelist[i,3]]
} else if(edgelist[i,4] == "1" & !(is.na(listAdj1$Apower9[edgelist[i,2],edgelist[i,3]]) |  listAdj1$Apower9[edgelist[i,2],edgelist[i,3]] == 0)){
  MFFL1[i]=listAdj1$Apower9[edgelist[i,2],edgelist[i,3]]
} else if(edgelist[i,4] == "1" & !(is.na(listAdj1$Apower10[edgelist[i,2],edgelist[i,3]]) |  listAdj1$Apower10[edgelist[i,2],edgelist[i,3]] == 0)){
  MFFL1[i]=listAdj1$Apower10[edgelist[i,2],edgelist[i,3]]
} else if(edgelist[i,4] == "1" & !(is.na(listAdj1$Apower11[edgelist[i,2],edgelist[i,3]]) |  listAdj1$Apower11[edgelist[i,2],edgelist[i,3]] == 0)){
  MFFL1[i]=listAdj1$Apower11[edgelist[i,2],edgelist[i,3]]
} else if(edgelist[i,4] == "1" & !(is.na(listAdj1$Apower12[edgelist[i,2],edgelist[i,3]]) |  listAdj1$Apower12[edgelist[i,2],edgelist[i,3]] == 0)){
  MFFL1[i]=listAdj1$Apower12[edgelist[i,2],edgelist[i,3]]
} else if(edgelist[i,4] == "1" & !(is.na(listAdj1$Apower13[edgelist[i,2],edgelist[i,3]]) |  listAdj1$Apower13[edgelist[i,2],edgelist[i,3]] == 0)){
  MFFL1[i]=listAdj1$Apower13[edgelist[i,2],edgelist[i,3]]
} else if(edgelist[i,4] == "1" & !(is.na(listAdj1$Apower14[edgelist[i,2],edgelist[i,3]]) |  listAdj1$Apower14[edgelist[i,2],edgelist[i,3]] == 0)){
  MFFL1[i]=listAdj1$Apower14[edgelist[i,2],edgelist[i,3]]
} else if(edgelist[i,4] == "1" & !(is.na(listAdj1$Apower15[edgelist[i,2],edgelist[i,3]]) |  listAdj1$Apower15[edgelist[i,2],edgelist[i,3]] == 0)){
  MFFL1[i]=listAdj1$Apower15[edgelist[i,2],edgelist[i,3]]
} else if(edgelist[i,4] == "1" & !(is.na(listAdj1$Apower16[edgelist[i,2],edgelist[i,3]]) |  listAdj1$Apower16[edgelist[i,2],edgelist[i,3]] == 0)){
  MFFL1[i]=listAdj1$Apower16[edgelist[i,2],edgelist[i,3]]
} else if(edgelist[i,4] == "1" & !(is.na(listAdj1$Apower17[edgelist[i,2],edgelist[i,3]]) |  listAdj1$Apower17[edgelist[i,2],edgelist[i,3]] == 0)){
  MFFL1[i]=listAdj1$Apower17[edgelist[i,2],edgelist[i,3]]
}

if( (edgelist[i,4] == "1" & !(is.na(listAdj1$Apower2[edgelist[i,2],edgelist[i,3]]) |  listAdj1$Apower2[edgelist[i,2],edgelist[i,3]] == 0)) | 
    (edgelist[i,4] == "1" & !(is.na(listAdj1$Apower3[edgelist[i,2],edgelist[i,3]]) |  listAdj1$Apower3[edgelist[i,2],edgelist[i,3]] == 0)) | 
    (edgelist[i,4] == "1" & !(is.na(listAdj1$Apower4[edgelist[i,2],edgelist[i,3]]) |  listAdj1$Apower4[edgelist[i,2],edgelist[i,3]] == 0)) |
    (edgelist[i,4] == "1" & !(is.na(listAdj1$Apower5[edgelist[i,2],edgelist[i,3]]) |  listAdj1$Apower5[edgelist[i,2],edgelist[i,3]] == 0)) |
    (edgelist[i,4] == "1" & !(is.na(listAdj1$Apower6[edgelist[i,2],edgelist[i,3]]) |  listAdj1$Apower6[edgelist[i,2],edgelist[i,3]] == 0)) |
    (edgelist[i,4] == "1" & !(is.na(listAdj1$Apower7[edgelist[i,2],edgelist[i,3]]) |  listAdj1$Apower7[edgelist[i,2],edgelist[i,3]] == 0)) | 
    (edgelist[i,4] == "1" & !(is.na(listAdj1$Apower8[edgelist[i,2],edgelist[i,3]]) |  listAdj1$Apower8[edgelist[i,2],edgelist[i,3]] == 0)) | 
    (edgelist[i,4] == "1" & !(is.na(listAdj1$Apower9[edgelist[i,2],edgelist[i,3]]) |  listAdj1$Apower9[edgelist[i,2],edgelist[i,3]] == 0)) |
    (edgelist[i,4] == "1" & !(is.na(listAdj1$Apower10[edgelist[i,2],edgelist[i,3]]) |  listAdj1$Apower10[edgelist[i,2],edgelist[i,3]] == 0)) |
    (edgelist[i,4] == "1" & !(is.na(listAdj1$Apower11[edgelist[i,2],edgelist[i,3]]) |  listAdj1$Apower11[edgelist[i,2],edgelist[i,3]] == 0)) |
    (edgelist[i,4] == "1" & !(is.na(listAdj1$Apower12[edgelist[i,2],edgelist[i,3]]) |  listAdj1$Apower12[edgelist[i,2],edgelist[i,3]] == 0)) | 
    (edgelist[i,4] == "1" & !(is.na(listAdj1$Apower13[edgelist[i,2],edgelist[i,3]]) |  listAdj1$Apower13[edgelist[i,2],edgelist[i,3]] == 0)) | 
    (edgelist[i,4] == "1" & !(is.na(listAdj1$Apower14[edgelist[i,2],edgelist[i,3]]) |  listAdj1$Apower14[edgelist[i,2],edgelist[i,3]] == 0)) |
    (edgelist[i,4] == "1" & !(is.na(listAdj1$Apower15[edgelist[i,2],edgelist[i,3]]) |  listAdj1$Apower15[edgelist[i,2],edgelist[i,3]] == 0)) |
    (edgelist[i,4] == "1" & !(is.na(listAdj1$Apower16[edgelist[i,2],edgelist[i,3]]) |  listAdj1$Apower16[edgelist[i,2],edgelist[i,3]] == 0)) |
    (edgelist[i,4] == "1" & !(is.na(listAdj1$Apower17[edgelist[i,2],edgelist[i,3]]) |  listAdj1$Apower17[edgelist[i,2],edgelist[i,3]] == 0))
){idxMFFL1[i] = i}
 
}

MFFL1 = unlist(MFFL1)

idxMFFL1 = unlist(idxMFFL1)
```

```
length(idxMFFL1)
```

```
## [1] 8425
```

\(~\)  
\(~\)

The indices of edges taking part in MPFBL1 subgraph are saved at **idxMPFBL1** object through the following code.

```
MPFBL1 = list()

idxMPFBL1 = list()
  
for(i in commonIndex){
  
  if(edgelist[i,4] == "1" & !(is.na(listAdj1$Apower2[edgelist[i,3],edgelist[i,2]]) |  listAdj1$Apower2[edgelist[i,3],edgelist[i,2]] == 0)){
    MPFBL1[i]=listAdj1$Apower2[edgelist[i,3],edgelist[i,2]]
  } else if(edgelist[i,4] == "1" & !(is.na(listAdj1$Apower3[edgelist[i,3],edgelist[i,2]]) |  listAdj1$Apower3[edgelist[i,3],edgelist[i,2]] == 0)){
    MPFBL1[i]=listAdj1$Apower3[edgelist[i,3],edgelist[i,2]]
  } else if(edgelist[i,4] == "1" & !(is.na(listAdj1$Apower4[edgelist[i,3],edgelist[i,2]]) |  listAdj1$Apower4[edgelist[i,3],edgelist[i,2]] == 0)){
    MPFBL1[i]=listAdj1$Apower4[edgelist[i,3],edgelist[i,2]]
  } else if(edgelist[i,4] == "1" & !(is.na(listAdj1$Apower5[edgelist[i,3],edgelist[i,2]]) |  listAdj1$Apower5[edgelist[i,3],edgelist[i,2]] == 0)){
    MPFBL1[i]=listAdj1$Apower5[edgelist[i,3],edgelist[i,2]]
  } else if(edgelist[i,4] == "1" & !(is.na(listAdj1$Apower6[edgelist[i,3],edgelist[i,2]]) |  listAdj1$Apower6[edgelist[i,3],edgelist[i,2]] == 0)){
    MPFBL1[i]=listAdj1$Apower6[edgelist[i,3],edgelist[i,2]]
  } else if(edgelist[i,4] == "1" & !(is.na(listAdj1$Apower7[edgelist[i,3],edgelist[i,2]]) |  listAdj1$Apower7[edgelist[i,3],edgelist[i,2]] == 0)){
    MPFBL1[i]=listAdj1$Apower7[edgelist[i,3],edgelist[i,2]]
  } else if(edgelist[i,4] == "1" & !(is.na(listAdj1$Apower8[edgelist[i,3],edgelist[i,2]]) |  listAdj1$Apower8[edgelist[i,3],edgelist[i,2]] == 0)){
    MPFBL1[i]=listAdj1$Apower8[edgelist[i,3],edgelist[i,2]]
  } else if(edgelist[i,4] == "1" & !(is.na(listAdj1$Apower9[edgelist[i,3],edgelist[i,2]]) |  listAdj1$Apower9[edgelist[i,3],edgelist[i,2]] == 0)){
    MPFBL1[i]=listAdj1$Apower9[edgelist[i,3],edgelist[i,2]]
  } else if(edgelist[i,4] == "1" & !(is.na(listAdj1$Apower10[edgelist[i,3],edgelist[i,2]]) |  listAdj1$Apower10[edgelist[i,3],edgelist[i,2]] == 0)){
    MPFBL1[i]=listAdj1$Apower10[edgelist[i,3],edgelist[i,2]]
  } else if(edgelist[i,4] == "1" & !(is.na(listAdj1$Apower11[edgelist[i,3],edgelist[i,2]]) |  listAdj1$Apower11[edgelist[i,3],edgelist[i,2]] == 0)){
    MPFBL1[i]=listAdj1$Apower11[edgelist[i,3],edgelist[i,2]]
  } else if(edgelist[i,4] == "1" & !(is.na(listAdj1$Apower12[edgelist[i,3],edgelist[i,2]]) |  listAdj1$Apower12[edgelist[i,3],edgelist[i,2]] == 0)){
    MPFBL1[i]=listAdj1$Apower12[edgelist[i,3],edgelist[i,2]]
  } else if(edgelist[i,4] == "1" & !(is.na(listAdj1$Apower13[edgelist[i,3],edgelist[i,2]]) |  listAdj1$Apower13[edgelist[i,3],edgelist[i,2]] == 0)){
    MPFBL1[i]=listAdj1$Apower13[edgelist[i,3],edgelist[i,2]]
  } else if(edgelist[i,4] == "1" & !(is.na(listAdj1$Apower14[edgelist[i,3],edgelist[i,2]]) |  listAdj1$Apower14[edgelist[i,3],edgelist[i,2]] == 0)){
    MPFBL1[i]=listAdj1$Apower14[edgelist[i,3],edgelist[i,2]]
  } else if(edgelist[i,4] == "1" & !(is.na(listAdj1$Apower15[edgelist[i,3],edgelist[i,2]]) |  listAdj1$Apower15[edgelist[i,3],edgelist[i,2]] == 0)){
    MPFBL1[i]=listAdj1$Apower15[edgelist[i,3],edgelist[i,2]]
  } else if(edgelist[i,4] == "1" & !(is.na(listAdj1$Apower16[edgelist[i,3],edgelist[i,2]]) |  listAdj1$Apower16[edgelist[i,3],edgelist[i,2]] == 0)){
    MPFBL1[i]=listAdj1$Apower16[edgelist[i,3],edgelist[i,2]]
  } else if(edgelist[i,4] == "1" & !(is.na(listAdj1$Apower17[edgelist[i,3],edgelist[i,2]]) |  listAdj1$Apower17[edgelist[i,3],edgelist[i,2]] == 0)){
    MPFBL1[i]=listAdj1$Apower17[edgelist[i,3],edgelist[i,2]]
  }
  
  if( (edgelist[i,4] == "1" & !(is.na(listAdj1$Apower2[edgelist[i,3],edgelist[i,2]]) |  listAdj1$Apower2[edgelist[i,3],edgelist[i,2]] == 0)) | 
      (edgelist[i,4] == "1" & !(is.na(listAdj1$Apower3[edgelist[i,3],edgelist[i,2]]) |  listAdj1$Apower3[edgelist[i,3],edgelist[i,2]] == 0)) | 
      (edgelist[i,4] == "1" & !(is.na(listAdj1$Apower4[edgelist[i,3],edgelist[i,2]]) |  listAdj1$Apower4[edgelist[i,3],edgelist[i,2]] == 0)) |
      (edgelist[i,4] == "1" & !(is.na(listAdj1$Apower5[edgelist[i,3],edgelist[i,2]]) |  listAdj1$Apower5[edgelist[i,3],edgelist[i,2]] == 0)) |
      (edgelist[i,4] == "1" & !(is.na(listAdj1$Apower6[edgelist[i,3],edgelist[i,2]]) |  listAdj1$Apower6[edgelist[i,3],edgelist[i,2]] == 0)) |
      (edgelist[i,4] == "1" & !(is.na(listAdj1$Apower7[edgelist[i,3],edgelist[i,2]]) |  listAdj1$Apower7[edgelist[i,3],edgelist[i,2]] == 0)) | 
      (edgelist[i,4] == "1" & !(is.na(listAdj1$Apower8[edgelist[i,3],edgelist[i,2]]) |  listAdj1$Apower8[edgelist[i,3],edgelist[i,2]] == 0)) | 
      (edgelist[i,4] == "1" & !(is.na(listAdj1$Apower9[edgelist[i,3],edgelist[i,2]]) |  listAdj1$Apower9[edgelist[i,3],edgelist[i,2]] == 0)) |
      (edgelist[i,4] == "1" & !(is.na(listAdj1$Apower10[edgelist[i,3],edgelist[i,2]]) |  listAdj1$Apower10[edgelist[i,3],edgelist[i,2]] == 0)) |
      (edgelist[i,4] == "1" & !(is.na(listAdj1$Apower11[edgelist[i,3],edgelist[i,2]]) |  listAdj1$Apower11[edgelist[i,3],edgelist[i,2]] == 0)) |
      (edgelist[i,4] == "1" & !(is.na(listAdj1$Apower12[edgelist[i,3],edgelist[i,2]]) |  listAdj1$Apower12[edgelist[i,3],edgelist[i,2]] == 0)) | 
      (edgelist[i,4] == "1" & !(is.na(listAdj1$Apower13[edgelist[i,3],edgelist[i,2]]) |  listAdj1$Apower13[edgelist[i,3],edgelist[i,2]] == 0)) | 
      (edgelist[i,4] == "1" & !(is.na(listAdj1$Apower14[edgelist[i,3],edgelist[i,2]]) |  listAdj1$Apower14[edgelist[i,3],edgelist[i,2]] == 0)) |
      (edgelist[i,4] == "1" & !(is.na(listAdj1$Apower15[edgelist[i,3],edgelist[i,2]]) |  listAdj1$Apower15[edgelist[i,3],edgelist[i,2]] == 0)) |
      (edgelist[i,4] == "1" & !(is.na(listAdj1$Apower16[edgelist[i,3],edgelist[i,2]]) |  listAdj1$Apower16[edgelist[i,3],edgelist[i,2]] == 0)) |
      (edgelist[i,4] == "1" & !(is.na(listAdj1$Apower17[edgelist[i,3],edgelist[i,2]]) |  listAdj1$Apower17[edgelist[i,3],edgelist[i,2]] == 0))
  ){idxMPFBL1[i] = i}
  
  
}


MPFBL1 = unlist(MPFBL1)

idxMPFBL1 = unlist(idxMPFBL1)
length(idxMPFBL1)
```

```
length(idxMPFBL1)
```

```
## [1] 2822
```

\(~\)  
\(~\)

The indices of edges taking part in MNFFL2 subgraph are stored at **idxMNFFL2** object through the following code.

```
MNFFL2 = list()

idxMNFFL2 = list()

for(i in commonIndex){
  
  if(edgelist[i,4] == "-1" & !(is.na(listAdj1$Apower2[edgelist[i,2],edgelist[i,3]]) |  listAdj1$Apower2[edgelist[i,2],edgelist[i,3]] == 0)){
    MNFFL2[i]=listAdj1$Apower2[edgelist[i,2],edgelist[i,3]]
  } else if(edgelist[i,4] == "-1" & !(is.na(listAdj1$Apower3[edgelist[i,2],edgelist[i,3]]) |  listAdj1$Apower3[edgelist[i,2],edgelist[i,3]] == 0)){
    MNFFL2[i]=listAdj1$Apower3[edgelist[i,2],edgelist[i,3]]
  } else if(edgelist[i,4] == "-1" & !(is.na(listAdj1$Apower4[edgelist[i,2],edgelist[i,3]]) |  listAdj1$Apower4[edgelist[i,2],edgelist[i,3]] == 0)){
    MNFFL2[i]=listAdj1$Apower4[edgelist[i,2],edgelist[i,3]]
  } else if(edgelist[i,4] == "-1" & !(is.na(listAdj1$Apower5[edgelist[i,2],edgelist[i,3]]) |  listAdj1$Apower5[edgelist[i,2],edgelist[i,3]] == 0)){
    MNFFL2[i]=listAdj1$Apower5[edgelist[i,2],edgelist[i,3]]
  } else if(edgelist[i,4] == "-1" & !(is.na(listAdj1$Apower6[edgelist[i,2],edgelist[i,3]]) |  listAdj1$Apower6[edgelist[i,2],edgelist[i,3]] == 0)){
    MNFFL2[i]=listAdj1$Apower6[edgelist[i,2],edgelist[i,3]]
  } else if(edgelist[i,4] == "-1" & !(is.na(listAdj1$Apower7[edgelist[i,2],edgelist[i,3]]) |  listAdj1$Apower7[edgelist[i,2],edgelist[i,3]] == 0)){
    MNFFL2[i]=listAdj1$Apower7[edgelist[i,2],edgelist[i,3]]
  } else if(edgelist[i,4] == "-1" & !(is.na(listAdj1$Apower8[edgelist[i,2],edgelist[i,3]]) |  listAdj1$Apower8[edgelist[i,2],edgelist[i,3]] == 0)){
    MNFFL2[i]=listAdj1$Apower8[edgelist[i,2],edgelist[i,3]]
  } else if(edgelist[i,4] == "-1" & !(is.na(listAdj1$Apower9[edgelist[i,2],edgelist[i,3]]) |  listAdj1$Apower9[edgelist[i,2],edgelist[i,3]] == 0)){
    MNFFL2[i]=listAdj1$Apower9[edgelist[i,2],edgelist[i,3]]
  } else if(edgelist[i,4] == "-1" & !(is.na(listAdj1$Apower10[edgelist[i,2],edgelist[i,3]]) |  listAdj1$Apower10[edgelist[i,2],edgelist[i,3]] == 0)){
    MNFFL2[i]=listAdj1$Apower10[edgelist[i,2],edgelist[i,3]]
  } else if(edgelist[i,4] == "-1" & !(is.na(listAdj1$Apower11[edgelist[i,2],edgelist[i,3]]) |  listAdj1$Apower11[edgelist[i,2],edgelist[i,3]] == 0)){
    MNFFL2[i]=listAdj1$Apower11[edgelist[i,2],edgelist[i,3]]
  } else if(edgelist[i,4] == "-1" & !(is.na(listAdj1$Apower12[edgelist[i,2],edgelist[i,3]]) |  listAdj1$Apower12[edgelist[i,2],edgelist[i,3]] == 0)){
    MNFFL2[i]=listAdj1$Apower12[edgelist[i,2],edgelist[i,3]]
  } else if(edgelist[i,4] == "-1" & !(is.na(listAdj1$Apower13[edgelist[i,2],edgelist[i,3]]) |  listAdj1$Apower13[edgelist[i,2],edgelist[i,3]] == 0)){
    MNFFL2[i]=listAdj1$Apower13[edgelist[i,2],edgelist[i,3]]
  } else if(edgelist[i,4] == "-1" & !(is.na(listAdj1$Apower14[edgelist[i,2],edgelist[i,3]]) |  listAdj1$Apower14[edgelist[i,2],edgelist[i,3]] == 0)){
    MNFFL2[i]=listAdj1$Apower14[edgelist[i,2],edgelist[i,3]]
  } else if(edgelist[i,4] == "-1" & !(is.na(listAdj1$Apower15[edgelist[i,2],edgelist[i,3]]) |  listAdj1$Apower15[edgelist[i,2],edgelist[i,3]] == 0)){
    MNFFL2[i]=listAdj1$Apower15[edgelist[i,2],edgelist[i,3]]
  } else if(edgelist[i,4] == "-1" & !(is.na(listAdj1$Apower16[edgelist[i,2],edgelist[i,3]]) |  listAdj1$Apower16[edgelist[i,2],edgelist[i,3]] == 0)){
    MNFFL2[i]=listAdj1$Apower16[edgelist[i,2],edgelist[i,3]]
  } else if(edgelist[i,4] == "-1" & !(is.na(listAdj1$Apower17[edgelist[i,2],edgelist[i,3]]) |  listAdj1$Apower17[edgelist[i,2],edgelist[i,3]] == 0)){
    MNFFL2[i]=listAdj1$Apower17[edgelist[i,2],edgelist[i,3]]
  }
  
  if( (edgelist[i,4] == "-1" & !(is.na(listAdj1$Apower2[edgelist[i,2],edgelist[i,3]]) |  listAdj1$Apower2[edgelist[i,2],edgelist[i,3]] == 0)) | 
      (edgelist[i,4] == "-1" & !(is.na(listAdj1$Apower3[edgelist[i,2],edgelist[i,3]]) |  listAdj1$Apower3[edgelist[i,2],edgelist[i,3]] == 0)) | 
      (edgelist[i,4] == "-1" & !(is.na(listAdj1$Apower4[edgelist[i,2],edgelist[i,3]]) |  listAdj1$Apower4[edgelist[i,2],edgelist[i,3]] == 0)) |
      (edgelist[i,4] == "-1" & !(is.na(listAdj1$Apower5[edgelist[i,2],edgelist[i,3]]) |  listAdj1$Apower5[edgelist[i,2],edgelist[i,3]] == 0)) |
      (edgelist[i,4] == "-1" & !(is.na(listAdj1$Apower6[edgelist[i,2],edgelist[i,3]]) |  listAdj1$Apower6[edgelist[i,2],edgelist[i,3]] == 0)) |
      (edgelist[i,4] == "-1" & !(is.na(listAdj1$Apower7[edgelist[i,2],edgelist[i,3]]) |  listAdj1$Apower7[edgelist[i,2],edgelist[i,3]] == 0)) | 
      (edgelist[i,4] == "-1" & !(is.na(listAdj1$Apower8[edgelist[i,2],edgelist[i,3]]) |  listAdj1$Apower8[edgelist[i,2],edgelist[i,3]] == 0)) | 
      (edgelist[i,4] == "-1" & !(is.na(listAdj1$Apower9[edgelist[i,2],edgelist[i,3]]) |  listAdj1$Apower9[edgelist[i,2],edgelist[i,3]] == 0)) |
      (edgelist[i,4] == "-1" & !(is.na(listAdj1$Apower10[edgelist[i,2],edgelist[i,3]]) |  listAdj1$Apower10[edgelist[i,2],edgelist[i,3]] == 0)) |
      (edgelist[i,4] == "-1" & !(is.na(listAdj1$Apower11[edgelist[i,2],edgelist[i,3]]) |  listAdj1$Apower11[edgelist[i,2],edgelist[i,3]] == 0)) |
      (edgelist[i,4] == "-1" & !(is.na(listAdj1$Apower12[edgelist[i,2],edgelist[i,3]]) |  listAdj1$Apower12[edgelist[i,2],edgelist[i,3]] == 0)) | 
      (edgelist[i,4] == "-1" & !(is.na(listAdj1$Apower13[edgelist[i,2],edgelist[i,3]]) |  listAdj1$Apower13[edgelist[i,2],edgelist[i,3]] == 0)) | 
      (edgelist[i,4] == "-1" & !(is.na(listAdj1$Apower14[edgelist[i,2],edgelist[i,3]]) |  listAdj1$Apower14[edgelist[i,2],edgelist[i,3]] == 0)) |
      (edgelist[i,4] == "-1" & !(is.na(listAdj1$Apower15[edgelist[i,2],edgelist[i,3]]) |  listAdj1$Apower15[edgelist[i,2],edgelist[i,3]] == 0)) |
      (edgelist[i,4] == "-1" & !(is.na(listAdj1$Apower16[edgelist[i,2],edgelist[i,3]]) |  listAdj1$Apower16[edgelist[i,2],edgelist[i,3]] == 0)) |
      (edgelist[i,4] == "-1" & !(is.na(listAdj1$Apower17[edgelist[i,2],edgelist[i,3]]) |  listAdj1$Apower17[edgelist[i,2],edgelist[i,3]] == 0))
  ){idxMNFFL2[i] = i}
  
  
}

MNFFL2 = unlist(MNFFL2)

idxMNFFL2 = unlist(idxMNFFL2)
```

```
length(idxMNFFL2)
```

```
## [1] 659
```

\(~\)  
\(~\)

The indices of edges taking part in MNFBL2 subgraph are stored at **idxMNFBL2** object through the following code.

```
MNFBL2 = list()

idxMNFBL2 = list()

for(i in commonIndex){
  
  if(edgelist[i,4] == "-1" & !(is.na(listAdj1$Apower2[edgelist[i,3],edgelist[i,2]]) |  listAdj1$Apower2[edgelist[i,3],edgelist[i,2]] == 0)){
    MNFBL2[i]=listAdj1$Apower2[edgelist[i,3],edgelist[i,2]]
  } else if(edgelist[i,4] == "-1" & !(is.na(listAdj1$Apower3[edgelist[i,3],edgelist[i,2]]) |  listAdj1$Apower3[edgelist[i,3],edgelist[i,2]] == 0)){
    MNFBL2[i]=listAdj1$Apower3[edgelist[i,3],edgelist[i,2]]
  } else if(edgelist[i,4] == "-1" & !(is.na(listAdj1$Apower4[edgelist[i,3],edgelist[i,2]]) |  listAdj1$Apower4[edgelist[i,3],edgelist[i,2]] == 0)){
    MNFBL2[i]=listAdj1$Apower4[edgelist[i,3],edgelist[i,2]]
  } else if(edgelist[i,4] == "-1" & !(is.na(listAdj1$Apower5[edgelist[i,3],edgelist[i,2]]) |  listAdj1$Apower5[edgelist[i,3],edgelist[i,2]] == 0)){
    MNFBL2[i]=listAdj1$Apower5[edgelist[i,3],edgelist[i,2]]
  } else if(edgelist[i,4] == "-1" & !(is.na(listAdj1$Apower6[edgelist[i,3],edgelist[i,2]]) |  listAdj1$Apower6[edgelist[i,3],edgelist[i,2]] == 0)){
    MNFBL2[i]=listAdj1$Apower6[edgelist[i,3],edgelist[i,2]]
  } else if(edgelist[i,4] == "-1" & !(is.na(listAdj1$Apower7[edgelist[i,3],edgelist[i,2]]) |  listAdj1$Apower7[edgelist[i,3],edgelist[i,2]] == 0)){
    MNFBL2[i]=listAdj1$Apower7[edgelist[i,3],edgelist[i,2]]
  } else if(edgelist[i,4] == "-1" & !(is.na(listAdj1$Apower8[edgelist[i,3],edgelist[i,2]]) |  listAdj1$Apower8[edgelist[i,3],edgelist[i,2]] == 0)){
    MNFBL2[i]=listAdj1$Apower8[edgelist[i,3],edgelist[i,2]]
  } else if(edgelist[i,4] == "-1" & !(is.na(listAdj1$Apower9[edgelist[i,3],edgelist[i,2]]) |  listAdj1$Apower9[edgelist[i,3],edgelist[i,2]] == 0)){
    MNFBL2[i]=listAdj1$Apower9[edgelist[i,3],edgelist[i,2]]
  } else if(edgelist[i,4] == "-1" & !(is.na(listAdj1$Apower10[edgelist[i,3],edgelist[i,2]]) |  listAdj1$Apower10[edgelist[i,3],edgelist[i,2]] == 0)){
    MNFBL2[i]=listAdj1$Apower10[edgelist[i,3],edgelist[i,2]]
  } else if(edgelist[i,4] == "-1" & !(is.na(listAdj1$Apower11[edgelist[i,3],edgelist[i,2]]) |  listAdj1$Apower11[edgelist[i,3],edgelist[i,2]] == 0)){
    MNFBL2[i]=listAdj1$Apower11[edgelist[i,3],edgelist[i,2]]
  } else if(edgelist[i,4] == "-1" & !(is.na(listAdj1$Apower12[edgelist[i,3],edgelist[i,2]]) |  listAdj1$Apower12[edgelist[i,3],edgelist[i,2]] == 0)){
    MNFBL2[i]=listAdj1$Apower12[edgelist[i,3],edgelist[i,2]]
  } else if(edgelist[i,4] == "-1" & !(is.na(listAdj1$Apower13[edgelist[i,3],edgelist[i,2]]) |  listAdj1$Apower13[edgelist[i,3],edgelist[i,2]] == 0)){
    MNFBL2[i]=listAdj1$Apower13[edgelist[i,3],edgelist[i,2]]
  } else if(edgelist[i,4] == "-1" & !(is.na(listAdj1$Apower14[edgelist[i,3],edgelist[i,2]]) |  listAdj1$Apower14[edgelist[i,3],edgelist[i,2]] == 0)){
    MNFBL2[i]=listAdj1$Apower14[edgelist[i,3],edgelist[i,2]]
  } else if(edgelist[i,4] == "-1" & !(is.na(listAdj1$Apower15[edgelist[i,3],edgelist[i,2]]) |  listAdj1$Apower15[edgelist[i,3],edgelist[i,2]] == 0)){
    MNFBL2[i]=listAdj1$Apower15[edgelist[i,3],edgelist[i,2]]
  } else if(edgelist[i,4] == "-1" & !(is.na(listAdj1$Apower16[edgelist[i,3],edgelist[i,2]]) |  listAdj1$Apower16[edgelist[i,3],edgelist[i,2]] == 0)){
    MNFBL2[i]=listAdj1$Apower16[edgelist[i,3],edgelist[i,2]]
  } else if(edgelist[i,4] == "-1" & !(is.na(listAdj1$Apower17[edgelist[i,3],edgelist[i,2]]) |  listAdj1$Apower17[edgelist[i,3],edgelist[i,2]] == 0)){
    MNFBL2[i]=listAdj1$Apower17[edgelist[i,3],edgelist[i,2]]
  }
  
  if( (edgelist[i,4] == "-1" & !(is.na(listAdj1$Apower2[edgelist[i,3],edgelist[i,2]]) |  listAdj1$Apower2[edgelist[i,3],edgelist[i,2]] == 0)) | 
      (edgelist[i,4] == "-1" & !(is.na(listAdj1$Apower3[edgelist[i,3],edgelist[i,2]]) |  listAdj1$Apower3[edgelist[i,3],edgelist[i,2]] == 0)) | 
      (edgelist[i,4] == "-1" & !(is.na(listAdj1$Apower4[edgelist[i,3],edgelist[i,2]]) |  listAdj1$Apower4[edgelist[i,3],edgelist[i,2]] == 0)) |
      (edgelist[i,4] == "-1" & !(is.na(listAdj1$Apower5[edgelist[i,3],edgelist[i,2]]) |  listAdj1$Apower5[edgelist[i,3],edgelist[i,2]] == 0)) |
      (edgelist[i,4] == "-1" & !(is.na(listAdj1$Apower6[edgelist[i,3],edgelist[i,2]]) |  listAdj1$Apower6[edgelist[i,3],edgelist[i,2]] == 0)) |
      (edgelist[i,4] == "-1" & !(is.na(listAdj1$Apower7[edgelist[i,3],edgelist[i,2]]) |  listAdj1$Apower7[edgelist[i,3],edgelist[i,2]] == 0)) | 
      (edgelist[i,4] == "-1" & !(is.na(listAdj1$Apower8[edgelist[i,3],edgelist[i,2]]) |  listAdj1$Apower8[edgelist[i,3],edgelist[i,2]] == 0)) | 
      (edgelist[i,4] == "-1" & !(is.na(listAdj1$Apower9[edgelist[i,3],edgelist[i,2]]) |  listAdj1$Apower9[edgelist[i,3],edgelist[i,2]] == 0)) |
      (edgelist[i,4] == "-1" & !(is.na(listAdj1$Apower10[edgelist[i,3],edgelist[i,2]]) |  listAdj1$Apower10[edgelist[i,3],edgelist[i,2]] == 0)) |
      (edgelist[i,4] == "-1" & !(is.na(listAdj1$Apower11[edgelist[i,3],edgelist[i,2]]) |  listAdj1$Apower11[edgelist[i,3],edgelist[i,2]] == 0)) |
      (edgelist[i,4] == "-1" & !(is.na(listAdj1$Apower12[edgelist[i,3],edgelist[i,2]]) |  listAdj1$Apower12[edgelist[i,3],edgelist[i,2]] == 0)) | 
      (edgelist[i,4] == "-1" & !(is.na(listAdj1$Apower13[edgelist[i,3],edgelist[i,2]]) |  listAdj1$Apower13[edgelist[i,3],edgelist[i,2]] == 0)) | 
      (edgelist[i,4] == "-1" & !(is.na(listAdj1$Apower14[edgelist[i,3],edgelist[i,2]]) |  listAdj1$Apower14[edgelist[i,3],edgelist[i,2]] == 0)) |
      (edgelist[i,4] == "-1" & !(is.na(listAdj1$Apower15[edgelist[i,3],edgelist[i,2]]) |  listAdj1$Apower15[edgelist[i,3],edgelist[i,2]] == 0)) |
      (edgelist[i,4] == "-1" & !(is.na(listAdj1$Apower16[edgelist[i,3],edgelist[i,2]]) |  listAdj1$Apower16[edgelist[i,3],edgelist[i,2]] == 0)) |
      (edgelist[i,4] == "-1" & !(is.na(listAdj1$Apower17[edgelist[i,3],edgelist[i,2]]) |  listAdj1$Apower17[edgelist[i,3],edgelist[i,2]] == 0))
  ){idxMNFBL2[i] = i}
  
}

MNFBL2 = unlist(MNFBL2)

idxMNFBL2 = unlist(idxMNFBL2)
```

```
length(idxMNFBL2)
```

```
## [1] 430
```

\(~\)  
\(~\)

Now it’s time to compute the number of edges involved in multiple-edge subgraphs (table 2).

\(~\)

##### MFFL1

```
MFFL1.pval.cor1 = 0
MFFL1.pval.cor.NA = 0
MFFL1.pval.cor2 = 0


for(i in idxMFFL1){
  logic =c()
  if(SignalingNet[[i]]$length > 1){
    
    for(j in 1:SignalingNet[[i]]$length){
      logic[j]= !any(unlist(lapply(SignalingNet[[i]]$corAnalysis[[j]],is.na)))
    }
    if(all(logic)){
      logic1 = c()
      logic2 = c()
      for(j in 1:SignalingNet[[i]]$length){
        
        logic1[j] = SignalingNet[[i]]$corAnalysis[[j]]$pearson$adjusted.pearsonpval < 0.05 & SignalingNet[[i]]$corAnalysis[[j]]$pearson$pearsoncor > 0 
        logic2[j] = SignalingNet[[i]]$corAnalysis[[j]]$pearson$adjusted.pearsonpval < 0.05 & SignalingNet[[i]]$corAnalysis[[j]]$pearson$pearsoncor < 0
      }
      if(all(logic1)) {MFFL1.pval.cor1=MFFL1.pval.cor1+1}
      if(all(logic2)) {MFFL1.pval.cor2=MFFL1.pval.cor2+1}
    }
  } else{
    if(!any(unlist(lapply(SignalingNet[[i]]$corAnalysis,is.na)))){
      a = SignalingNet[[i]]$corAnalysis$pearson$adjusted.pearsonpval < 0.05 & SignalingNet[[i]]$corAnalysis$pearson$pearsoncor > 0 
      b = SignalingNet[[i]]$corAnalysis$pearson$adjusted.pearsonpval < 0.05 & SignalingNet[[i]]$corAnalysis$pearson$pearsoncor < 0
      if(a){MFFL1.pval.cor1=MFFL1.pval.cor1+1}
      if(b){MFFL1.pval.cor2=MFFL1.pval.cor2+1}
    }
  }
}


for(i in idxMFFL1){
  logic = c()
  if(SignalingNet[[i]]$length > 1){
    
    for(j in 1:SignalingNet[[i]]$length){
      
      logic[j]=any(unlist(lapply(SignalingNet[[i]]$corAnalysis[[j]],is.na))) | SignalingNet[[i]]$corAnalysis[[j]]$pearson$adjusted.pearsonpval > 0.05
    }
    if(all(logic)){ MFFL1.pval.cor.NA = MFFL1.pval.cor.NA + 1 }
    
  }else{ if(any(unlist(lapply(SignalingNet[[i]]$corAnalysis,is.na)))| SignalingNet[[i]]$corAnalysis$pearson$adjusted.pearsonpval > 0.05){
    index.pval.cor.NA = MFFL1.pval.cor.NA + 1}
  }
}
```

```
MFFL1.pval.cor1
```

```
## [1] 537
```

```
MFFL1.pval.cor.NA
```

```
## [1] 1384
```

```
MFFL1.pval.cor2
```

```
## [1] 270
```

```
length(idxMFFL1) - (MFFL1.pval.cor1 + MFFL1.pval.cor.NA + MFFL1.pval.cor2)
```

```
## [1] 6234
```

\(~\)  
\(~\)

##### MNFBL2

```
MNFBL2.pval.cor1 = 0
MNFBL2.pval.cor.NA = 0
MNFBL2.pval.cor2 = 0


for(i in idxMNFBL2){
  logic =c()
  if(SignalingNet[[i]]$length > 1){
    
    for(j in 1:SignalingNet[[i]]$length){
      logic[j]= !any(unlist(lapply(SignalingNet[[i]]$corAnalysis[[j]],is.na)))
    }
    if(all(logic)){
      logic1 = c()
      logic2 = c()
      for(j in 1:SignalingNet[[i]]$length){
        
        logic1[j] = SignalingNet[[i]]$corAnalysis[[j]]$pearson$adjusted.pearsonpval < 0.05 & SignalingNet[[i]]$corAnalysis[[j]]$pearson$pearsoncor > 0 
        logic2[j] = SignalingNet[[i]]$corAnalysis[[j]]$pearson$adjusted.pearsonpval < 0.05 & SignalingNet[[i]]$corAnalysis[[j]]$pearson$pearsoncor < 0
      }
      if(all(logic1)) {MNFBL2.pval.cor1=MNFBL2.pval.cor1+1}
      if(all(logic2)) {MNFBL2.pval.cor2=MNFBL2.pval.cor2+1}
    }
  } else{
    if(!any(unlist(lapply(SignalingNet[[i]]$corAnalysis,is.na)))){
      a = SignalingNet[[i]]$corAnalysis$pearson$adjusted.pearsonpval < 0.05 & SignalingNet[[i]]$corAnalysis$pearson$pearsoncor > 0 
      b = SignalingNet[[i]]$corAnalysis$pearson$adjusted.pearsonpval < 0.05 & SignalingNet[[i]]$corAnalysis$pearson$pearsoncor < 0
      if(a){MNFBL2.pval.cor1=MNFBL2.pval.cor1+1}
      if(b){MNFBL2.pval.cor2=MNFBL2.pval.cor2+1}
    }
  }
}


for(i in idxMNFBL2){
  logic = c()
  if(SignalingNet[[i]]$length > 1){
    
    for(j in 1:SignalingNet[[i]]$length){
      
      logic[j]=any(unlist(lapply(SignalingNet[[i]]$corAnalysis[[j]],is.na))) | SignalingNet[[i]]$corAnalysis[[j]]$pearson$adjusted.pearsonpval > 0.05
    }
    if(all(logic)){ MNFBL2.pval.cor.NA = MNFBL2.pval.cor.NA + 1 }
    
  }else{ if(any(unlist(lapply(SignalingNet[[i]]$corAnalysis,is.na)))| SignalingNet[[i]]$corAnalysis$pearson$adjusted.pearsonpval > 0.05){
    index.pval.cor.NA = MNFBL2.pval.cor.NA + 1}
  }
}
```

```
MNFBL2.pval.cor1
```

```
## [1] 24
```

```
MNFBL2.pval.cor.NA
```

```
## [1] 83
```

```
MNFBL2.pval.cor2
```

```
## [1] 13
```

```
length(idxMNFBL2) - (MNFBL2.pval.cor1 + MNFBL2.pval.cor.NA + MNFBL2.pval.cor2)
```

```
## [1] 310
```

\(~\)  
\(~\)

##### MNFFL2

```
MNFFL2.pval.cor1 = 0
MNFFL2.pval.cor.NA = 0
MNFFL2.pval.cor2 = 0


for(i in idxMNFFL2){
  logic =c()
  if(SignalingNet[[i]]$length > 1){
    
    for(j in 1:SignalingNet[[i]]$length){
      logic[j]= !any(unlist(lapply(SignalingNet[[i]]$corAnalysis[[j]],is.na)))
    }
    if(all(logic)){
      logic1 = c()
      logic2 = c()
      for(j in 1:SignalingNet[[i]]$length){
        
        logic1[j] = SignalingNet[[i]]$corAnalysis[[j]]$pearson$adjusted.pearsonpval < 0.05 & SignalingNet[[i]]$corAnalysis[[j]]$pearson$pearsoncor > 0 
        logic2[j] = SignalingNet[[i]]$corAnalysis[[j]]$pearson$adjusted.pearsonpval < 0.05 & SignalingNet[[i]]$corAnalysis[[j]]$pearson$pearsoncor < 0
      }
      if(all(logic1)) {MNFFL2.pval.cor1=MNFFL2.pval.cor1+1}
      if(all(logic2)) {MNFFL2.pval.cor2=MNFFL2.pval.cor2+1}
    }
  } else{
    if(!any(unlist(lapply(SignalingNet[[i]]$corAnalysis,is.na)))){
      a = SignalingNet[[i]]$corAnalysis$pearson$adjusted.pearsonpval < 0.05 & SignalingNet[[i]]$corAnalysis$pearson$pearsoncor > 0 
      b = SignalingNet[[i]]$corAnalysis$pearson$adjusted.pearsonpval < 0.05 & SignalingNet[[i]]$corAnalysis$pearson$pearsoncor < 0
      if(a){MNFFL2.pval.cor1=MNFFL2.pval.cor1+1}
      if(b){MNFFL2.pval.cor2=MNFFL2.pval.cor2+1}
    }
  }
}


for(i in idxMNFFL2){
  logic = c()
  if(SignalingNet[[i]]$length > 1){
    
    for(j in 1:SignalingNet[[i]]$length){
      
      logic[j]=any(unlist(lapply(SignalingNet[[i]]$corAnalysis[[j]],is.na))) | SignalingNet[[i]]$corAnalysis[[j]]$pearson$adjusted.pearsonpval > 0.05
    }
    if(all(logic)){ MNFFL2.pval.cor.NA = MNFFL2.pval.cor.NA + 1 }
    
  }else{ if(any(unlist(lapply(SignalingNet[[i]]$corAnalysis,is.na)))| SignalingNet[[i]]$corAnalysis$pearson$adjusted.pearsonpval > 0.05){
    index.pval.cor.NA = MNFFL2.pval.cor.NA + 1}
  }
}
```

```
MNFFL2.pval.cor1
```

```
## [1] 43
```

```
MNFFL2.pval.cor.NA
```

```
## [1] 108
```

```
MNFFL2.pval.cor2
```

```
## [1] 27
```

```
length(idxMNFFL2) - (MNFFL2.pval.cor1 + MNFFL2.pval.cor.NA + MNFFL2.pval.cor2)
```

```
## [1] 481
```

\(~\)

###### MPFBL1

```
MPFBL1.pval.cor1 = 0
MPFBL1.pval.cor.NA = 0
MPFBL1.pval.cor2 = 0


for(i in idxMPFBL1){
  logic =c()
  if(SignalingNet[[i]]$length > 1){
    
    for(j in 1:SignalingNet[[i]]$length){
      logic[j]= !any(unlist(lapply(SignalingNet[[i]]$corAnalysis[[j]],is.na)))
    }
    if(all(logic)){
      logic1 = c()
      logic2 = c()
      for(j in 1:SignalingNet[[i]]$length){
        
        logic1[j] = SignalingNet[[i]]$corAnalysis[[j]]$pearson$adjusted.pearsonpval < 0.05 & SignalingNet[[i]]$corAnalysis[[j]]$pearson$pearsoncor > 0 
        logic2[j] = SignalingNet[[i]]$corAnalysis[[j]]$pearson$adjusted.pearsonpval < 0.05 & SignalingNet[[i]]$corAnalysis[[j]]$pearson$pearsoncor < 0
      }
      if(all(logic1)) {MPFBL1.pval.cor1=MPFBL1.pval.cor1+1}
      if(all(logic2)) {MPFBL1.pval.cor2=MPFBL1.pval.cor2+1}
    }
  } else{
    if(!any(unlist(lapply(SignalingNet[[i]]$corAnalysis,is.na)))){
      a = SignalingNet[[i]]$corAnalysis$pearson$adjusted.pearsonpval < 0.05 & SignalingNet[[i]]$corAnalysis$pearson$pearsoncor > 0 
      b = SignalingNet[[i]]$corAnalysis$pearson$adjusted.pearsonpval < 0.05 & SignalingNet[[i]]$corAnalysis$pearson$pearsoncor < 0
      if(a){MPFBL1.pval.cor1=MPFBL1.pval.cor1+1}
      if(b){MPFBL1.pval.cor2=MPFBL1.pval.cor2+1}
    }
  }
}


for(i in idxMPFBL1){
  logic = c()
  if(SignalingNet[[i]]$length > 1){
    
    for(j in 1:SignalingNet[[i]]$length){
      
      logic[j]=any(unlist(lapply(SignalingNet[[i]]$corAnalysis[[j]],is.na))) | SignalingNet[[i]]$corAnalysis[[j]]$pearson$adjusted.pearsonpval > 0.05
    }
    if(all(logic)){ MPFBL1.pval.cor.NA = MPFBL1.pval.cor.NA + 1 }
    
  }else{ if(any(unlist(lapply(SignalingNet[[i]]$corAnalysis,is.na)))| SignalingNet[[i]]$corAnalysis$pearson$adjusted.pearsonpval > 0.05){
    index.pval.cor.NA = MPFBL1.pval.cor.NA + 1}
  }
}
```

```
MPFBL1.pval.cor1
```

```
## [1] 224
```

```
MPFBL1.pval.cor.NA
```

```
## [1] 426
```

```
MPFBL1.pval.cor2
```

```
## [1] 85
```

```
length(idxMPFBL1) - (MPFBL1.pval.cor1 + MPFBL1.pval.cor.NA + MPFBL1.pval.cor2)
```

```
## [1] 2087
```

\(~\)  
\(~\)  
\(~\)  
\(~\)

##### 7.2.Computing the number of edges participating in MNFBL1, MPFBL2, MFFL2 and MNFFL1 multiple-edge subgraphs

\(~\)

The indices of edges which are involved in MNFBL1 subgraph are stored at **idxMNFBL1** object through the following code.

```
MNFBL1= list()

idxMNFBL1= list()

for(i in commonIndex){
  
  if(edgelist[i,4] == "1" & !(is.na(listAdj2$Apower2[edgelist[i,2],edgelist[i,3]]) |  listAdj2$Apower2[edgelist[i,2],edgelist[i,3]] == 0)){
    MNFBL1[i]=listAdj2$Apower2[edgelist[i,2],edgelist[i,3]]
  } else if(edgelist[i,4] == "1" & !(is.na(listAdj2$Apower3[edgelist[i,2],edgelist[i,3]]) |  listAdj2$Apower3[edgelist[i,2],edgelist[i,3]] == 0)){
    MNFBL1[i]=listAdj2$Apower3[edgelist[i,2],edgelist[i,3]]
  } else if(edgelist[i,4] == "1" & !(is.na(listAdj2$Apower4[edgelist[i,2],edgelist[i,3]]) |  listAdj2$Apower4[edgelist[i,2],edgelist[i,3]] == 0)){
    MNFBL1[i]=listAdj2$Apower4[edgelist[i,2],edgelist[i,3]]
  } else if(edgelist[i,4] == "1" & !(is.na(listAdj2$Apower5[edgelist[i,2],edgelist[i,3]]) |  listAdj2$Apower5[edgelist[i,2],edgelist[i,3]] == 0)){
    MNFBL1[i]=listAdj2$Apower5[edgelist[i,2],edgelist[i,3]]
  } else if(edgelist[i,4] == "1" & !(is.na(listAdj2$Apower6[edgelist[i,2],edgelist[i,3]]) |  listAdj2$Apower6[edgelist[i,2],edgelist[i,3]] == 0)){
    MNFBL1[i]=listAdj2$Apower6[edgelist[i,2],edgelist[i,3]]
  } else if(edgelist[i,4] == "1" & !(is.na(listAdj2$Apower7[edgelist[i,2],edgelist[i,3]]) |  listAdj2$Apower7[edgelist[i,2],edgelist[i,3]] == 0)){
    MNFBL1[i]=listAdj2$Apower7[edgelist[i,2],edgelist[i,3]]
  } else if(edgelist[i,4] == "1" & !(is.na(listAdj2$Apower8[edgelist[i,2],edgelist[i,3]]) |  listAdj2$Apower8[edgelist[i,2],edgelist[i,3]] == 0)){
    MNFBL1[i]=listAdj2$Apower8[edgelist[i,2],edgelist[i,3]]
  } else if(edgelist[i,4] == "1" & !(is.na(listAdj2$Apower9[edgelist[i,2],edgelist[i,3]]) |  listAdj2$Apower9[edgelist[i,2],edgelist[i,3]] == 0)){
    MNFBL1[i]=listAdj2$Apower9[edgelist[i,2],edgelist[i,3]]
  } else if(edgelist[i,4] == "1" & !(is.na(listAdj2$Apower10[edgelist[i,2],edgelist[i,3]]) |  listAdj2$Apower10[edgelist[i,2],edgelist[i,3]] == 0)){
    MNFBL1[i]=listAdj2$Apower10[edgelist[i,2],edgelist[i,3]]
  } else if(edgelist[i,4] == "1" & !(is.na(listAdj2$Apower11[edgelist[i,2],edgelist[i,3]]) |  listAdj2$Apower11[edgelist[i,2],edgelist[i,3]] == 0)){
    MNFBL1[i]=listAdj2$Apower11[edgelist[i,2],edgelist[i,3]]
  } else if(edgelist[i,4] == "1" & !(is.na(listAdj2$Apower12[edgelist[i,2],edgelist[i,3]]) |  listAdj2$Apower12[edgelist[i,2],edgelist[i,3]] == 0)){
    MNFBL1[i]=listAdj2$Apower12[edgelist[i,2],edgelist[i,3]]
  } else if(edgelist[i,4] == "1" & !(is.na(listAdj2$Apower13[edgelist[i,2],edgelist[i,3]]) |  listAdj2$Apower13[edgelist[i,2],edgelist[i,3]] == 0)){
    MNFBL1[i]=listAdj2$Apower13[edgelist[i,2],edgelist[i,3]]
  } else if(edgelist[i,4] == "1" & !(is.na(listAdj2$Apower14[edgelist[i,2],edgelist[i,3]]) |  listAdj2$Apower14[edgelist[i,2],edgelist[i,3]] == 0)){
    MNFBL1[i]=listAdj2$Apower14[edgelist[i,2],edgelist[i,3]]
  } else if(edgelist[i,4] == "1" & !(is.na(listAdj2$Apower15[edgelist[i,2],edgelist[i,3]]) |  listAdj2$Apower15[edgelist[i,2],edgelist[i,3]] == 0)){
    MNFBL1[i]=listAdj2$Apower15[edgelist[i,2],edgelist[i,3]]
  } else if(edgelist[i,4] == "1" & !(is.na(listAdj2$Apower16[edgelist[i,2],edgelist[i,3]]) |  listAdj2$Apower16[edgelist[i,2],edgelist[i,3]] == 0)){
    MNFBL1[i]=listAdj2$Apower16[edgelist[i,2],edgelist[i,3]]
  } else if(edgelist[i,4] == "1" & !(is.na(listAdj2$Apower17[edgelist[i,2],edgelist[i,3]]) |  listAdj2$Apower17[edgelist[i,2],edgelist[i,3]] == 0)){
    MNFBL1[i]=listAdj2$Apower17[edgelist[i,2],edgelist[i,3]]
  }
  
  if( (edgelist[i,4] == "1" & !(is.na(listAdj2$Apower2[edgelist[i,2],edgelist[i,3]]) |  listAdj2$Apower2[edgelist[i,2],edgelist[i,3]] == 0)) | 
      (edgelist[i,4] == "1" & !(is.na(listAdj2$Apower3[edgelist[i,2],edgelist[i,3]]) |  listAdj2$Apower3[edgelist[i,2],edgelist[i,3]] == 0)) | 
      (edgelist[i,4] == "1" & !(is.na(listAdj2$Apower4[edgelist[i,2],edgelist[i,3]]) |  listAdj2$Apower4[edgelist[i,2],edgelist[i,3]] == 0)) |
      (edgelist[i,4] == "1" & !(is.na(listAdj2$Apower5[edgelist[i,2],edgelist[i,3]]) |  listAdj2$Apower5[edgelist[i,2],edgelist[i,3]] == 0)) |
      (edgelist[i,4] == "1" & !(is.na(listAdj2$Apower6[edgelist[i,2],edgelist[i,3]]) |  listAdj2$Apower6[edgelist[i,2],edgelist[i,3]] == 0)) |
      (edgelist[i,4] == "1" & !(is.na(listAdj2$Apower7[edgelist[i,2],edgelist[i,3]]) |  listAdj2$Apower7[edgelist[i,2],edgelist[i,3]] == 0)) | 
      (edgelist[i,4] == "1" & !(is.na(listAdj2$Apower8[edgelist[i,2],edgelist[i,3]]) |  listAdj2$Apower8[edgelist[i,2],edgelist[i,3]] == 0)) | 
      (edgelist[i,4] == "1" & !(is.na(listAdj2$Apower9[edgelist[i,2],edgelist[i,3]]) |  listAdj2$Apower9[edgelist[i,2],edgelist[i,3]] == 0)) |
      (edgelist[i,4] == "1" & !(is.na(listAdj2$Apower10[edgelist[i,2],edgelist[i,3]]) |  listAdj2$Apower10[edgelist[i,2],edgelist[i,3]] == 0)) |
      (edgelist[i,4] == "1" & !(is.na(listAdj2$Apower11[edgelist[i,2],edgelist[i,3]]) |  listAdj2$Apower11[edgelist[i,2],edgelist[i,3]] == 0)) |
      (edgelist[i,4] == "1" & !(is.na(listAdj2$Apower12[edgelist[i,2],edgelist[i,3]]) |  listAdj2$Apower12[edgelist[i,2],edgelist[i,3]] == 0)) | 
      (edgelist[i,4] == "1" & !(is.na(listAdj2$Apower13[edgelist[i,2],edgelist[i,3]]) |  listAdj2$Apower13[edgelist[i,2],edgelist[i,3]] == 0)) | 
      (edgelist[i,4] == "1" & !(is.na(listAdj2$Apower14[edgelist[i,2],edgelist[i,3]]) |  listAdj2$Apower14[edgelist[i,2],edgelist[i,3]] == 0)) |
      (edgelist[i,4] == "1" & !(is.na(listAdj2$Apower15[edgelist[i,2],edgelist[i,3]]) |  listAdj2$Apower15[edgelist[i,2],edgelist[i,3]] == 0)) |
      (edgelist[i,4] == "1" & !(is.na(listAdj2$Apower16[edgelist[i,2],edgelist[i,3]]) |  listAdj2$Apower16[edgelist[i,2],edgelist[i,3]] == 0)) |
      (edgelist[i,4] == "1" & !(is.na(listAdj2$Apower17[edgelist[i,2],edgelist[i,3]]) |  listAdj2$Apower17[edgelist[i,2],edgelist[i,3]] == 0))
  ){idxMNFBL1[i] = i}
  
}

MNFBL1 = unlist(MNFBL1)

idxMNFBL1 = unlist(idxMNFBL1)
```

```
length(idxMNFBL1)
```

```
## [1] 11543
```

\(~\)  
\(~\)

The indices of edges involved in MPFBL2 subgraph are stored at **idxMPFBL2** object through the following code.

```
MPFBL2= list()

idxMPFBL2 = list()

for(i in commonIndex){
  
  if(edgelist[i,4] == "-1" & !(is.na(listAdj2$Apower2[edgelist[i,3],edgelist[i,2]]) |  listAdj2$Apower2[edgelist[i,3],edgelist[i,2]] == 0)){
    MPFBL2[i]=listAdj2$Apower2[edgelist[i,3],edgelist[i,2]]
  } else if(edgelist[i,4] == "-1" & !(is.na(listAdj2$Apower3[edgelist[i,3],edgelist[i,2]]) |  listAdj2$Apower3[edgelist[i,3],edgelist[i,2]] == 0)){
    MPFBL2[i]=listAdj2$Apower3[edgelist[i,3],edgelist[i,2]]
  } else if(edgelist[i,4] == "-1" & !(is.na(listAdj2$Apower4[edgelist[i,3],edgelist[i,2]]) |  listAdj2$Apower4[edgelist[i,3],edgelist[i,2]] == 0)){
    MPFBL2[i]=listAdj2$Apower4[edgelist[i,3],edgelist[i,2]]
  } else if(edgelist[i,4] == "-1" & !(is.na(listAdj2$Apower5[edgelist[i,3],edgelist[i,2]]) |  listAdj2$Apower5[edgelist[i,3],edgelist[i,2]] == 0)){
    MPFBL2[i]=listAdj2$Apower5[edgelist[i,3],edgelist[i,2]]
  } else if(edgelist[i,4] == "-1" & !(is.na(listAdj2$Apower6[edgelist[i,3],edgelist[i,2]]) |  listAdj2$Apower6[edgelist[i,3],edgelist[i,2]] == 0)){
    MPFBL2[i]=listAdj2$Apower6[edgelist[i,3],edgelist[i,2]]
  } else if(edgelist[i,4] == "-1" & !(is.na(listAdj2$Apower7[edgelist[i,3],edgelist[i,2]]) |  listAdj2$Apower7[edgelist[i,3],edgelist[i,2]] == 0)){
    MPFBL2[i]=listAdj2$Apower7[edgelist[i,3],edgelist[i,2]]
  } else if(edgelist[i,4] == "-1" & !(is.na(listAdj2$Apower8[edgelist[i,3],edgelist[i,2]]) |  listAdj2$Apower8[edgelist[i,3],edgelist[i,2]] == 0)){
    MPFBL2[i]=listAdj2$Apower8[edgelist[i,3],edgelist[i,2]]
  } else if(edgelist[i,4] == "-1" & !(is.na(listAdj2$Apower9[edgelist[i,3],edgelist[i,2]]) |  listAdj2$Apower9[edgelist[i,3],edgelist[i,2]] == 0)){
    MPFBL2[i]=listAdj2$Apower9[edgelist[i,3],edgelist[i,2]]
  } else if(edgelist[i,4] == "-1" & !(is.na(listAdj2$Apower10[edgelist[i,3],edgelist[i,2]]) |  listAdj2$Apower10[edgelist[i,3],edgelist[i,2]] == 0)){
    MPFBL2[i]=listAdj2$Apower10[edgelist[i,3],edgelist[i,2]]
  } else if(edgelist[i,4] == "-1" & !(is.na(listAdj2$Apower11[edgelist[i,3],edgelist[i,2]]) |  listAdj2$Apower11[edgelist[i,3],edgelist[i,2]] == 0)){
    MPFBL2[i]=listAdj2$Apower11[edgelist[i,3],edgelist[i,2]]
  } else if(edgelist[i,4] == "-1" & !(is.na(listAdj2$Apower12[edgelist[i,3],edgelist[i,2]]) |  listAdj2$Apower12[edgelist[i,3],edgelist[i,2]] == 0)){
    MPFBL2[i]=listAdj2$Apower12[edgelist[i,3],edgelist[i,2]]
  } else if(edgelist[i,4] == "-1" & !(is.na(listAdj2$Apower13[edgelist[i,3],edgelist[i,2]]) |  listAdj2$Apower13[edgelist[i,3],edgelist[i,2]] == 0)){
    MPFBL2[i]=listAdj2$Apower13[edgelist[i,3],edgelist[i,2]]
  } else if(edgelist[i,4] == "-1" & !(is.na(listAdj2$Apower14[edgelist[i,3],edgelist[i,2]]) |  listAdj2$Apower14[edgelist[i,3],edgelist[i,2]] == 0)){
    MPFBL2[i]=listAdj2$Apower14[edgelist[i,3],edgelist[i,2]]
  } else if(edgelist[i,4] == "-1" & !(is.na(listAdj2$Apower15[edgelist[i,3],edgelist[i,2]]) |  listAdj2$Apower15[edgelist[i,3],edgelist[i,2]] == 0)){
    MPFBL2[i]=listAdj2$Apower15[edgelist[i,3],edgelist[i,2]]
  } else if(edgelist[i,4] == "-1" & !(is.na(listAdj2$Apower16[edgelist[i,3],edgelist[i,2]]) |  listAdj2$Apower16[edgelist[i,3],edgelist[i,2]] == 0)){
    MPFBL2[i]=listAdj2$Apower16[edgelist[i,3],edgelist[i,2]]
  } else if(edgelist[i,4] == "-1" & !(is.na(listAdj2$Apower17[edgelist[i,3],edgelist[i,2]]) |  listAdj2$Apower17[edgelist[i,3],edgelist[i,2]] == 0)){
    MPFBL2[i]=listAdj2$Apower17[edgelist[i,3],edgelist[i,2]]
  }
  
  if( (edgelist[i,4] == "-1" & !(is.na(listAdj2$Apower2[edgelist[i,3],edgelist[i,2]]) |  listAdj2$Apower2[edgelist[i,3],edgelist[i,2]] == 0)) | 
      (edgelist[i,4] == "-1" & !(is.na(listAdj2$Apower3[edgelist[i,3],edgelist[i,2]]) |  listAdj2$Apower3[edgelist[i,3],edgelist[i,2]] == 0)) | 
      (edgelist[i,4] == "-1" & !(is.na(listAdj2$Apower4[edgelist[i,3],edgelist[i,2]]) |  listAdj2$Apower4[edgelist[i,3],edgelist[i,2]] == 0)) |
      (edgelist[i,4] == "-1" & !(is.na(listAdj2$Apower5[edgelist[i,3],edgelist[i,2]]) |  listAdj2$Apower5[edgelist[i,3],edgelist[i,2]] == 0)) |
      (edgelist[i,4] == "-1" & !(is.na(listAdj2$Apower6[edgelist[i,3],edgelist[i,2]]) |  listAdj2$Apower6[edgelist[i,3],edgelist[i,2]] == 0)) |
      (edgelist[i,4] == "-1" & !(is.na(listAdj2$Apower7[edgelist[i,3],edgelist[i,2]]) |  listAdj2$Apower7[edgelist[i,3],edgelist[i,2]] == 0)) | 
      (edgelist[i,4] == "-1" & !(is.na(listAdj2$Apower8[edgelist[i,3],edgelist[i,2]]) |  listAdj2$Apower8[edgelist[i,3],edgelist[i,2]] == 0)) | 
      (edgelist[i,4] == "-1" & !(is.na(listAdj2$Apower9[edgelist[i,3],edgelist[i,2]]) |  listAdj2$Apower9[edgelist[i,3],edgelist[i,2]] == 0)) |
      (edgelist[i,4] == "-1" & !(is.na(listAdj2$Apower10[edgelist[i,3],edgelist[i,2]]) |  listAdj2$Apower10[edgelist[i,3],edgelist[i,2]] == 0)) |
      (edgelist[i,4] == "-1" & !(is.na(listAdj2$Apower11[edgelist[i,3],edgelist[i,2]]) |  listAdj2$Apower11[edgelist[i,3],edgelist[i,2]] == 0)) |
      (edgelist[i,4] == "-1" & !(is.na(listAdj2$Apower12[edgelist[i,3],edgelist[i,2]]) |  listAdj2$Apower12[edgelist[i,3],edgelist[i,2]] == 0)) | 
      (edgelist[i,4] == "-1" & !(is.na(listAdj2$Apower13[edgelist[i,3],edgelist[i,2]]) |  listAdj2$Apower13[edgelist[i,3],edgelist[i,2]] == 0)) | 
      (edgelist[i,4] == "-1" & !(is.na(listAdj2$Apower14[edgelist[i,3],edgelist[i,2]]) |  listAdj2$Apower14[edgelist[i,3],edgelist[i,2]] == 0)) |
      (edgelist[i,4] == "-1" & !(is.na(listAdj2$Apower15[edgelist[i,3],edgelist[i,2]]) |  listAdj2$Apower15[edgelist[i,3],edgelist[i,2]] == 0)) |
      (edgelist[i,4] == "-1" & !(is.na(listAdj2$Apower16[edgelist[i,3],edgelist[i,2]]) |  listAdj2$Apower16[edgelist[i,3],edgelist[i,2]] == 0)) |
      (edgelist[i,4] == "-1" & !(is.na(listAdj2$Apower17[edgelist[i,3],edgelist[i,2]]) |  listAdj2$Apower17[edgelist[i,3],edgelist[i,2]] == 0))
  ){idxMPFBL2[i] = i}
  
}


MPFBL2 = unlist(MPFBL2)

idxMPFBL2 = unlist(idxMPFBL2)
```

```
length(idxMPFBL2)
```

```
## [1] 974
```

\(~\)  
\(~\)

The indices of edges involved in MFFL2 subgraph are stored at **idxMFFL2** object through the following code.

```
MFFL2 = list()

idxMFFL2 = list()

for(i in commonIndex){
  
  if(edgelist[i,4] == "-1" & !(is.na(listAdj2$Apower2[edgelist[i,2],edgelist[i,3]]) |  listAdj2$Apower2[edgelist[i,2],edgelist[i,3]] == 0)){
    MFFL2[i]=listAdj2$Apower2[edgelist[i,2],edgelist[i,3]]
  } else if(edgelist[i,4] == "-1" & !(is.na(listAdj2$Apower3[edgelist[i,2],edgelist[i,3]]) |  listAdj2$Apower3[edgelist[i,2],edgelist[i,3]] == 0)){
    MFFL2[i]=listAdj2$Apower3[edgelist[i,2],edgelist[i,3]]
  } else if(edgelist[i,4] == "-1" & !(is.na(listAdj2$Apower4[edgelist[i,2],edgelist[i,3]]) |  listAdj2$Apower4[edgelist[i,2],edgelist[i,3]] == 0)){
    MFFL2[i]=listAdj2$Apower4[edgelist[i,2],edgelist[i,3]]
  } else if(edgelist[i,4] == "-1" & !(is.na(listAdj2$Apower5[edgelist[i,2],edgelist[i,3]]) |  listAdj2$Apower5[edgelist[i,2],edgelist[i,3]] == 0)){
    MFFL2[i]=listAdj2$Apower5[edgelist[i,2],edgelist[i,3]]
  } else if(edgelist[i,4] == "-1" & !(is.na(listAdj2$Apower6[edgelist[i,2],edgelist[i,3]]) |  listAdj2$Apower6[edgelist[i,2],edgelist[i,3]] == 0)){
    MFFL2[i]=listAdj2$Apower6[edgelist[i,2],edgelist[i,3]]
  } else if(edgelist[i,4] == "-1" & !(is.na(listAdj2$Apower7[edgelist[i,2],edgelist[i,3]]) |  listAdj2$Apower7[edgelist[i,2],edgelist[i,3]] == 0)){
    MFFL2[i]=listAdj2$Apower7[edgelist[i,2],edgelist[i,3]]
  } else if(edgelist[i,4] == "-1" & !(is.na(listAdj2$Apower8[edgelist[i,2],edgelist[i,3]]) |  listAdj2$Apower8[edgelist[i,2],edgelist[i,3]] == 0)){
    MFFL2[i]=listAdj2$Apower8[edgelist[i,2],edgelist[i,3]]
  } else if(edgelist[i,4] == "-1" & !(is.na(listAdj2$Apower9[edgelist[i,2],edgelist[i,3]]) |  listAdj2$Apower9[edgelist[i,2],edgelist[i,3]] == 0)){
    MFFL2[i]=listAdj2$Apower9[edgelist[i,2],edgelist[i,3]]
  } else if(edgelist[i,4] == "-1" & !(is.na(listAdj2$Apower10[edgelist[i,2],edgelist[i,3]]) |  listAdj2$Apower10[edgelist[i,2],edgelist[i,3]] == 0)){
    MFFL2[i]=listAdj2$Apower10[edgelist[i,2],edgelist[i,3]]
  } else if(edgelist[i,4] == "-1" & !(is.na(listAdj2$Apower11[edgelist[i,2],edgelist[i,3]]) |  listAdj2$Apower11[edgelist[i,2],edgelist[i,3]] == 0)){
    MFFL2[i]=listAdj2$Apower11[edgelist[i,2],edgelist[i,3]]
  } else if(edgelist[i,4] == "-1" & !(is.na(listAdj2$Apower12[edgelist[i,2],edgelist[i,3]]) |  listAdj2$Apower12[edgelist[i,2],edgelist[i,3]] == 0)){
    MFFL2[i]=listAdj2$Apower12[edgelist[i,2],edgelist[i,3]]
  } else if(edgelist[i,4] == "-1" & !(is.na(listAdj2$Apower13[edgelist[i,2],edgelist[i,3]]) |  listAdj2$Apower13[edgelist[i,2],edgelist[i,3]] == 0)){
    MFFL2[i]=listAdj2$Apower13[edgelist[i,2],edgelist[i,3]]
  } else if(edgelist[i,4] == "-1" & !(is.na(listAdj2$Apower14[edgelist[i,2],edgelist[i,3]]) |  listAdj2$Apower14[edgelist[i,2],edgelist[i,3]] == 0)){
    MFFL2[i]=listAdj2$Apower14[edgelist[i,2],edgelist[i,3]]
  } else if(edgelist[i,4] == "-1" & !(is.na(listAdj2$Apower15[edgelist[i,2],edgelist[i,3]]) |  listAdj2$Apower15[edgelist[i,2],edgelist[i,3]] == 0)){
    MFFL2[i]=listAdj2$Apower15[edgelist[i,2],edgelist[i,3]]
  } else if(edgelist[i,4] == "-1" & !(is.na(listAdj2$Apower16[edgelist[i,2],edgelist[i,3]]) |  listAdj2$Apower16[edgelist[i,2],edgelist[i,3]] == 0)){
    MFFL2[i]=listAdj2$Apower16[edgelist[i,2],edgelist[i,3]]
  } else if(edgelist[i,4] == "-1" & !(is.na(listAdj2$Apower17[edgelist[i,2],edgelist[i,3]]) |  listAdj2$Apower17[edgelist[i,2],edgelist[i,3]] == 0)){
    MFFL2[i]=listAdj2$Apower17[edgelist[i,2],edgelist[i,3]]
  }
  
  if( (edgelist[i,4] == "-1" & !(is.na(listAdj2$Apower2[edgelist[i,2],edgelist[i,3]]) |  listAdj2$Apower2[edgelist[i,2],edgelist[i,3]] == 0)) | 
      (edgelist[i,4] == "-1" & !(is.na(listAdj2$Apower3[edgelist[i,2],edgelist[i,3]]) |  listAdj2$Apower3[edgelist[i,2],edgelist[i,3]] == 0)) | 
      (edgelist[i,4] == "-1" & !(is.na(listAdj2$Apower4[edgelist[i,2],edgelist[i,3]]) |  listAdj2$Apower4[edgelist[i,2],edgelist[i,3]] == 0)) |
      (edgelist[i,4] == "-1" & !(is.na(listAdj2$Apower5[edgelist[i,2],edgelist[i,3]]) |  listAdj2$Apower5[edgelist[i,2],edgelist[i,3]] == 0)) |
      (edgelist[i,4] == "-1" & !(is.na(listAdj2$Apower6[edgelist[i,2],edgelist[i,3]]) |  listAdj2$Apower6[edgelist[i,2],edgelist[i,3]] == 0)) |
      (edgelist[i,4] == "-1" & !(is.na(listAdj2$Apower7[edgelist[i,2],edgelist[i,3]]) |  listAdj2$Apower7[edgelist[i,2],edgelist[i,3]] == 0)) | 
      (edgelist[i,4] == "-1" & !(is.na(listAdj2$Apower8[edgelist[i,2],edgelist[i,3]]) |  listAdj2$Apower8[edgelist[i,2],edgelist[i,3]] == 0)) | 
      (edgelist[i,4] == "-1" & !(is.na(listAdj2$Apower9[edgelist[i,2],edgelist[i,3]]) |  listAdj2$Apower9[edgelist[i,2],edgelist[i,3]] == 0)) |
      (edgelist[i,4] == "-1" & !(is.na(listAdj2$Apower10[edgelist[i,2],edgelist[i,3]]) |  listAdj2$Apower10[edgelist[i,2],edgelist[i,3]] == 0)) |
      (edgelist[i,4] == "-1" & !(is.na(listAdj2$Apower11[edgelist[i,2],edgelist[i,3]]) |  listAdj2$Apower11[edgelist[i,2],edgelist[i,3]] == 0)) |
      (edgelist[i,4] == "-1" & !(is.na(listAdj2$Apower12[edgelist[i,2],edgelist[i,3]]) |  listAdj2$Apower12[edgelist[i,2],edgelist[i,3]] == 0)) | 
      (edgelist[i,4] == "-1" & !(is.na(listAdj2$Apower13[edgelist[i,2],edgelist[i,3]]) |  listAdj2$Apower13[edgelist[i,2],edgelist[i,3]] == 0)) | 
      (edgelist[i,4] == "-1" & !(is.na(listAdj2$Apower14[edgelist[i,2],edgelist[i,3]]) |  listAdj2$Apower14[edgelist[i,2],edgelist[i,3]] == 0)) |
      (edgelist[i,4] == "-1" & !(is.na(listAdj2$Apower15[edgelist[i,2],edgelist[i,3]]) |  listAdj2$Apower15[edgelist[i,2],edgelist[i,3]] == 0)) |
      (edgelist[i,4] == "-1" & !(is.na(listAdj2$Apower16[edgelist[i,2],edgelist[i,3]]) |  listAdj2$Apower16[edgelist[i,2],edgelist[i,3]] == 0)) |
      (edgelist[i,4] == "-1" & !(is.na(listAdj2$Apower17[edgelist[i,2],edgelist[i,3]]) |  listAdj2$Apower17[edgelist[i,2],edgelist[i,3]] == 0))
  ){idxMFFL2[i] = i}
  
}

MFFL2 = unlist(MFFL2)

idxMFFL2 = unlist(idxMFFL2)
```

```
length(idxMFFL2)
```

```
## [1] 2609
```

\(~\)  
\(~\)

The indices of edges involved in MNFFL1 subgraph are stored at **MNFFL1** object through the following code.

```
MNFFL1 = list()

idxMNFFL1 = list()

for(i in commonIndex){
  
  if(edgelist[i,4] == "1" & !(is.na(listAdj2$Apower2[edgelist[i,3],edgelist[i,2]]) |  listAdj2$Apower2[edgelist[i,3],edgelist[i,2]] == 0)){
    MNFFL1[i]=listAdj2$Apower2[edgelist[i,3],edgelist[i,2]]
  } else if(edgelist[i,4] == "1" & !(is.na(listAdj2$Apower3[edgelist[i,3],edgelist[i,2]]) |  listAdj2$Apower3[edgelist[i,3],edgelist[i,2]] == 0)){
    MNFFL1[i]=listAdj2$Apower3[edgelist[i,3],edgelist[i,2]]
  } else if(edgelist[i,4] == "1" & !(is.na(listAdj2$Apower4[edgelist[i,3],edgelist[i,2]]) |  listAdj2$Apower4[edgelist[i,3],edgelist[i,2]] == 0)){
    MNFFL1[i]=listAdj2$Apower4[edgelist[i,3],edgelist[i,2]]
  } else if(edgelist[i,4] == "1" & !(is.na(listAdj2$Apower5[edgelist[i,3],edgelist[i,2]]) |  listAdj2$Apower5[edgelist[i,3],edgelist[i,2]] == 0)){
    MNFFL1[i]=listAdj2$Apower5[edgelist[i,3],edgelist[i,2]]
  } else if(edgelist[i,4] == "1" & !(is.na(listAdj2$Apower6[edgelist[i,3],edgelist[i,2]]) |  listAdj2$Apower6[edgelist[i,3],edgelist[i,2]] == 0)){
    MNFFL1[i]=listAdj2$Apower6[edgelist[i,3],edgelist[i,2]]
  } else if(edgelist[i,4] == "1" & !(is.na(listAdj2$Apower7[edgelist[i,3],edgelist[i,2]]) |  listAdj2$Apower7[edgelist[i,3],edgelist[i,2]] == 0)){
    MNFFL1[i]=listAdj2$Apower7[edgelist[i,3],edgelist[i,2]]
  } else if(edgelist[i,4] == "1" & !(is.na(listAdj2$Apower8[edgelist[i,3],edgelist[i,2]]) |  listAdj2$Apower8[edgelist[i,3],edgelist[i,2]] == 0)){
    MNFFL1[i]=listAdj2$Apower8[edgelist[i,3],edgelist[i,2]]
  } else if(edgelist[i,4] == "1" & !(is.na(listAdj2$Apower9[edgelist[i,3],edgelist[i,2]]) |  listAdj2$Apower9[edgelist[i,3],edgelist[i,2]] == 0)){
    MNFFL1[i]=listAdj2$Apower9[edgelist[i,3],edgelist[i,2]]
  } else if(edgelist[i,4] == "1" & !(is.na(listAdj2$Apower10[edgelist[i,3],edgelist[i,2]]) |  listAdj2$Apower10[edgelist[i,3],edgelist[i,2]] == 0)){
    MNFFL1[i]=listAdj2$Apower10[edgelist[i,3],edgelist[i,2]]
  } else if(edgelist[i,4] == "1" & !(is.na(listAdj2$Apower11[edgelist[i,3],edgelist[i,2]]) |  listAdj2$Apower11[edgelist[i,3],edgelist[i,2]] == 0)){
    MNFFL1[i]=listAdj2$Apower11[edgelist[i,3],edgelist[i,2]]
  } else if(edgelist[i,4] == "1" & !(is.na(listAdj2$Apower12[edgelist[i,3],edgelist[i,2]]) |  listAdj2$Apower12[edgelist[i,3],edgelist[i,2]] == 0)){
    MNFFL1[i]=listAdj2$Apower12[edgelist[i,3],edgelist[i,2]]
  } else if(edgelist[i,4] == "1" & !(is.na(listAdj2$Apower13[edgelist[i,3],edgelist[i,2]]) |  listAdj2$Apower13[edgelist[i,3],edgelist[i,2]] == 0)){
    MNFFL1[i]=listAdj2$Apower13[edgelist[i,3],edgelist[i,2]]
  } else if(edgelist[i,4] == "1" & !(is.na(listAdj2$Apower14[edgelist[i,3],edgelist[i,2]]) |  listAdj2$Apower14[edgelist[i,3],edgelist[i,2]] == 0)){
    MNFFL1[i]=listAdj2$Apower14[edgelist[i,3],edgelist[i,2]]
  } else if(edgelist[i,4] == "1" & !(is.na(listAdj2$Apower15[edgelist[i,3],edgelist[i,2]]) |  listAdj2$Apower15[edgelist[i,3],edgelist[i,2]] == 0)){
    MNFFL1[i]=listAdj2$Apower15[edgelist[i,3],edgelist[i,2]]
  } else if(edgelist[i,4] == "1" & !(is.na(listAdj2$Apower16[edgelist[i,3],edgelist[i,2]]) |  listAdj2$Apower16[edgelist[i,3],edgelist[i,2]] == 0)){
    MNFFL1[i]=listAdj2$Apower16[edgelist[i,3],edgelist[i,2]]
  } else if(edgelist[i,4] == "1" & !(is.na(listAdj2$Apower17[edgelist[i,3],edgelist[i,2]]) |  listAdj2$Apower17[edgelist[i,3],edgelist[i,2]] == 0)){
    MNFFL1[i]=listAdj2$Apower17[edgelist[i,3],edgelist[i,2]]
  }
  
  if( (edgelist[i,4] == "1" & !(is.na(listAdj2$Apower2[edgelist[i,3],edgelist[i,2]]) |  listAdj2$Apower2[edgelist[i,3],edgelist[i,2]] == 0)) | 
      (edgelist[i,4] == "1" & !(is.na(listAdj2$Apower3[edgelist[i,3],edgelist[i,2]]) |  listAdj2$Apower3[edgelist[i,3],edgelist[i,2]] == 0)) | 
      (edgelist[i,4] == "1" & !(is.na(listAdj2$Apower4[edgelist[i,3],edgelist[i,2]]) |  listAdj2$Apower4[edgelist[i,3],edgelist[i,2]] == 0)) |
      (edgelist[i,4] == "1" & !(is.na(listAdj2$Apower5[edgelist[i,3],edgelist[i,2]]) |  listAdj2$Apower5[edgelist[i,3],edgelist[i,2]] == 0)) |
      (edgelist[i,4] == "1" & !(is.na(listAdj2$Apower6[edgelist[i,3],edgelist[i,2]]) |  listAdj2$Apower6[edgelist[i,3],edgelist[i,2]] == 0)) |
      (edgelist[i,4] == "1" & !(is.na(listAdj2$Apower7[edgelist[i,3],edgelist[i,2]]) |  listAdj2$Apower7[edgelist[i,3],edgelist[i,2]] == 0)) | 
      (edgelist[i,4] == "1" & !(is.na(listAdj2$Apower8[edgelist[i,3],edgelist[i,2]]) |  listAdj2$Apower8[edgelist[i,3],edgelist[i,2]] == 0)) | 
      (edgelist[i,4] == "1" & !(is.na(listAdj2$Apower9[edgelist[i,3],edgelist[i,2]]) |  listAdj2$Apower9[edgelist[i,3],edgelist[i,2]] == 0)) |
      (edgelist[i,4] == "1" & !(is.na(listAdj2$Apower10[edgelist[i,3],edgelist[i,2]]) |  listAdj2$Apower10[edgelist[i,3],edgelist[i,2]] == 0)) |
      (edgelist[i,4] == "1" & !(is.na(listAdj2$Apower11[edgelist[i,3],edgelist[i,2]]) |  listAdj2$Apower11[edgelist[i,3],edgelist[i,2]] == 0)) |
      (edgelist[i,4] == "1" & !(is.na(listAdj2$Apower12[edgelist[i,3],edgelist[i,2]]) |  listAdj2$Apower12[edgelist[i,3],edgelist[i,2]] == 0)) | 
      (edgelist[i,4] == "1" & !(is.na(listAdj2$Apower13[edgelist[i,3],edgelist[i,2]]) |  listAdj2$Apower13[edgelist[i,3],edgelist[i,2]] == 0)) | 
      (edgelist[i,4] == "1" & !(is.na(listAdj2$Apower14[edgelist[i,3],edgelist[i,2]]) |  listAdj2$Apower14[edgelist[i,3],edgelist[i,2]] == 0)) |
      (edgelist[i,4] == "1" & !(is.na(listAdj2$Apower15[edgelist[i,3],edgelist[i,2]]) |  listAdj2$Apower15[edgelist[i,3],edgelist[i,2]] == 0)) |
      (edgelist[i,4] == "1" & !(is.na(listAdj2$Apower16[edgelist[i,3],edgelist[i,2]]) |  listAdj2$Apower16[edgelist[i,3],edgelist[i,2]] == 0)) |
      (edgelist[i,4] == "1" & !(is.na(listAdj2$Apower17[edgelist[i,3],edgelist[i,2]]) |  listAdj2$Apower17[edgelist[i,3],edgelist[i,2]] == 0))
  ){idxMNFFL1[i] = i}
  
}

MNFFL1 = unlist(MNFFL1)

idxMNFFL1 = unlist(idxMNFFL1)
```

```
length(idxMNFFL1)
```

```
## [1] 5271
```

\(~\)  
\(~\)  
\(~\)  
\(~\)

Through the following code, the number of edges which are engaged in the multiple-edge subgraghs are computed.

\(~\)

##### MFFL2

```
MFFL2.pval.cor1 = 0
MFFL2.pval.cor.NA = 0
MFFL2.pval.cor2 = 0


for(i in idxMFFL2){
  logic =c()
  if(SignalingNet[[i]]$length > 1){
    
    for(j in 1:SignalingNet[[i]]$length){
      logic[j]= !any(unlist(lapply(SignalingNet[[i]]$corAnalysis[[j]],is.na)))
    }
    if(all(logic)){
      logic1 = c()
      logic2 = c()
      for(j in 1:SignalingNet[[i]]$length){
        
        logic1[j] = SignalingNet[[i]]$corAnalysis[[j]]$pearson$adjusted.pearsonpval < 0.05 & SignalingNet[[i]]$corAnalysis[[j]]$pearson$pearsoncor > 0 
        logic2[j] = SignalingNet[[i]]$corAnalysis[[j]]$pearson$adjusted.pearsonpval < 0.05 & SignalingNet[[i]]$corAnalysis[[j]]$pearson$pearsoncor < 0
      }
      if(all(logic1)) {MFFL2.pval.cor1=MFFL2.pval.cor1+1}
      if(all(logic2)) {MFFL2.pval.cor2=MFFL2.pval.cor2+1}
    }
  } else{
    if(!any(unlist(lapply(SignalingNet[[i]]$corAnalysis,is.na)))){
      a = SignalingNet[[i]]$corAnalysis$pearson$adjusted.pearsonpval < 0.05 & SignalingNet[[i]]$corAnalysis$pearson$pearsoncor > 0 
      b = SignalingNet[[i]]$corAnalysis$pearson$adjusted.pearsonpval < 0.05 & SignalingNet[[i]]$corAnalysis$pearson$pearsoncor < 0
      if(a){MFFL2.pval.cor1=MFFL2.pval.cor1+1}
      if(b){MFFL2.pval.cor2=MFFL2.pval.cor2+1}
    }
  }
}


for(i in idxMFFL2){
  logic = c()
  if(SignalingNet[[i]]$length > 1){
    
    for(j in 1:SignalingNet[[i]]$length){
      
      logic[j]=any(unlist(lapply(SignalingNet[[i]]$corAnalysis[[j]],is.na))) | SignalingNet[[i]]$corAnalysis[[j]]$pearson$adjusted.pearsonpval > 0.05
    }
    if(all(logic)){ MFFL2.pval.cor.NA = MFFL2.pval.cor.NA + 1 }
    
  }else{ if(any(unlist(lapply(SignalingNet[[i]]$corAnalysis,is.na)))| SignalingNet[[i]]$corAnalysis$pearson$adjusted.pearsonpval > 0.05){
    index.pval.cor.NA = MFFL2.pval.cor.NA + 1}
  }
}
```

```
MFFL2.pval.cor1
```

```
## [1] 200
```

```
MFFL2.pval.cor.NA
```

```
## [1] 457
```

```
MFFL2.pval.cor2
```

```
## [1] 106
```

```
length(idxMFFL2) - (MFFL2.pval.cor1 + MFFL2.pval.cor.NA + MFFL2.pval.cor2)
```

```
## [1] 1846
```

\(~\)  
\(~\)

##### MNFBL1

```
MNFBL1.pval.cor1 = 0
MNFBL1.pval.cor.NA = 0
MNFBL1.pval.cor2 = 0


for(i in idxMNFBL1){
  logic =c()
  if(SignalingNet[[i]]$length > 1){
    
    for(j in 1:SignalingNet[[i]]$length){
      logic[j]= !any(unlist(lapply(SignalingNet[[i]]$corAnalysis[[j]],is.na)))
    }
    if(all(logic)){
      logic1 = c()
      logic2 = c()
      for(j in 1:SignalingNet[[i]]$length){
        
        logic1[j] = SignalingNet[[i]]$corAnalysis[[j]]$pearson$adjusted.pearsonpval < 0.05 & SignalingNet[[i]]$corAnalysis[[j]]$pearson$pearsoncor > 0 
        logic2[j] = SignalingNet[[i]]$corAnalysis[[j]]$pearson$adjusted.pearsonpval < 0.05 & SignalingNet[[i]]$corAnalysis[[j]]$pearson$pearsoncor < 0
      }
      if(all(logic1)) {MNFBL1.pval.cor1=MNFBL1.pval.cor1+1}
      if(all(logic2)) {MNFBL1.pval.cor2=MNFBL1.pval.cor2+1}
    }
  } else{
    if(!any(unlist(lapply(SignalingNet[[i]]$corAnalysis,is.na)))){
      a = SignalingNet[[i]]$corAnalysis$pearson$adjusted.pearsonpval < 0.05 & SignalingNet[[i]]$corAnalysis$pearson$pearsoncor > 0 
      b = SignalingNet[[i]]$corAnalysis$pearson$adjusted.pearsonpval < 0.05 & SignalingNet[[i]]$corAnalysis$pearson$pearsoncor < 0
      if(a){MNFBL1.pval.cor1=MNFBL1.pval.cor1+1}
      if(b){MNFBL1.pval.cor2=MNFBL1.pval.cor2+1}
    }
  }
}


for(i in idxMNFBL1){
  logic = c()
  if(SignalingNet[[i]]$length > 1){
    
    for(j in 1:SignalingNet[[i]]$length){
      
      logic[j]=any(unlist(lapply(SignalingNet[[i]]$corAnalysis[[j]],is.na))) | SignalingNet[[i]]$corAnalysis[[j]]$pearson$adjusted.pearsonpval > 0.05
    }
    if(all(logic)){ MNFBL1.pval.cor.NA = MNFBL1.pval.cor.NA + 1 }
    
  }else{ if(any(unlist(lapply(SignalingNet[[i]]$corAnalysis,is.na)))| SignalingNet[[i]]$corAnalysis$pearson$adjusted.pearsonpval > 0.05){
    index.pval.cor.NA = MNFBL1.pval.cor.NA + 1}
  }
}
```

```
MNFBL1.pval.cor1
```

```
## [1] 773
```

```
MNFBL1.pval.cor.NA
```

```
## [1] 1851
```

```
MNFBL1.pval.cor2
```

```
## [1] 423
```

```
length(idxMNFBL1) - (MNFBL1.pval.cor1 + MNFBL1.pval.cor.NA + MNFBL1.pval.cor2)
```

```
## [1] 8496
```

\(~\)  
\(~\)

##### MNFFL1

```
MNFFL1.pval.cor1 = 0
MNFFL1.pval.cor.NA = 0
MNFFL1.pval.cor2 = 0


for(i in idxMNFFL1){
  logic =c()
  if(SignalingNet[[i]]$length > 1){
    
    for(j in 1:SignalingNet[[i]]$length){
      logic[j]= !any(unlist(lapply(SignalingNet[[i]]$corAnalysis[[j]],is.na)))
    }
    if(all(logic)){
      logic1 = c()
      logic2 = c()
      for(j in 1:SignalingNet[[i]]$length){
        
        logic1[j] = SignalingNet[[i]]$corAnalysis[[j]]$pearson$adjusted.pearsonpval < 0.05 & SignalingNet[[i]]$corAnalysis[[j]]$pearson$pearsoncor > 0 
        logic2[j] = SignalingNet[[i]]$corAnalysis[[j]]$pearson$adjusted.pearsonpval < 0.05 & SignalingNet[[i]]$corAnalysis[[j]]$pearson$pearsoncor < 0
      }
      if(all(logic1)) {MNFFL1.pval.cor1=MNFFL1.pval.cor1+1}
      if(all(logic2)) {MNFFL1.pval.cor2=MNFFL1.pval.cor2+1}
    }
  } else{
    if(!any(unlist(lapply(SignalingNet[[i]]$corAnalysis,is.na)))){
      a = SignalingNet[[i]]$corAnalysis$pearson$adjusted.pearsonpval < 0.05 & SignalingNet[[i]]$corAnalysis$pearson$pearsoncor > 0 
      b = SignalingNet[[i]]$corAnalysis$pearson$adjusted.pearsonpval < 0.05 & SignalingNet[[i]]$corAnalysis$pearson$pearsoncor < 0
      if(a){MNFFL1.pval.cor1=MNFFL1.pval.cor1+1}
      if(b){MNFFL1.pval.cor2=MNFFL1.pval.cor2+1}
    }
  }
}


for(i in idxMNFFL1){
  logic = c()
  if(SignalingNet[[i]]$length > 1){
    
    for(j in 1:SignalingNet[[i]]$length){
      
      logic[j]=any(unlist(lapply(SignalingNet[[i]]$corAnalysis[[j]],is.na))) | SignalingNet[[i]]$corAnalysis[[j]]$pearson$adjusted.pearsonpval > 0.05
    }
    if(all(logic)){ MNFFL1.pval.cor.NA = MNFFL1.pval.cor.NA + 1 }
    
  }else{ if(any(unlist(lapply(SignalingNet[[i]]$corAnalysis,is.na)))| SignalingNet[[i]]$corAnalysis$pearson$adjusted.pearsonpval > 0.05){
    index.pval.cor.NA = MNFFL1.pval.cor.NA + 1}
  }
}
```

```
MNFFL1.pval.cor1
```

```
## [1] 393
```

```
MNFFL1.pval.cor.NA
```

```
## [1] 801
```

```
MNFFL1.pval.cor2
```

```
## [1] 193
```

```
length(idxMNFFL1) - (MNFFL1.pval.cor1 + MNFFL1.pval.cor.NA + MNFFL1.pval.cor2)
```

```
## [1] 3884
```

\(~\)  
\(~\)

##### MPFBL2

```
MPFBL2.pval.cor1 = 0
MPFBL2.pval.cor.NA = 0
MPFBL2.pval.cor2 = 0


for(i in idxMPFBL2){
  logic =c()
  if(SignalingNet[[i]]$length > 1){
    
    for(j in 1:SignalingNet[[i]]$length){
      logic[j]= !any(unlist(lapply(SignalingNet[[i]]$corAnalysis[[j]],is.na)))
    }
    if(all(logic)){
      logic1 = c()
      logic2 = c()
      for(j in 1:SignalingNet[[i]]$length){
        
        logic1[j] = SignalingNet[[i]]$corAnalysis[[j]]$pearson$adjusted.pearsonpval < 0.05 & SignalingNet[[i]]$corAnalysis[[j]]$pearson$pearsoncor > 0 
        logic2[j] = SignalingNet[[i]]$corAnalysis[[j]]$pearson$adjusted.pearsonpval < 0.05 & SignalingNet[[i]]$corAnalysis[[j]]$pearson$pearsoncor < 0
      }
      if(all(logic1)) {MPFBL2.pval.cor1=MPFBL2.pval.cor1+1}
      if(all(logic2)) {MPFBL2.pval.cor2=MPFBL2.pval.cor2+1}
    }
  } else{
    if(!any(unlist(lapply(SignalingNet[[i]]$corAnalysis,is.na)))){
      a = SignalingNet[[i]]$corAnalysis$pearson$adjusted.pearsonpval < 0.05 & SignalingNet[[i]]$corAnalysis$pearson$pearsoncor > 0 
      b = SignalingNet[[i]]$corAnalysis$pearson$adjusted.pearsonpval < 0.05 & SignalingNet[[i]]$corAnalysis$pearson$pearsoncor < 0
      if(a){MPFBL2.pval.cor1=MPFBL2.pval.cor1+1}
      if(b){MPFBL2.pval.cor2=MPFBL2.pval.cor2+1}
    }
  }
}


for(i in idxMPFBL2){
  logic = c()
  if(SignalingNet[[i]]$length > 1){
    
    for(j in 1:SignalingNet[[i]]$length){
      
      logic[j]=any(unlist(lapply(SignalingNet[[i]]$corAnalysis[[j]],is.na))) | SignalingNet[[i]]$corAnalysis[[j]]$pearson$adjusted.pearsonpval > 0.05
    }
    if(all(logic)){ MPFBL2.pval.cor.NA = MPFBL2.pval.cor.NA + 1 }
    
  }else{ if(any(unlist(lapply(SignalingNet[[i]]$corAnalysis,is.na)))| SignalingNet[[i]]$corAnalysis$pearson$adjusted.pearsonpval > 0.05){
    index.pval.cor.NA = MPFBL2.pval.cor.NA + 1}
  }
}
```

```
MPFBL2.pval.cor1
```

```
## [1] 45
```

```
MPFBL2.pval.cor.NA
```

```
## [1] 160
```

```
MPFBL2.pval.cor2
```

```
## [1] 34
```

```
length(idxMPFBL2) - (MPFBL2.pval.cor1 + MPFBL2.pval.cor.NA + MPFBL2.pval.cor2)
```

```
## [1] 735
```

\(~\)

\(~\)

\(~\)

\(~\)

\(~\)

\(~\)

\(~\)

\(~\)

Table1

\(~\)

\(~\)

### Results

\(~\)  
\(~\)

Table2

\(~\)

Table3

\(~\)

\(~\)

\(~\)

\(~\)

\(~\)
